# An mTOR-independent Macroautophagy Activator Ameliorates Tauopathy and Prionopathy Neurodegeneration Phenotypes

**DOI:** 10.1101/2022.09.29.509997

**Authors:** Leonard Yoon, Rachel C. Botham, Adriaan Verhelle, Yin Wu, Christian M. Cole, Ee Phie Tan, Lynée A. Massey, Pablo Sanz-Martinez, Ching-Chieh Chou, Jin Xu, Lisa P. Elia, Kyunga Lee, Sergio R. Labra, Gabriel M. Kline, Qiang Xiao, Derek Rhoades, Sara Cano-Franco, Caroline A. Cuoco, Amina Ta, Wen Ren, William C. Hou, Cristian Wulkop-Gil, Lara H. Ibrahim, Kayla Nutsch, Yeonjin Ko, Oren L. Lederberg, Hongfan Peng, Starr Jiang, Stuart A. Lipton, Michael J. Bollong, Malene Hansen, Steven Finkbeiner, Daniel Finley, Miguel A. Prado, H. Michael Petrassi, Ivan Dikic, Fulvio Reggiori, Danny Garza, R. Luke Wiseman, Evan T. Powers, Judith Frydman, Stephen J. Haggarty, Kristen A. Johnson, M. Catarina Silva, Alexandra Stolz, Sandra E. Encalada, Jeffery W. Kelly

## Abstract

Autophagy-lysosomal impairment is an early and prominent feature of neurodegeneration. Autophagy activation reduces protein aggregates and lipid level abnormalities. We performed a high-content imaging-based screen assessing 940,000 small molecules to identify those that reduce lipid droplet numbers. Of 77 validated, structurally diverse hits, 24 increased autophagy flux reporter activity, consistent with accelerated lipid droplet clearance by lipophagy. Of these, we show that CCT020312 activates autophagy independently of mammalian target of rapamycin (mTOR) inhibition, to avoid immunosuppression. CCT020312 reduced insoluble phosphorylated tau levels and tau-mediated neuronal stress vulnerability, as well as reducing intracellular Aβ levels within directly induced neurons bearing epigenetic marks of aging derived from Alzheimer’s patient fibroblasts. Moreover, CCT020312 cleared mutant prion protein aggregates and normalized trafficking deficiencies in axons of a cellular model of familial prion disease. Autophagy is widely considered a promising strategy to attenuate neurodegeneration, and here we introduce a strategy to discover new pharmacology.

## Introduction

Autophagy is an intracellular lysosome-mediated degradative pathway critical for maintaining cellular homeostasis and resilience in response to diverse stresses.^1–3^ Autophagy turns over potentially cytotoxic and superfluous substrates, including pathogens, dysfunctional organelles,^4–10^ aggregated proteins, lipids,^11,12^ and macromolecular assemblies of biomolecules.^13–16^ Macroautophagy is one type of autophagy that initiates with nucleation of the phagophore, which gathers the substrates targeted for degradation and closes to form an autophagosome delimited by a double membrane (Figure 1A)^1,3^. Subsequent fusion of the autophagosome with late endosomes and/or lysosomes yields the autolysosome, wherein the inner lipid bilayer derived from the autophagosome and substrates are degraded by lysosomal hydrolases (Figure 1A)^1,3^.

**Figure 1:**
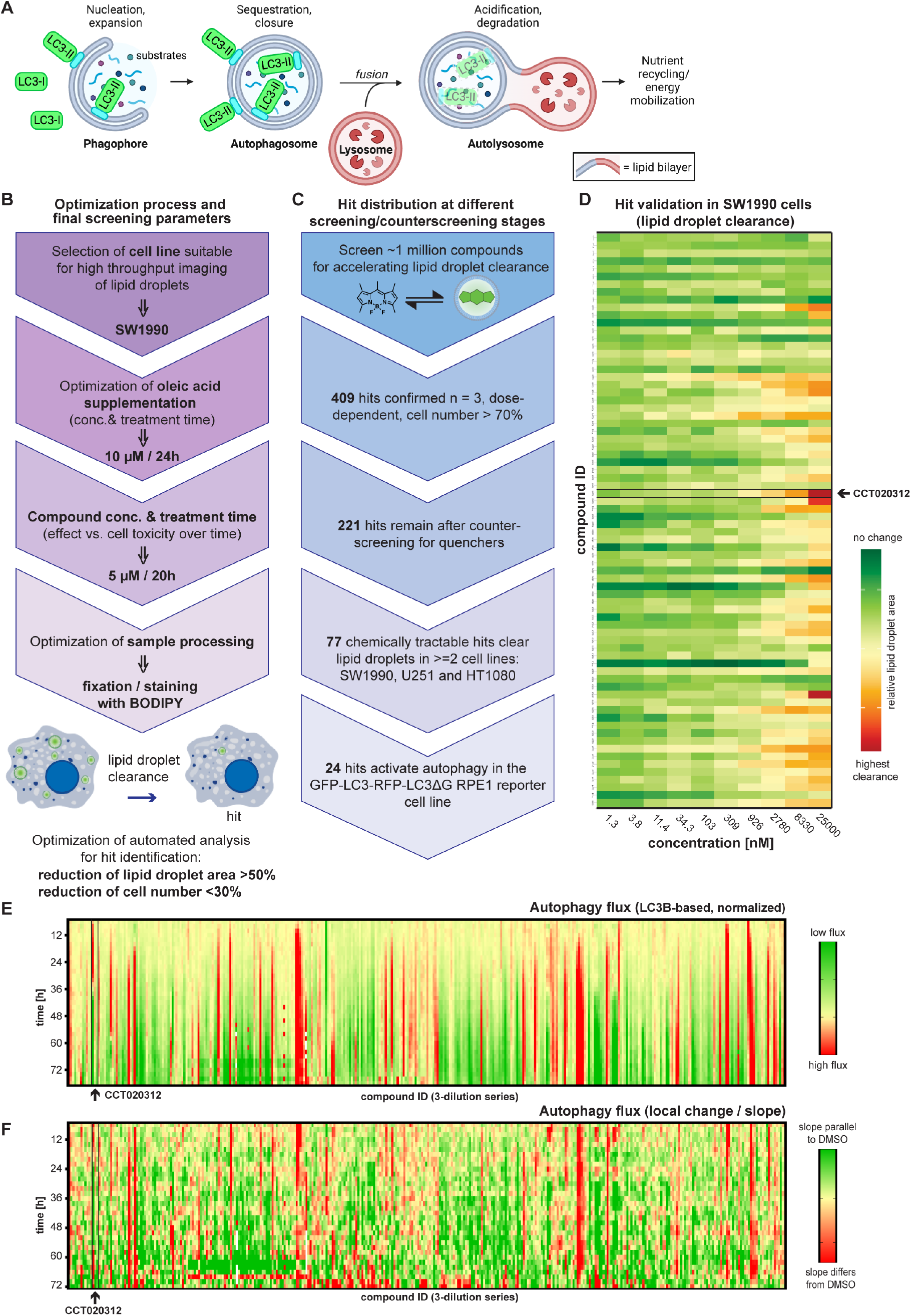
Lipid droplet degradation screen identifies novel autophagy activators. (**A**) Schematic representation of macroautophagy. Cytosolic cargo is sequestered in an LC3-dependent manner into *de novo*-synthesized, double-membrane-bound autophagosomes that subsequently fuse with lysosomes to degrade their contents. (**B**) Schematic representation of the process to set up and optimize a cell-based, high-throughput screen for the identification of small molecules (compounds) accelerating lipid droplet clearance. Parameters chosen for the final screen setting are indicated in bold. The screen uses a fluorescent probe, BODIPY 493/503, that accumulates in hydrophobic environments like lipid droplets. Cells treated with vehicle control display higher number and area of fluorescent punctae relative to cells treated with autophagy activator compounds. (**C**) Schematic representation of the screening pipeline employed to identify and prioritize compounds that clear lipid droplets (which become fluorescent upon BODIPY 493/503 staining) out of a 1 million compound library. The number of hits remaining at each step of the validation process is indicated in bold. The pipeline includes a primary screen to identify compounds that clear lipid droplets dose-dependently in a non-cytotoxic manner, the removal of compounds that are known fluorescent artifacts, quenchers, or PAINS, the validation of lipid droplet clearance in multiple immortalized cell lines, and a final autophagy flux screen. (**D**) Heatmap of the number of lipid droplets per cell (normalized to DMSO) in SW1990 cells pretreated for 24 h with 10µM oleic acid followed by treatment for 20 h with 77 hit compounds at different concentrations (range: 1.3nM-25µM). The heat map tiles of CCT020312 are surrounded by a black rectangle identified by an arrow. (**E,F**) Autophagy flux screen of 120 compounds in RPE1 cells using the reporter GFP-LC3-RFP-LC3ΔG. (**E**) Heatmap representing the difference between the average (n = 3) of GFP/RFP ratio of total integrated fluorescence intensities of individual compounds at three different concentrations (0.05µM, 0.5µM, and 5µM) to the average (n = 18) GFP/RFP ratio of DMSO. (**F**) Heatmap representing the difference between the slope of individual compounds and DMSO (n = 3): the slope of each subsequent set of two timepoints was averaged and compared to respective DMSO data. Red represents higher autophagy flux and green lower autophagy flux induction than DMSO. Thresholds of heatmaps were adjusted to fully represent the whole set of data. The heat map tiles of CCT020312 are surrounded by a black rectangle identified by an arrow.

Autophagy occurs at a basal level and can be induced in response to numerous stimuli, including stress, nutrient starvation, or inhibition of mammalian target of rapamycin mTOR complex 1 (mTORC1), the latter a central hub in the regulation of cell metabolism and other cellular/organismal processes.^17–19^ Prior cell-based screening efforts to identify small molecule autophagy activators have largely been based on the increased autophagosomal localization of a fluorescently-labeled autophagy effector protein called microtubule-associated protein 1A/1B light chain 3 or LC3.^20–23^ LC3 is expressed cytosolically and is constitutively cleaved upon translation into LC3-I; upon macroautophagy initiation, it is lipidated to LC3-II, which localizes to the inner and outer membranes of the autophagosome and binds different protein adaptors, including selective autophagy receptors on the inner membrane (Figure 1A).^24^ The LC3-II pool localized in the interior of autophagosomes becomes degraded by lysosomal proteases in autolysosomes; these dynamic changes can be visualized by either immunoblotting (LC3-I and LC3-II are separable) or imaging fluorescently labeled LC3 proteins, which transition from a cytosolic distribution to autophagosome-localized puncta.^25^

Autophagy dysfunction is associated with a variety of central nervous system (CNS) neurodegenerative diseases, including Alzheimer’s (AD), Parkinson’s, Huntington’s, prion disease, and amyotrophic lateral sclerosis.^10,13,26–57^ Naturally occurring mutations in protein effectors of autophagy have revealed a crucial role for autophagy in preventing neurodegeneration upon aging; findings that have been robustly validated in murine models.^10,13,26–48,50–58^ During aging, the efficiency of autophagy can decline for genetic and/or environmental reasons, leading to the detrimental accumulation of protein aggregates, lipids and damaged organelles, thereby predisposing an individual to exhibit post-mitotic tissue degeneration.^59^ Current evidence supports the notion that autophagy plays a neuroprotective role in post-mitotic cells by maintaining cellular homeostasis through organelle turnover and the continuous removal of aggregated proteins.^60^ Thus, pharmacological activation of autophagy represents a promising strategy for mitigating CNS diseases.^19,26,61–65^ Autophagy activation may also be able to attenuate inflammatory processes linked to neurodegeneration.^66–69^

Numerous efforts have been undertaken to identify small-molecule autophagy activators.^20–23,70–72^ Pharmacological activation of autophagy has been historically achieved by treatment with the natural product rapamycin, an inhibitor of mTORC1, as well as more potent second-generation mTORC1 inhibitors such as Torin-1 or AZD8055.^22,64,70,73,74^ In addition to being a master regulator of autophagy, mTORC1 influences immune function;^75–77^ with mTORC1 inhibition mediating immunosuppression.^78^ Therefore, we sought mTORC1 inhibition–independent autophagy activators to complement the progress being made in generating potent and selective mTORC1 inhibitors, which are under clinical evaluation.^79–84^ Though a handful of mTORC1-independent autophagy activators have been reported in the literature,^83–85^ there are likely additional mTORC1-independent mechanisms of autophagy activation that have yet to be pharmacologically modulated based on a genome-wide siRNA screen.^21^ In addition, the combined induction of mTORC1-dependent and mTORC1-independent pathways of autophagy activation improved AD-like pathology in a mouse model, highlighting the potential for mTORC1-independent autophagy activation to ameliorate ND phenotypes.^86^

Oleic acid (OA)-based triglycerides accumulate in AD.^87^ Moreover exposing cells to fatty acids such as OA results in the fatty acids being converted to triacylglycerides within the endoplasmic reticulum (ER) bilayer, resulting in lipid droplets (LDs) enclosed within a single phospholipid monolayer.^88,89^ LDs are dynamic, ubiquitous intracellular organelles that serve as critical hubs regulating cellular lipid metabolism.^90–92^ Prior studies in hepatocytes show that pharmacological inhibition of autophagy increases triacylglycerol content almost twofold, whereas pharmacological activation of autophagy by the mTORC1 inhibitor rapamycin decreases LD count and levels of triacylglycerols.^12^ Therefore it is established that LDs are known autophagy substrates. We thus hypothesized that we could shift the focus away from monitoring protein effectors of autophagy toward quantifying the successful degradation of LD substrate to uncover new autophagy activators, as not all autophagy pathways are LC3-mediated. Literature precedent exists for quantifying BODIPY-stained LDs via fluorescence microscopy, flow cytometry and in a plate reader.^93–96^ Lipolysis also breaks down stored lipids via LD– associated cytosolic lipases, such as the adipose triglyceride lipase.^97,98^ The fatty acids resulting from both degradation pathways can be utilized by mitochondria and peroxisomes for energy production through β-oxidation.^90,99^ Thus, the LD screen was expected to identify small molecules that reduce LD numbers by both autophagic and non-autophagic pathways.

The potential for discovering autophagy activators by monitoring endogenous LD clearance was first demonstrated in a HeLa cell–based high-content imaging screen of 1400 small molecules. This screen identified one mTORC1-independent autophagy activator that is structurally similar to tamoxifen, P29A03.^82^ Inspired by this previous screen, we envisioned treating cells with a fatty acid to increase the number of LDs in order to increase the sensitivity of the screen. We describe the optimization, miniaturization, and execution of a high-content imaging-based high-throughput screen (HTS) for identifying small molecules that reduce cellular LD numbers. Validated hits of interest were further scrutinized in multiple follow-up autophagy flux assays in multiple cell types, including in neurons. We highlight one of our hit compounds, CCT020312, for its potential value in restoring neuronal function and viability in multiple cellular models of neurodegeneration. CCT020312 treatment of primary murine hippocampal neurons at concentrations above 50nM reduced cytotoxic axonal endosome-delimited aggregates of the mutant prion protein (PrP^PG14^) and restored axonal vesicular trafficking. In addition, CCT020312 robustly reduced phosphorylated and insoluble tau in a human induced pluripotent stem cell (iPSC)-derived neuronal hereditary tauopathy model, while also reducing tau-mediated neuronal stress vulnerability. Moreover, CCT020312 reduced Aβ levels in directly induced neurons harboring the epigenetic marks of aging derived from fibroblasts from a patient with sporadic AD and from a hereditary AD patient carrying the presenilin 1 (PSEN1) A431E mutation. Together, these data demonstrate the potential of CCT020312 to be a medicinal chemistry starting point for improving potency, solubility and blood-brain barrier permeability to slow neurodegenerative disease progression in humans. Other mTORC1-independent hit compounds from this screen are being optimized and will be published separately as autophagy activators that slow neurodegenerative disease progression.

## Results

### Lipid Droplet clearance as a basis for discovering autophagy flux activators

We set out to discover novel pharmacological activators of autophagy by first optimizing the amount of OA to feed to human SW1990 adenocarcinoma cells, chosen for their large size, uniform growth patterns, and adherent properties that facilitate image-based analysis of cellular LD load. SW1990 cells were plated in a standard complete growth medium supplemented with 10µM OA complexed to bovine serum albumin (BSA) and allowed to adhere for approximately 24 h to elevate their cellular LD load, envisioned to render this screen more sensitive (Figure 1B). Notably, this screen was carried out in rich media to discover compounds able to activate autophagy by mechanisms not involving nutrient starvation (which is the most studied mode of non-pharmacological autophagy activation in the literature).^18^ A prior manuscript reported that 50µM OA induced autophagy in tongue squamous cell carcinomas;^100^ however, we found that supplementing the SW1990 cell culture medium with up to 100µM OA to increase cellular LD density did not activate autophagy by itself, as evidenced by the lack of increase in LC3-II immunoblot band intensity (Figure S1A). In LC3-II immunoblot analysis, both activation (e.g. via rapamycin or AZD8055) and inhibition (e.g. via chloroquine) of autophagy increase the cellular abundance of LC3-II, either by inducing the aforementioned maturation of LC3-I into LC3-II or by inhibiting LC3-II degradation.^101^ To distinguish between these two scenarios, we co-treated with chloroquine (CQ) for the final 4 h of treatment to inhibit LC3-II degradation, and visualized expected results from positive control compounds.^102^ Up to 100µM OA, meanwhile, did not modulate autophagy flux in SW1990 cells (Figure S1A).

For compound treatment time optimization, cells were initially treated with 50µM OA for 24 h, at which time two autophagy activators—rapamycin and G protein-coupled estrogen receptor agonist tamoxifen—were added individually without a media change for 8 h, 16 h or 24 h (Figure S1B). Following compound treatment, SW1990 cells were fixed, stained with BODIPY 493/503, a dye that integrates into LDs^28,82,103,104^, and imaged. We detected significant decrease of LD number in SW1990 cells with rapamycin and tamoxifen after 16 h and 24 h (Figure S1B). After this preliminary experiment, we wanted to see if we could improve the signal-to-noise by dropping OA supplementation to 10µM for 24 h. Additional autophagy activators were employed to scrutinize this LD change, including PI-103, a class 1A phosphatidylinositide 3 kinase/mTORC1/mTORC2 inhibitor, and PP242, a mTORC1/mTORC2 dual inhibitor. We correlated LD decrease with a classical autophagy effector, the conversion of cytosolic LC3-I into lipidated LC3-II via Western blot (Figure S1C).^101^ LD decrease and autophagy induction based on LC3-II levels were found to be dose-dependently correlated for the three autophagy activators tested (Figure S1C), indicating that our LD assay was suitable for identifying autophagy inducers employing a 24-h pretreatment with 10µM OA media supplementation followed by 20-h treatment with a candidate autophagy activator. We also observed a substantial decrease in cellular LD area in HeLa cells, U251 cells and HT-1080 cells (cells commonly used in research), suggesting the generalizability of this approach (Figure S1D).

To interrogate screening compound dose and whether the screen could be performed in a high-throughput 1536-well format, we assessed a collection of 1280 annotated bioactive small molecules, the Library of Pharmacologically Active Compounds (LOPAC, see Materials and Methods). Using the selections outlined in Figure 1B, coupled with the development and application of a semi-automated pipeline for analysis of cellular LD images, we performed the screen three times with 59% of the LOPAC hits being confirmed in at least 2 of the 3 1536-well screening plates. Since many of the top hits reducing LD number were modulators of adrenergic signaling, known to activate lipolysis in adipocytes,^105^ these compounds validated the HTS strategy.

### Million-compound phenotypic high-throughput screen for compounds reducing LD number

With the LD assay optimized for screening in singlet at a 5µM candidate small molecule concentration, we employed a diverse 940,000-member small molecule library built over 20 years from smaller compound sets (e.g. Enamine, ChemDiv, Life Chemicals) at Calibr (a drug discovery division of Scripps Research). A total of 7105 primary hit compounds decreased LD count using single-factor normalization to select hits that had a significant difference from the median compound signal on the plates (RBZ’ score). These primary hits were reassessed in triplicate in a dose response, with 729 hits being confirmed. Hits were further assessed for noncytotoxic dose-dependent reduction (concentration range: 0.1 to 10µM) in cellular lipid load. Of these, 409 achieved a >50% reduction in total lipid area while exhibiting <30% reduction in cell number at the maximum nontoxic dose.

Next, a counter screen was performed on these 409 compounds to identify and remove those hits that caused significant quenching of BODIPY 493/503 fluorescence in fixed cells, where biological processes like macroautophagy to clear LDs cannot occur (see Materials and Methods for more details). This approach eliminated 188 false positive BODIPY quenchers. The remaining 221 hit compounds (Figure 1C) were manually curated to eliminate pan-assay interference compounds,^106^ compounds bearing significant metabolic liabilities, compounds that were often hits in multiple unrelated high-throughput screens, and compounds displaying a high degree of structural similarity. This effort yielded 108 compounds, of which 81 were commercially available.

The 81 reordered compounds were analyzed for their ability to reduce LD number in three different cancer cell lines: the original screening SW1990 adenocarcinoma cell line, an HT-1080 fibrosarcoma cell line, and a U251 glioblastoma cell line. 77 compounds decreased LD count in at least two out of the three cell lines (Figure S1E), suggesting that these compounds function through a conserved cellular pathway. The concentration-dependent LD reduction data from the 77 reordered hit compounds in SW1990 cells is shown in Figure 1D.

### A subset of LD-reducing compounds activates autophagy flux

To identify those hit compounds that accelerate LD clearance through macroautophagy activation, we made use of the previously published autophagy flux reporter GFP-LC3-RFP-LC3ΔG in the immortalized human retinal pigment epithelial RPE1 cells to complement our LD screen.^20^ This LC3B-based, ratiometric reporter is compatible with live cell analysis. We anticipated that mechanistically diverse autophagy activators would yield distinct GFP-LC3/RFP-LC3ΔG kinetic plots over 72h time courses. Unaltered GFP-LC3/RFP-LC3ΔG kinetic plots relative to vehicle control would also allow us to discover compounds that reduced LD load by autophagy-independent pathways; in fact, two known diacylglycerol O-acyltransferase 1 (DGAT) inhibitor hits blocking LD biosynthesis were identified.

Out of the 221 compounds remaining after counter-screening for quenchers, including the 77 compounds validated to dose-dependently enhance LD clearance in at least 2 out of 3 cell lines, 120 were tested in triplicate at three different concentrations (50nM, 500nM, and 5µM) in the GFP-LC3-RFP-LC3ΔG RPE1 reporter cell line (Figure 1E).^20^ Images in the green, red, and phase channels were taken every 2 h over a 72h period and analyzed for total integrated intensity (green, red) and cell confluency (phase). Autophagy modulator hits were defined as compounds causing a ≥20% increase in autophagy flux induction over a ≥6h treatment period relative to the activation obtained following treatment with the established macroautophagy activator Torin-1 (cf. CCT020312 to Torin-1 in Figure S1F; also see Materials and Methods).

40 of 120 hits tested were identified as primary macroautophagy hits, of which 24 were found to be in common with the subset of 77 validated LD clearance hits. 35 out of 40 primary autophagy hits were retested in a dose-dependent manner using 12 dilutions (50nM to 10µM) Multiple distinct GFP-LC3/RFP-LC3ΔG 72h time courses were observed for different hits, consistent with the hypothesis that hit compounds activated autophagy via different mechanisms of activation (Figure S1F, G).

### Identification of novel small-molecule autophagy activator CCT020312

We prioritized hit compound CCT020312 (structure in Figure 2A) for further analysis due to (i) top-quartile (low single-digit micromolar) activity in the LD assay in all three cancer cell lines (Figure 2B, 2C, S1E) while not causing toxicity in CellTiter-Glo luminescent cell viability assay in SW1990 and U251 cells at doses below 10µM (Figure S2A), (ii) prior literature evidence of CCT020312 activating PERK, which has been linked to macroautophagy activation^107,108^ and (iii) induction of macroautophagy flux in RPE1 cells stably expressing the GFP-LC3-RFP-LC3ΔG reporter at concentrations between 600nM-4µM (Figure 2D, 2E, S1F, S1G). Cell toxicity with this immortalized cell line was noticeable at concentrations ≥ 3µM (Figure S2B). We independently synthesized CCT020312 to produce enough material for all downstream biological assays, to assess the compound for stability in PBS at room temperature for up to 12 days and characterize its proclivity for self-assembly or colloid formation (Data S1).^109^ We found that this independently synthesized CCT020312 was also a macroautophagy activator. A classical lipidation assay of LC3B in RPE1 cells^101^ indicated that CCT020312 induced lipidation of LC3B (LC3-II) in a dose-dependent manner (Figure 2F). LC3-II band intensity is further increased upon BafA1 cotreatment with CCT020312, which is consistent with the expected phenotype of a macroautophagy activator.^102^ HeLa cells treated for 20 h with CCT020312 formed more LC3B puncta than vehicle-treated control cells, and these puncta colocalized with lysosome-associated membrane protein 2 (LAMP2), suggesting CCT020312 increased autolysosome number (Figure S2C, S2D), which is consistent with our observed result in RPE1 cells stably expressing the GFP-LC3-RFP-LC3ΔG reporter demonstrating that autophagy flux is increased.

**Figure 2:**
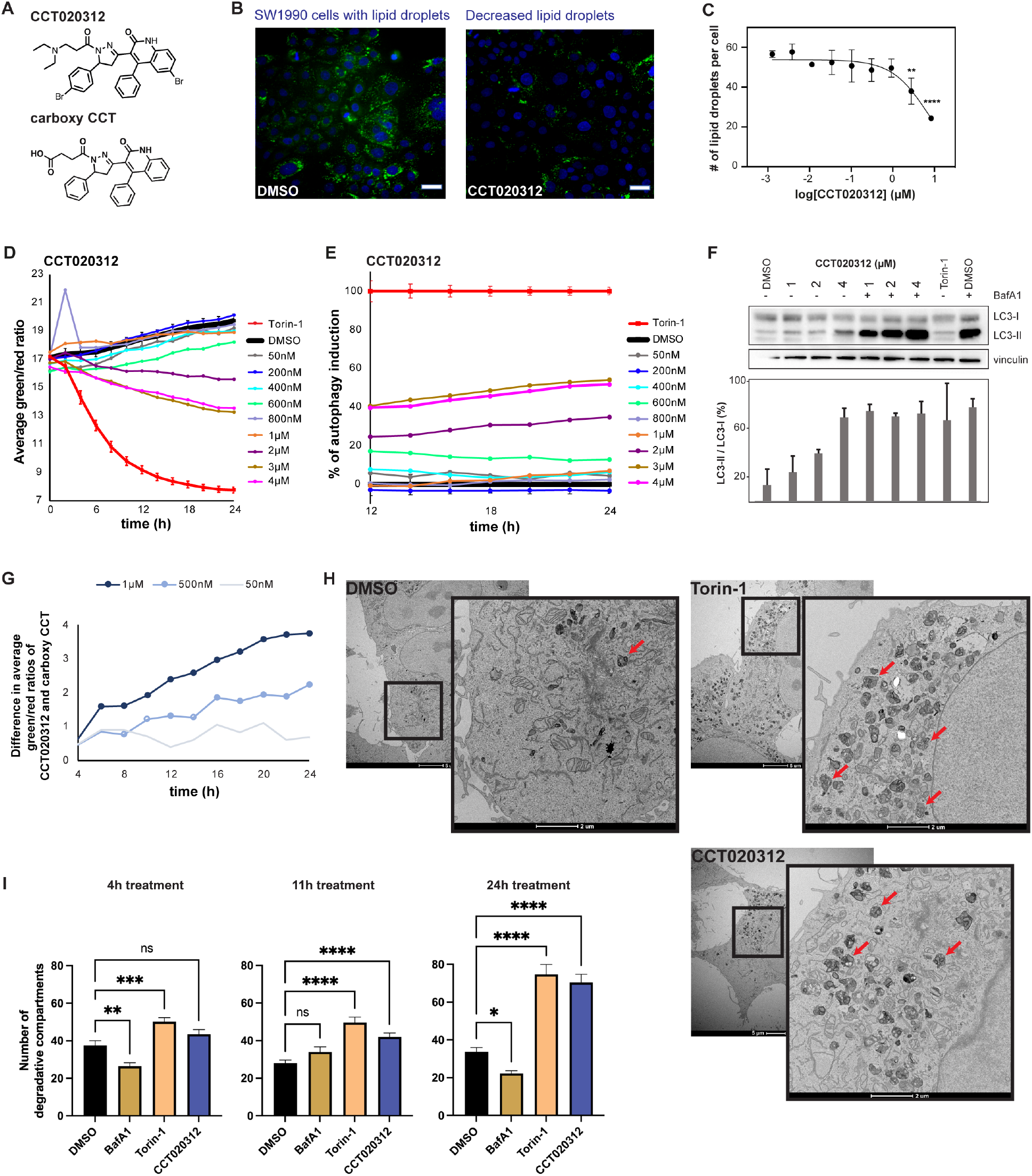
CCT020312 induces autophagic flux. (**A**) Chemical structures of CCT020312 and inactive analog carboxy CCT. (**B**) Representative images of SW1990 cells fed with 10µM oleic acid for 24 hours followed by 20-hour treatment with 0.1% DMSO or 2.7µM CCT020312. Lipid droplets are stained with BODIPY 493/503 fluorescence (green); nuclei are stained with Hoechst 33342 (blue), respectively. Scale bar = 100 µm. (**C**) CCT020312 dose-response curve from SW1990 cells fed with 10µM oleic acid followed by 18-hour treatment. n = 18. Points and error bars depict mean ± SEM ** Holm multiple comparisons test following one-way ANOVA p < .01, **** p < .0001. (**D**) Plot of average red/green fluorescence ratio over time of RPE1 cells stably expressing the reporter GFP-LC3-mCherry-LC3ΔG and treated with different concentrations of CCT020312, DMSO, or 250nM Torin-1. (**E**) Normalized data shown in Figure 2D. Percent of autophagy flux over time in RPE1 cells stably expressing the reporter GFP-LC3-mCherry-LC3ΔG and treated with different concentrations of CCT020312. (**F**) Example western blot analysis of RPE1 cells treated with indicated concentrations of compounds over 4.5 hours (upper panel) with corresponding LC3-II / LC3-I ratio quantification (lower panel, n = 3, bars and error bars depict mean + SEM). BafA1 = 0.2 µg/mL Bafilomycin A1. (**G**) Plot depicting the difference between the green/red ratios of total integrated fluorescence intensities of CCT020312 and carboxy CCT (inactive analog) over time. Higher numbers represent higher autophagy flux upon CCT020312 treatment in comparison to carboxy CCT treatment at the same concentration. Statistical significance for each time point has been calculated by one-sided Student’s T-distribution and indicated as a circle if significant; n = 8; no circle: not significant, empty circle: p < 0.05, filled circle: p < 0.0005. (**H**) Representative transmission electron microscopy images of HeLa cells treated for 4, 11 or 24 h with vehicle (0.1% DMSO), 250nM Torin-1, 100nM Bafilomycin A1 (BafA1) or 2.5µM CCT020312. Only the vehicle, Torin-1 and CCT020312 sample at the 24 h time point are shown. High magnification image depicts the boxed area in the low magnification image. Scale bar of low magnification images, 5 µm; scale bar of high magnification images, 2 µm. Some DGCs are highlighted with a red arrow. (**I**) Quantification of the DGCs per cell section in the experiment shown in Panel A. n = 42-100 HeLa cells per treatment condition, coming from N = 2 biological replicates of cells. Bars and error bars depict mean + SEM. * Dunn’s multiple comparisons test (to DMSO for a given treatment time) following Kruskal-Wallis * p < .05, ** p < .01, *** p < .001, **** p < .0001.

To further differentiate between targeted macroautophagy induction versus a response caused by compound toxicity,^22^ we compared the macroautophagy flux induction caused by CCT020312 to a structurally similar yet inactive compound, carboxy CCT (Figure 2A). The difference in autophagy flux induction between active and inactive compound using the ratiometric GFP-LC3-RFP-LC3ΔG time course was apparent at concentrations ≥ 500nM, with increasing significance at higher doses (Figure 2G).

### CCT020312 increases macroautophagy flux in HeLa cells analyzed by transmission electron microscopy

Transmission electron microscopy (EM) has historically been used to examine macroautophagy flux in cells via quantification of degradative compartments.^110,111^ The degradative compartments (DGCs) such as amphisomes, autolysosomes and lysosomes are characterized by amorphous electron-dense content and their number increases upon autophagy induction.^112–114^ We performed a four-point time course experiment in HeLa cells, treating them with 250nM Torin-1, 100nM BafA1, 2.5µM CCT020312 or vehicle for 4, 11, 15.5 and 24 h, followed by EM sample prep and image acquisition (see Materials and Methods). Representative images for Torin-1 and CCT020312 (N≥50) treatment show a significant increase in the number of electron-dense DGCs compared to vehicle treatment over time, unlike cells treated with the autophagy inhibitor BafA1 (Figure 2H, Data S2). Quantification of electron-dense DGCs reveal their statistically significant increase upon 11-h and 24-h CCT020312 treatment (Figure 2I). At 15.5h, vehicle control samples were not evaluated; however, CCT020312-treated cells at this time point look similar to that of the 11h treatment. Torin-1 treatment also caused a statistically significant augmentation in DGCs at all time points, but BafA1 treatment did not (Figure 2I). In addition, CCT020312-treated cells did not appear to cause formation of light-colored compartments representing DGCs with impaired hydrolytic activity that characterize BafA1-treated cells. Interestingly, contrary to electron-dense DGCs resulting from Torin-1 treatment, the DGCs in CCT020312-treated cells appeared to have secondary lighter-colored vesicles within them. This suggests that different cargoes are within the Torin-1 vs. CCT020312 DGCs, perhaps due to a different mode of macroautophagy activation. The electron micrographs revealed no major cellular alterations at the ultrastructural level, including no signs of cell toxicity (e.g. blebbing of nuclei that would suggest apoptosis) upon any of the treatments (Figure 2H, Data S2). Taken together, this quantitative EM analysis indicates that CCT020312 activates macroautophagy in HeLa cells in a time-dependent manner.

### CCT020312 does not modulate stress-responsive signaling pathways

Prior literature suggests that CCT020312 can activate the PERK signaling arm of the unfolded protein response (UPR) stress-responsive signaling pathway in U-2OS cells.^107^ Activation of PERK by the global ER stressor tunicamycin has been shown to activate autophagy, whereas PERK silencing reduces this effect.^108^ We thought it was possible that CCT020312 induces macroautophagy via PERK activation. However, a 4-h treatment of HEK293T cells with 1µM CCT020312 – a dose that is lower than the one that has been reported to activate PERK in MEF cells,^115^ but near the maximum dose used in our experiments to demonstrate macroautophagy activation – did not activate PERK as measured by RNA-Seq analysis of PERK-regulated target genes (Figure S3A, S3B). PERK was also not activated by treatment with CCT020312 (1µM) for 18 h in HEK293T cells expressing activating transcription factor 4 fused to Firefly luciferase (ATF4-Fluc), ATF4 being a PERK transcriptional reporter (Figure S3C).^116^ PERK activation was not observed in human cells at >10X the concentration of CCT020312 that activates macroautophagy in neurons (*vide infra*).

Co-treatment with the PERK inhibitor GSK2656157^117^ did not affect LD clearance induced by CCT020312 treatment of U251 cells (Figure S3D). Moreover, co-treatment with the PERK-mediated integrated stress response signaling inhibitor ISRIB^118^ did not impact the LC3-I to LC3-II conversion in U251 cells (Figure S3E). Similarly, LC3-I to LC3-II band conversion upon CCT020312 treatment was no different in *Perk^−/−^*mouse embryonic fibroblasts (MEFs) relative to *Perk^+/+^* MEFs, suggesting that PERK does not affect CCT020312-mediated macroautophagy activation in MEFs (Figure S3F). Overall, these data demonstrate that CCT020312-mediated LD clearance and macroautophagy activation are independent of PERK signaling.

Apart from canonical PERK signaling, it has been suggested that CCT020312 promotes protection against proteotoxic insults through PERK-dependent activation of the nuclear factor erythroid 2-related factor 2 (NRF2)-regulated oxidative stress response.^119^ NRF2 activity is implicated in the induction of autophagy,^120^ suggesting that CCT020312 could induce autophagy by activating NRF2. However, we did not observe CCT020312-dependent induction of NRF2 target genes by RNA-Seq (Figure S3A, S3B) or CCT020312-dependent activation of the NRF2-responsive antioxidant response element (ARE)-FLuc reporter in cell-based assays (Figure S3G). NRF2 can also be activated by radical oxidative stress.^121^ Thus, we tested whether treatment of HEK293T cells with 1µM, 5µM or 10µM CCT020312 for 2 h would change the ratio of reduced to oxidized glutathione as an indicator of oxidative stress (Figure S3H).^122^ However, no significant changes were observed, suggesting that CCT020312 does not induce radical oxidative stress.

In addition, we did not observe activation of the inositol-requiring enzyme 1/spliced X-box binding protein 1 (IRE1/XBP1s)^123^ or activating transcription factor 6 (ATF6)^124^ signaling arms of the UPR or the heat shock factor 1 (HSF1)-regulated heat shock response (HSR) by RNA-Seq (Figure S3A, S3B) or in cells expressing luciferase reporters specific to these three stress-responsive signaling pathways (Figure S3I, S3J, S3K). Collectively, these results indicate that CCT020312 does not promote macroautophagy induction through activation of stress-responsive signaling pathways, based on the lack of transcriptional changes after a 4-h treatment.

### CCT020312 activates TFEB transcriptional program in HeLa cells

Since the stress-responsive signaling pathways tested above were not activated, we explored other mechanisms by which CCT020312 treatment could activate macroautophagy. Transcription factor EB (TFEB) is well-established for its role in regulating autophagy-related genes via its translocation into the nucleus and transcriptional activation.^125,126^ To discern if CCT02312 could promote TFEB transcriptional activation, we conducted TFEB immunofluorescence (IF) imaging and immunoblotting after nuclear-cytoplasmic separation in HeLa cells. CCT020312 treatment dose-dependently increased the percentage of TFEB localized in the nucleus per cell (Figures 3A) and the TFEB nuclear/cytoplasmic ratio (Figure 3B) in HeLa cells.

**Figure 3:**
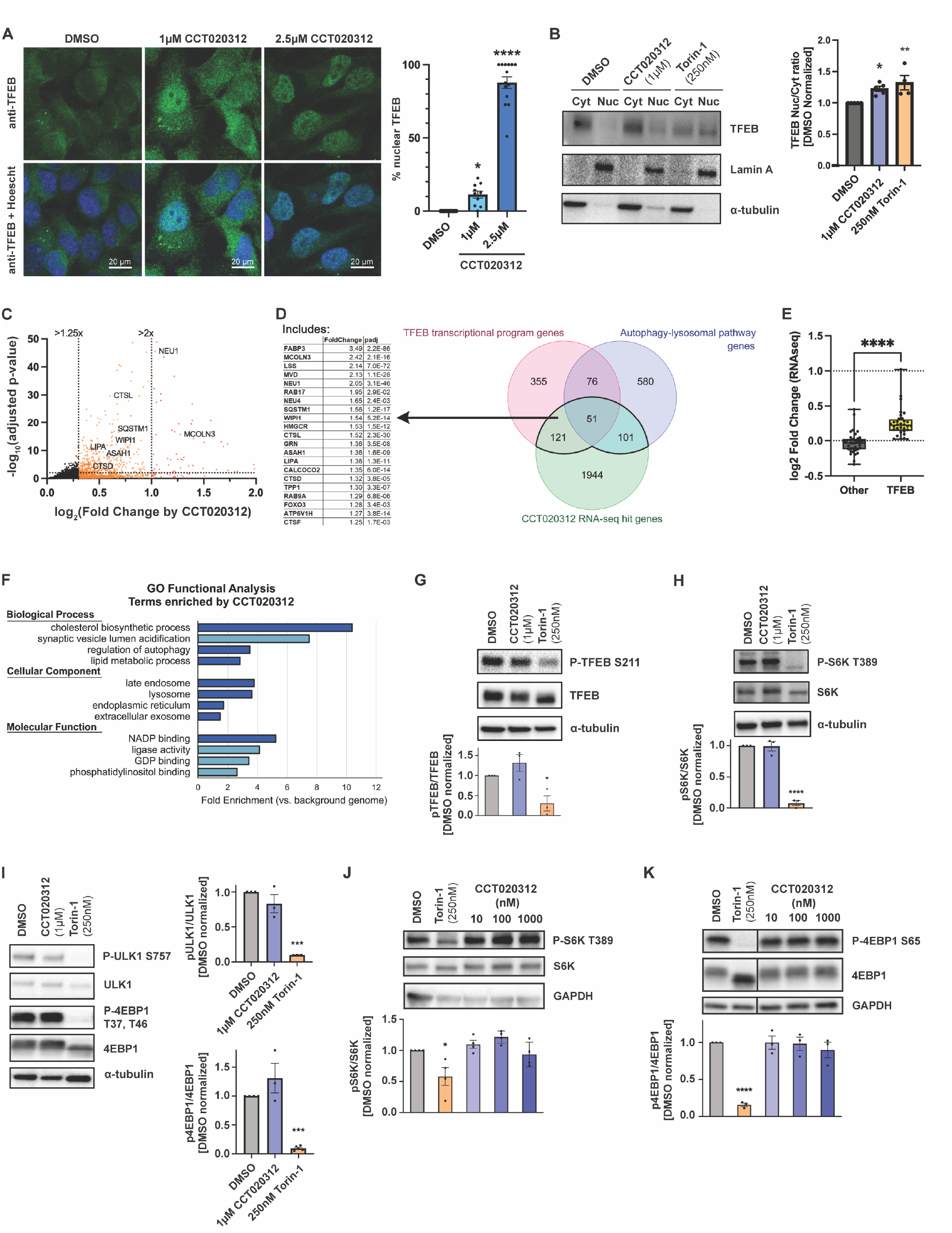
CCT020312 activates autophagic flux via TFEB transcriptional program independent of mTORC1 inhibition. (**A**) Representative images and bar graph representing the percentage of HeLa cells eliciting TFEB nuclear localization calculated from fluorescence microscopy images (representative images shown) taken after 20 hours of treatment with DMSO (vehicle), 1µM CCT020312 or 2.5µM CCT020312 (total 200-500 cells, n = 9-14 images per condition). * Dunnett’s multiple comparisons test (to DMSO) following one-way ANOVA p < .05, **** p < .0001. (**B**) HeLa cells treated with 250nM Torin-1, 1µM CCT020312, or DMSO for 20 hours, fractionated into cytosolic (Cyt) and nuclear (Nuc) fractions and immunoblotted for TFEB. Lamin A is nuclear protein and α-Tubulin is cytoplasmic protein loading control. * Dunnett’s multiple comparisons test following one-way ANOVA p < .05, ** p < .01. (**C**) Volcano plot depicting upregulated genes from RNA-Seq analysis of HeLa cells treated with 2.5µM CCT020312 for 20 h. Genes with fold change greater than 1.25 are colored orange, and genes with fold change greater than 2 are colored red. Some genes related to the autophagy-lysosome pathway and TFEB activation are labeled on the plot. Horizontal dotted line denotes adjusted p-value = 0.01. Axis adjusted for improved visualization of highlighted genes; some data is omitted by axis adjustment but full data can be seen in Figure S3M. (**D**) Venn diagram depicting gene overlap between significant hit genes (those with adjusted p-value < 0.05) from Supplemental Table 1 with published lists of genes from the autophagy-lysosomal pathway{The Proteostasis, 2023 #806} and the TFEB transcriptional program.{Palmieri, 2011 #725}{Sardiello, 2009 #714}{Settembre, 2011 #713} Selected genes of interest from the overlap between the groups are listed in the table next to the Venn diagram. (**E**) Gene set enrichment analysis of adjacent RNA-Seq data for activation of TFEB and other genes as described in Table 1 of {Grandjean et al, 2019}. (**F**) GO terms enriched by 20-h treatment with 2.5µM CCT020312 using DAVID Bioinformatics Database {Huang da, 2009 #805} for all genes from Supplemental Table 1 with adjusted p-value < .05 and fold change > 1 compared to DMSO. Terms displayed are those with the lowest DAVID false discovery rate and highest fold enrichment versus background genome (all genes identified in the experiment). Overly broad terms, such as “cytosol”, and redundant terms, such as “sterol metabolism” versus “cholesterol metabolism”, were removed from the bar chart for the sake of space. Bars in light blue represent GO terms that have a DAVID false discovery rate > 0.05 (but had statistically significant EASE score prior to post hoc correction). (**G**) HeLa cells treated with 250nM Torin-1, 1µM CCT020312, or DMSO for 20 h and immunoblotted for phospho-TFEB (S211). Bar graph depicts quantification of n = 3 replicates; one representative western blot shown. **** Dunnett’s multiple comparisons test following one-way ANOVA p < .0001. (**H**-**I**) HeLa cells treated with 250nM Torin-1, 1µM CCT020312, or DMSO for 20 h and immunoblotted for (**H**) phospho-S6K (T389), (**I**) phospho-ULK1 (S757) and phospho-4EBP1 (T37, T46). Bar graph depicts quantification of n = 3 replicates, one representative western blot shown. *** Dunnett’s multiple comparisons test following one-way ANOVA p < .001, **** p < .0001. (**J**-**K**) U251 cells treated with 250nM Torin-1, CCT020312 (10nM, 100nM or 1µM) or DMSO for 20 h and immunoblotted for (**J**) phospho-S6K (T389) and (**K**) phospho-4EBP1 (S65). Bar graph depicts quantification of n = 3 (in **K**) or n = 4 (in **J**) replicates; one representative western blot shown. * Dunnett’s multiple comparisons test following one-way ANOVA p < .05, **** p < .0001.

TFEB transcriptionally controls genes that belong to the Coordinate Lysosomal Expression and Regulation (CLEAR) network.^127^ To scrutinize whether the induced nuclear localization of TFEB by CCT020312 in HeLa cells is consistent with its transcriptional activation, we conducted RNA-Seq analysis of the same batch of cells treated with CCT020312 (2.5µM) for 20 h. RNA-Seq analysis revealed that CCT020312 treatment (2.5µM) significantly (adjusted p-value < 0.05) upregulated 1039 genes and downregulated 1177 genes (Supplemental Table 1), including 152 associated with the autophagy-lysosomal pathway (Figure 3C).^128^ To discern which genes are specifically modulated by the TFEB transcriptional program, we cross-referenced our findings with TFEB transcriptional program gene sets from previous studies,^125–127,129^ identifying 172 genes significantly altered by CCT020312 treatment (Supplemental Table 1, Figures 3C, 3D, S3L). These genes, including lysosomal acid hydrolases and selective autophagy receptors exhibited significant upregulation relative to vehicle-treated samples, establishing a link between CCT020312-induced gene expression changes and autophagy-lysosomal pathway regulation (NEU1: 2.1x DMSO, CTSL: 1.5x DMSO, ASAH1: 1.4x DMSO, LIPA: 1.4x DMSO, CTSD: 1.3x DMSO, CTSF: 1.3x DMSO, SQSTM1: 1.6x DMSO, CALCOCO2: 1.4x DMSO).

We next analyzed 30 CLEAR-regulated genes frequently used in publications to represent each step of the autophagy-lysosomal pathway (pre-initiation signaling, phagophore nucleation and elongation, substrate selection, autophagosome maturation, lysosome fusion and catabolism) reflecting the TFEB transcriptional program.^125–127^ Comparing their fold change in expression level upon CCT020312 treatment relative to other genes not affected by stress-responsive signaling pathways,^130^ we observed a significant induction of the TFEB transcriptional program by CCT020312 treatment (Figure 3E, Supplemental Table 2). Analysis of the bulk RNA-Seq data upon CCT020312 treatment also revealed upregulation of genes related to lipid metabolism, consistent with previous reports that TFEB controls cellular lipid metabolism (Figure S3M).^129^

Lastly, the 172 genes significantly altered by CCT020312 treatment were analyzed using the Database for Annotation, Visualization and Integrated Discovery (DAVID) bioinformatics tool.^131^ The top 4 terms from each category (Biological Process, Cellular Component, and Molecular Function) were plotted by fold enrichment (Figure 3F). Top terms from GO Biological Process include cholesterol biosynthetic process (GO:0006695), synaptic vesicle lumen acidification (GO:0097401), regulation of autophagy (GO:0010506), and lipid metabolic process (GO:0006629). Top terms from GO Cellular Component coincided well with our previous analysis of TFEB and autophagy-lysosomal pathway up-regulation, with late endosome (GO:0005770) and lysosome (GO:0005764) among the most highly enriched cellular components. Collectively, our results support the hypothesis that CCT020312 promotes autophagy activation in HeLa cells by triggering the TFEB transcriptional program.

### CCT020312 activates TFEB transcriptional program without mTORC1 inhibition

TFEB nuclear translocation is negatively regulated by phosphorylation at Ser142 and Ser211 by mTORC1.^132–134^ We assessed the phosphorylation state of TFEB at Ser211 to determine whether CCT020312 inhibits mTORC1. Treatment of HeLa cells with 250nM Torin-1 for 20 h led to a statistically significant reduction of phosphorylation of TFEB at S211 as expected (Figure 3G). In contrast, treatment with 1µM CCT020312 did not change phosphorylation of TFEB at S211 relative to the vehicle control (Figure 3G).

We next probed whether CCT020312 treatment alters mTORC1 signaling by interrogating the phosphorylation state of direct downstream substrates by western blot. These substrates include p70S6 kinase 1 (S6K), where phosphorylation occurs at Thr389;^135^ Unc-like autophagy activating kinase ULK1, where phosphorylation occurs at Ser757;^136^ and eukaryotic translation initiation factor 4E-binding protein 1 (4E-BP1), where phosphorylation occurs at Thr37/Thr46 or Ser65.^137^ S6K and 4E-BP1 are key regulators of mRNA translation activation,^138–140^ while ULK1 is phosphorylated during high mTOR activity to prevent autophagy activation.^136^ Phosphorylation of these downstream substrates was evaluated in HeLa and/or in U251 cells (Figure 3H-3K). In contrast to Torin-1 treatment, which served as a positive control, CCT020312 did not change the phosphorylation levels of any of these mTORC1 substrates in both tested cells. All together, these results suggest that CCT020312 activates autophagy through a different mechanism than one involving mTORC1 inhibition.

### CCT020312 increases macroautophagy flux in primary mouse cortical neurons

In this section, and several subsequent sections, we probed the ability of CCT020312 to activate macroautophagy in mouse and human neurons. We first evaluated the ability of CCT020312 to hasten macroautophagy flux in neurons using an optical pulse-labeling assay which employs a fusion between LC3B and the photoswitchable fluorescent protein Dendra2.^141^ Primary mouse cortical neurons were transfected with LC3-Dendra2 and subsequently irradiated with 405 nm light to irreversibly convert Dendra2 emission from green to red fluorescence (Figure 4A, left). Over 100 transfected neurons were monitored longitudinally by automated fluorescence microscopy to quantify the mean rate of decrease in red fluorescence over time, allowing a determination of the mean half-life of the photoconverted LC3 protein upon CCT020312 treatment (Figure 4A, right).^142^ We observed a dose-dependent reduction in the mean half-life of LC3-Dendra2 upon CCT020312 treatment above a concentration of 5nM, consistent with increased macroautophagy flux in mouse cortical neurons (Figure 4B). Administration of the inactive control compound carboxy CCT did not lead to a dose-dependent reduction in the mean half-life of LC3-Dendra2 (Figure 4B).

**Figure 4:**
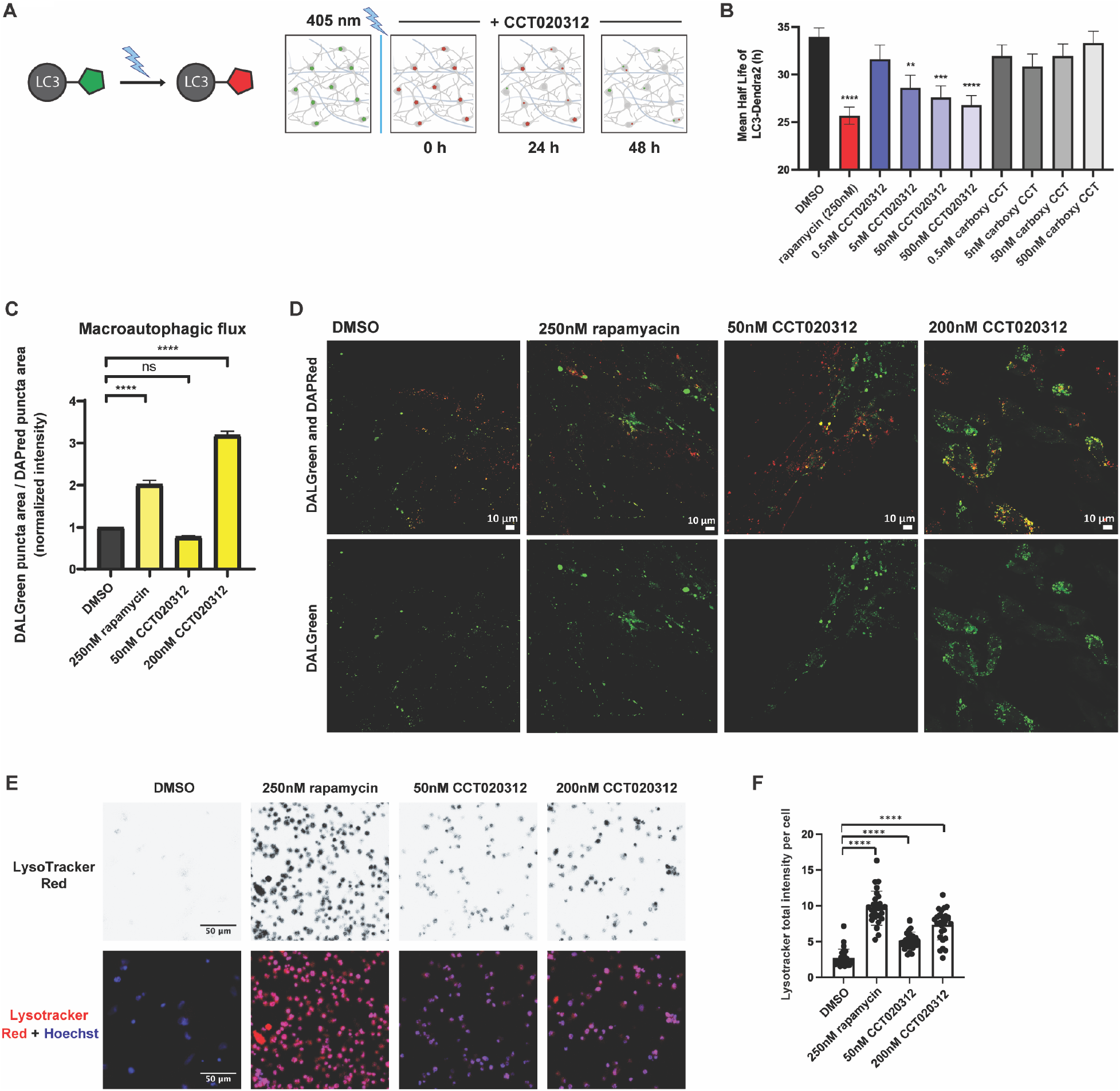
CCT020312 activates autophagic flux in neurons. **(A**) Optical pulse labeling enables tracking of protein degradation. Representation of an LC3B degradation over time assay with photoconvertible Dendra2 linked to LC3B. (**B**) Half-life of Dendra2-LC3 transfected in primary mouse cortical neurons upon treatment of indicated compounds (rapamycin = 250nM). DMSO serves as control. Bars and error bars depict mean + SEM, n = 116. * Dunnett’s multiple comparisons test following one-way ANOVA p < .05, ** p < .01, *** p < .001, **** p < .0001. (**C**) Quantification of autophagic flux ratio, defined as the total area of autolysosomes (green puncta) divided by the total area of autophagosomes and autolysosomes (red puncta) in human iPSC-derived cerebrocortical neurons after 24-h treatment with CCT020312 (50nM or 200nM), 250nM rapamycin or DMSO. Bars and error bars depict mean + SD, N = 4, n = 21. **** Dunn’s multiple comparisons test following Kruskal-Wallis p < .0001. (**D**) Representative images of human iPSC-derived cerebrocortical neurons stained with DALGreen and DAPRed after 24-h treatment with CCT020312 (50nM or 200nM), 250nM rapamycin or DMSO. (**E**) Representative confocal images quantified in (**F**). Lysotracker Red channel has been changed to grayscale when separate for improved contrast, but the merge depicts Lysotracker Red in red and Hoescht stain in blue. (**F**) Quantification of the normalized LysoTracker intensity signal of live human iPSC-derived cerebrocortical neurons after 24-h treatment with indicated compounds. N = 4, n = 28. **** Dunn’s multiple comparisons test following Kruskal-Wallis p < 0.0001.

### CCT020312 increases macroautophagy flux in human iPSC-derived cortical neurons

We next tested the activity of CCT020312 in human cortical neurons derived from induced pluripotent stem cell (iPSC) differentiation.^143,144^ To monitor macroautophagic flux in this cell line we utilize the amphiphilic fluorescent probes DALGreen and DAPRed. These 1,8-naphthalimide probes contain a terminal amine, which mimics the intramembrane phospholipid, phosphatidylethanolamine. Consequently, the probes become embedded into the growing phagophore membrane and render the corresponding downstream autophagic vesicles (autophagosomes and/or autolysosomes) fluorescent.^145^ DALGreen fluorescence is enhanced by acidification of membrane-delimited compartments, which occurs after autophagosomes fuse with lysosomes. In contrast, DAPRed fluorescence is not affected by pH changes and therefore measures total autophagosome and autolysosome levels.^146^ After a 24-h treatment of human iPSC-derived cortical neurons with CCT020312, DALGreen and DAPRed were added for the final 1 h (see Materials and Methods). Relative to DMSO-treated neurons, neurons treated with CCT020312 exhibited a dose-dependent increase in macroautophagy flux (increase in ratio of green to red fluorescence area, as this represents a greater number of autolysosomes forming relative to the number of autophagosomes forming; Figure 4C, 4D). Control autophagy activator rapamycin treatment analogously increased macroautophagic flux in the same assay (Figure 4C, 4D). LysoTracker Red is a lysosomotropic dye used to stain acidic compartments in the cell and its increased fluorescence intensity suggests larger and/or greater number of lysosomes in CCT020312-treated neurons (Figure 4E and 4F). Together, these results demonstrate that CCT020312 treatment enhances autophagic flux in human iPSC-derived cortical neurons above a concentration of 50nM.

### Tau levels reduced by CCT020312 treatment in a cellular tauopathy frontotemporal dementia disease model

Encouraged by the ability of CCT020312 treatment to activate autophagy in neurons in the nM concentration range, we next set out to test whether it could ameliorate ND phenotypes. Abnormal tau protein conformations and post-translational modifications, as well as increased protein burden in the CNS leading to tau aggregation and neuronal toxicity, are hallmarks of a group of disorders referred to as tauopathies, which include AD and frontotemporal dementia (FTD). In this group, like in other neurodegenerative disorders, autophagy impairment in disease and pharmacological enhancement as potential therapeutic strategies are of great relevance.^26,28,29,33,40,51^ Previously, CCT020312 had been shown to clear pathological tau species in a mouse model–at that time it was hypothesized to be a PERK activator (not consistent with evidence summarized herein above).^147^ To investigate the effects of CCT020312 in human neurons derived from FTD patients, we utilized an iPSC-derived neural progenitor cell (NPC) line designated MGH-2046-RC1, derived from an individual carrying the Tau-P301L autosomal dominant mutation. When differentiated into a mixed population of neural types, these patient-specific ex vivo neurons represent well-established models for studying tauopathy phenotypes.^148–150^

Tau-P301L neurons differentiated for six weeks and treated with CCT020312 at concentrations between 10nM and 10µM for 24 h (Figure 5A), revealed a concentration-dependent reduction in tau (Figure 5A and 5B), including total tau (TAU5 antibody) and phospho-tau (P-tau) levels. Specifically for P-tau, both the AT8 antibody and the Ser396 antibody recognize P-tau epitopes previously associated with human post-mortem pathology, Ser202 / Thr205 and Ser 396, respectively, and were employed for quantification. The maximum effect over 24 h was achieved at 10µM CCT020312, which reduced tau levels by 80% relative to vehicle control (Figure 5B). Importantly, a reduction in monomeric P-tau S396 (∼55 kDa band) as well as high MW oligomeric P-tau S396 (at ≥250 kDa) were both observed (Figure 5A). CCT020312-mediated autophagy activation in the iPSC-derived tau-P301L neurons was demonstrated by a dose-dependent increase in LC3-II, by western blot analysis (Figure S4A and S4B). The effect of CCT020312 on other macroautophagy and lysosomal activity markers such as lysosomal-associated membrane protein 1 (LAMP1), p62 and the lysosomal cathepsin D (CTSD) was also positive. CCT020312 treatment achieved a maximal effect on their levels at a concentration of 10nM (Figure S4C). Rapamycin, which also reduced tau and P-tau levels in human ex vivo neurons,^148^ had a different profile on LAMP1 and p62 levels with much lower upregulation of LC3-II (Figure S4D), indicative of a distinct mechanism of action in comparison to CCT020312.

**Figure 5:**
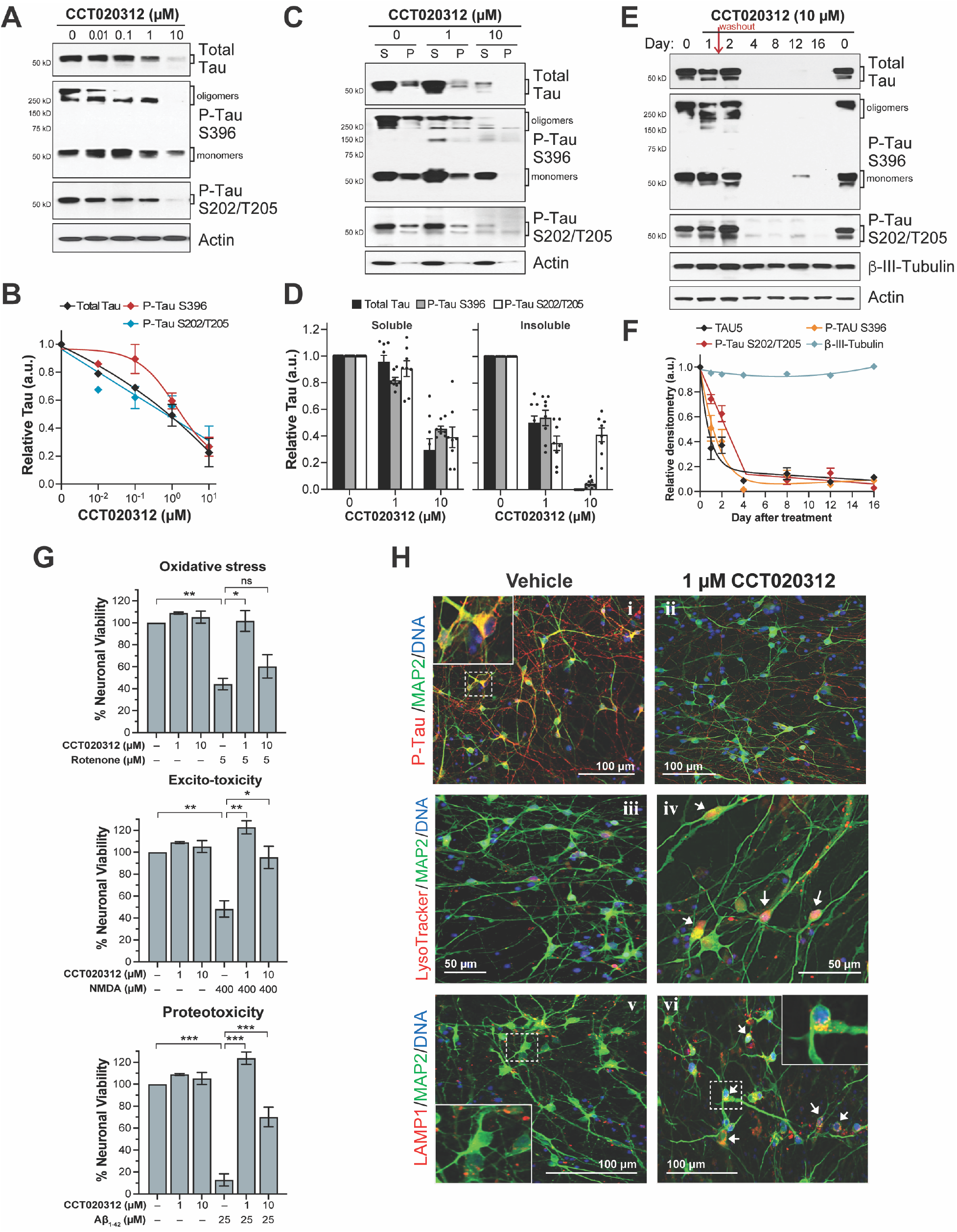
CCT020312 rescues tauopathy phenotypes in FTD patient iPSC-derived tau-P301L neurons. (**A**) CCT020312 concentration effect on total tau (TAU5) and P-tau S396 and S202/T205 (AT8) of P301L neurons after 24-h treatment. Detection of monomeric (∼55 kDa) and high MW oligomeric (≥250 kDa) P-tau S396. Brackets depict bands used for densitometry analysis in (**B**). (**B**) Western blot densitometry, quantification of bands’ pixel intensity (in arbitrary units, a.u.). Data points represent mean densitometry normalized to actin and relative to vehicle (DMSO) ± SD (n = 2). (**C, D**) CCT020312 effect on tau solubility in P301L neurons. Western blot (**C**) and densitometry (**D**) quantification of total tau (TAU5), P-tau S396 and P-tau S202/T205 (AT8) in the soluble (S) and insoluble-pellet (P) fractions of neurons treated for 24 h. Brackets (**C**) indicate the bands used for analysis in (**D**). Graph bars represent mean densitometry ± SD (n = 2) for soluble (left) and insoluble (right) tau levels relative to vehicle samples (0µM, DMSO). (**E, F**) Single 24-h dose of CCT020312 has a prolonged effect on tau reduction. Tau-P301L neurons were treated for 24 h, followed by compound washout (red arrow). Total tau and P-tau were measured by western blot (**E**) over a period of 16 d post-treatment. Graph data points (**F**) represent mean densitometry (bands within brackets) ± SD (n = 2), relative to day 0/Vehicle DMSO. β-III tubulin served as a control for neuronal integrity during the time course. (**G**) Rescue of stress vulnerability in a tauopathy neuronal model by CCT020312. The stressors rotenone (top), NMDA (center) and Aβ (1-42) (bottom) reveal P301L neuronal stress vulnerability by causing approximately 60%-90% loss in viability. Pre-treatment with 1µM or 10µM CCT020312 rescued neuronal viability to 60%-100% of vehicle (DMSO). Graph bars represent mean viability relative to vehicle-treated neurons (100%) ± SD (n = 2, and four technical replicates). Two-tailed unpaired Student’s T-test ^ns^ p > 0.05; * p < 0.05, ** p < 0.01, *** p < 0.001. **(H**) Fluorescence imaging of tau-P301L neurons treated for 24 h with vehicle (DMSO) or 1µM CCT020312, probed with LysoTracker Red, and the antibodies for P-tau PHF-1 (red), LAMP1 (red) and the neuronal marker MAP2 (green). White arrows point to the accumulation of LysoTracker (ii) and LAMP1 (vi) in the cell body of neurons treated with CCT020312. Insets show zoomed images of the dotted boxes.

The influence of CCT020312 treatment on detergent-insoluble aggregated forms of tau, the quaternary structures predicted to be more proteotoxic, was investigated next. Neurons were treated with CCT020312 at the concentrations that caused the strongest reduction in tau (1µM and 10µM) and after 24 h, detergent-based protein fractionation of the neuronal lysates was performed as previously described.^151–153^ By western blot analysis (Figure 5C) and semi-quantitative analysis (Figure 5D), the effect of CCT020312 on soluble (S) and insoluble (P) tau fractions was measured. The 1µM dose had a negligible effect on the soluble fraction of tau (Figure 5D, left) but reduced insoluble total tau and P-tau by ≥50% (Figure 5D, right). At the higher dose of 10µM, CCT020312 reduced soluble tau by ∼50% and insoluble total tau by 100% (as detected by the TAU5 antibody), and nearly completely reduced P-tau S396, while insoluble P-tau S202/T205 was reduced by about 60% (Figure 5D, right). These results indicate a stronger effect by CCT020312 on the insoluble fraction of tau of P301L neurons.

To evaluate the long-term effects of a 24-h 10µM CCT020312 treatment of P301L neurons on Tau pathology over a 16 d post-treatment period, the culture medium was replaced with CCT020312-free medium at 24 h (“washout”, Figure 5E, *red arrow*). Neurons were collected over a time course between 1 and 16 d post-treatment, and total tau, P-tau S396, and S202/T205 levels were measured by western blot (Figures 5E and 5F). Day zero corresponds to vehicle-treated samples. After 24 h of treatment (d 1), tau was reduced by ∼50% (Figure 5F). Notably, after media washout, the levels of total tau and P-tau continued to decrease up to d 4, bringing tau levels further down to 0%-10% of vehicle-treated samples (Figure 5F). A remarkable prolonged loss of total tau and P-tau was observed up to d 16 post-treatment. In parallel, the effect of CCT020312 treatment on neuronal integrity and neurite outgrowth, by itself or because of tau reduction, was evaluated by measuring levels of the microtubule protein β-III-tubulin, which remained unchanged during the 16d period (Figure 5E, 5F). This suggests that in parallel with CCT020312-mediated tau reduction, neuronal integrity was not affected, per β-III-tubulin and actin levels over time.

### CCT020312 treatment decreases tau-P301L neuronal vulnerability to stress

In FTD patient iPSC–derived mutant-tau neurons, tau-mediated toxicity is assessed by increased neuronal vulnerability to stress.^64^ Stressors that reveal this cell death phenotype in tau-P301L neurons include treatment with the mitochondrial electron transport chain inhibitor rotenone (5µM), high-dose administration of the excitatory neurotransmitter NMDA (400µM), and exposure to the highly aggregation-prone peptide Aβ(1-42) (25µM). Treatment with these stressors led to a loss of viability of P301L neurons: 60% loss by rotenone administration (Figure 5G, top panel), 60% loss associated with NMDA treatment (Figure 5G, middle panel) and >80% loss by exposure to Aβ(1-42) (Figure 5G, lower panel). When neurons were pretreated with the autophagy activator CCT020312 for 8 h and then cotreated with the stressor for an additional 16 h, there was a consistent protective effect and rescue of neuronal viability (Figure 5G). This was particularly the case for the lower dose of 1µM CCT020312, which protected neurons from the stressors maintaining viability at ∼100%.

Representative immunofluorescence imaging of tau-P301L neurons treated with 1µM CCT020312 show a clear decrease in P-tau S396/S404 staining relative to vehicle-treated neurons (PHF1 antibody, Figure 5H, compare panels i and ii) without affecting the microtubule-associated protein 2 (MAP2) marker staining or neuronal density/morphology. Neurons treated with CCT020312 also show upregulation and accumulation of macroautophagy and lysosomal pathway markers, mainly in the cell body, as shown by staining with the lysosomal fluorophore LysoTracker Red (Figure 5H, compare panels iii and iv), and increased staining of LAMP1 (Figure 5H, compare panels v and vi). These results suggest that tau reduction is a consequence of pharmacological macroautophagy activation by CCT020312.

### Reducing Aβ peptides in AD direct differentiated neurons

Intrigued by the ability of CCT020312 treatment to protect tau-P301L neurons from Aβ(1-42)-mediated proteotoxic stress, we investigated whether CCT020312 treatment can clear Aβ peptides within AD patient–derived neurons. In previous studies, direct lineage reprogramming has enabled the conversion of adult somatic cells directly into neurons without intermediate pluripotent cell stages.^154,155^ These directly induced neurons (iNs) can substantially preserve epigenetic marks of aging and recapitulate molecular factors involved in the etiology and pathogenesis of AD.^156,157^ The lack of intracellular clearance of Aβ peptides is associated with Aβ aggregation and neuronal cell death. Increasing evidence indicates that defective lysosomes in neurons promotes intraluminal formation of Aβ fibrils within the lysosomal compartment, and it has been hypothesized that these contribute to extracellular plaque formation in AD brains.^26^ We generated iNs from the fibroblasts of a sporadic AD (sAD) patient and from a hereditary AD donor harboring a A431E *PSEN1* mutation (AD-PSEN1).^158^ Direct lineage reprogramming of fibroblasts into iNs was accomplished by overexpression of pro-neuronal transcription factors (Brn2, Ascl1, Myt1l, and Ngn2). The AD-PSEN1 iNs had increased intracellular levels of total Aβ (∼31% increase) and Aβ(1-42) (∼38% increase; more proteotoxic isoform), relative to levels in sAD iNs, as measured by indirect immunofluorescence in neuronal cell bodies (Figure 6). Treatment by the β-secretase inhibitor IV (BACE-1 inhibitor, 1µM) for 48 h revealed that inhibition of the initial step in amyloid precursor protein processing by β-secretase inhibition lowers the production of total Aβ and Aβ(1-42) in both sAD and AD-PSEN1 iNs by 50%-66%. Rapamycin administration (250nM; 48 h) also mediated the reduction of intracellular Aβ peptides in both sAD and AD-PSEN1 iNs by about 50%. Notably, CCT020312 treatment (125nM, 250nM and 500nM; 48 h) significantly reduced total Aβ and Aβ(1-42) levels by about 33%-51% in sAD iNs and about 38%-50% in AD-PSEN1 iNs. Moreover, CCT020312 at doses used in this experiment exhibited no discernable cytotoxicity to iNs in the form of degenerated neurites or cell bodies lifting from the coverslip after the 48-h treatment period. Collectively, the rapamycin and CCT020312 results suggest that activation of macroautophagy has a beneficial effect by reducing intracellular Aβ burden in sporadic and familial AD neurons derived from patient fibroblasts.

**Figure 6:**
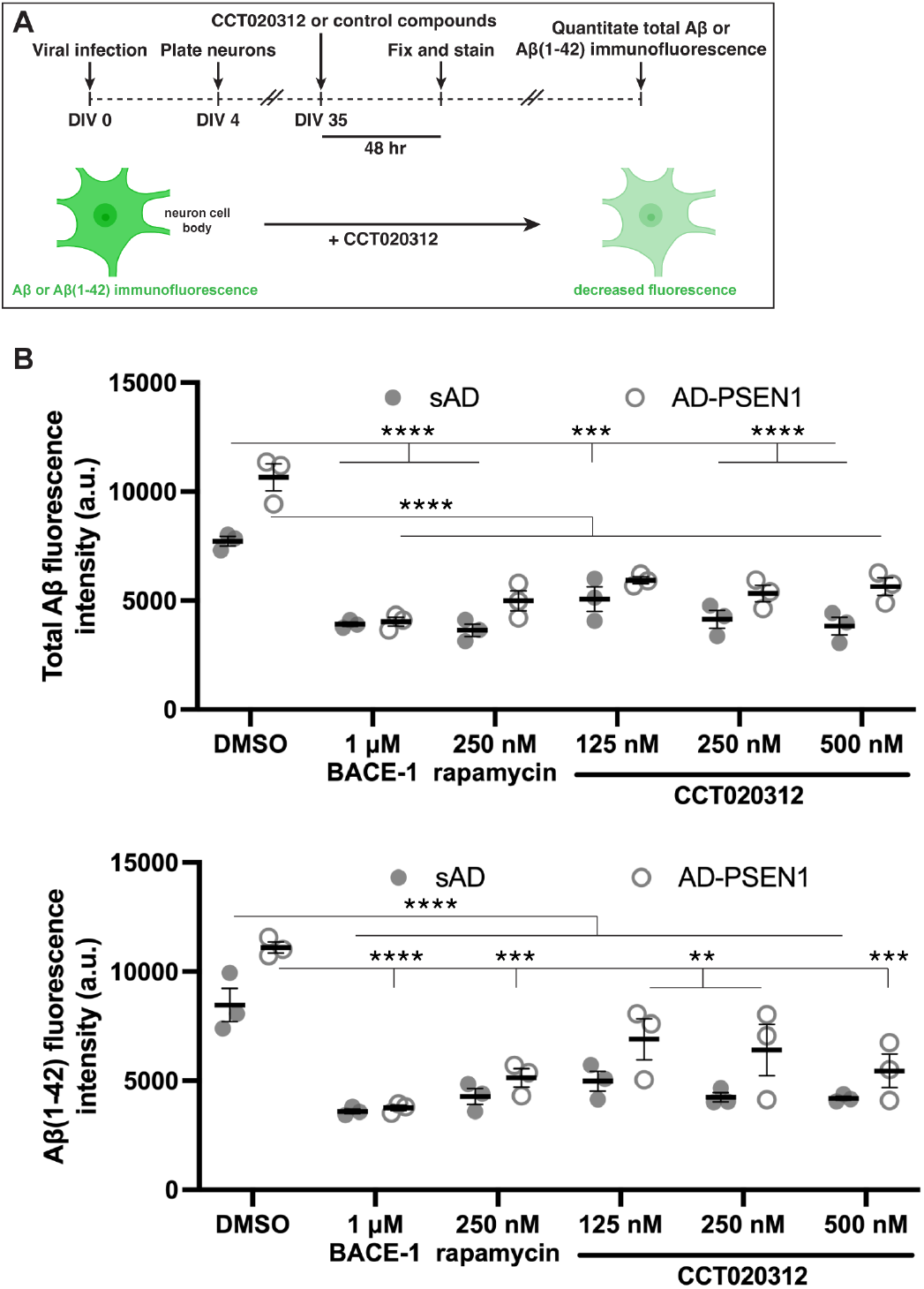
CCT020312 reduces Aβ and Aβ(1-42) peptide levels in AD-patient iNS. **(A)** Scheme depicting AD-patient iNS treatment regimen with CCT020312. **(B)** Quantification of mean fluorescence intensity of total amyloid beta (Aβ) and the aggregation-prone peptide species, Aβ(1-42), in the cell body of directly induced neurons (iNs) from fibroblasts of patients with sporadic Alzheimer’s disease (sAD) or AD carrying PSEN1 A431E mutation (AD-PSEN1). Treatment of 1µM BACE-1 inhibitor (β-Secretase Inhibitor IV, Sigma-Aldrich), 250nM rapamycin, or CCT020312 at 125nM, 250nM or 500nM for 48 h. Statistical analysis was performed with one-way ANOVA and Holm-Sidak’s post hoc test. ** p < 0.01, *** p < 0.001, **** p < 0.0001 (vs. DMSO). n = 3, means of each experimental replicate shown on graph. Cell number quantified n ≥ 35 per experimental replicate. Data represented as dot plot with standard error bars.

### CCT020312 reduces axonal endoggresomes in a familial prion disease model

Intraneuronal non-native protein aggregates that compromise axonal vesicular trafficking are a common and early feature of many neurodegenerative diseases.^159,160^ These aggregates are often located within dystrophic axonal domains, where accumulation of organelles (mitochondria), neurofilaments, and endosomal vesicles including enlarged lysosome-like structures that contain incompletely digested material, also occurs.^161–165^ A mutant prion protein (PrP) fusion construct containing an expanded 14 octapeptide repeat (PrP^PG14^; there is a 9 octapeptide insertion in wild-type PrP (PrP^WT^); see Figure S5A) is associated with Gerstmann-Sträussler-Scheinker (GSS), a human familial prion disorder.^166^ Previous work showed that expression of PrP^PG14^ in cultured murine hippocampal neurons results in the formation of enlarged endolysosomes along axons containing PrP^PG14^ aggregates, known as endoggresomes (aggregates within endosomes) (Figure S5A, bottom panels).^165,167^ Mutant and WT PrP sequences contain a PrP signal sequence (SS) and PrP glycosylphosphatidylinositol (GPI)-anchoring sequence to ensure proper transport through the secretory pathway within the lumen of vesicles and compartments.^167^ Endoggresomes form within axonal dystrophic swellings following the active transport of PrP^PG14^-carrying Golgi-derived vesicles into the axon by their association with the small endolysosomal GTPase ARL8b, which recruits kinesin-1 and the multiunit homotypic fusion and protein sorting (HOPS) complex subunit VPS41 onto PrP^PG14^-carrying endosomes. In the axon, HOPS-mediated homotypic fusion of PrP^PG14^-containing vesicles result in the formation of enlarged endoggresomes, which disrupt calcium dynamics and impair the axonal transport of vesicular cargos, reducing neuronal viability.^167^ Importantly, expression of PrP^PG14^ in primary cultured neurons results in an aberrant LysoTracker signal in axons, suggesting reduced lysosomal clearance capacity within endolysosomes.^167^

To determine whether CCT020312 could degrade PrP^PG14^ endoggresomes within axons, primary hippocampal neurons were transfected with PrP^PG14^-mCh at 7 d in vitro (DIV) and at 8 DIV were treated for 2 h with different concentrations of CCT020312. Endoggresome densities were quantified at DIV 9 on fixed samples (Figure 7A). Treatment of neurons with CCT020312 above a median concentration of 50nM significantly reduced PrP^PG14^-mCh endoggresome densities in axons in a dose-dependent manner, resulting in densities of vesicles similar to those in neurons expressing PrP^WT^-mCh (Figures 7B, 7C), which were unaffected by treatment. Because axonal PrP^PG14^-mCh endoggresomes are well formed by DIV 8, but densities continue to increase by DIV 9 (Figure 7D), the reduced endoggresome densities observed following treatment with CCT020312 at DIV 8 indicate that CCT020312 can efficiently clear endoggresomes from mammalian axons, in addition to potentially inhibiting their formation (Figure 7C, pink dot plots). As a control, we observed that treatment of neurons with the inactive analog carboxy CCT did not alter PrP^PG14^-mCh endoggresome densities, while also not influencing PrP^WT^-mCh vesicle numbers (Figure S5B). To rule out the possibility that the reduced PrP^PG14^-mCh endoggresome densities were a result of observational bias of neurons that survived putative CCT020312-mediated cell toxicity, we quantified neuronal viability using a NucGreen Dead cell death assay (see Materials and Methods). Quantification of the percentage of dead neurons at DIV 9 using NucGreen Dead (dead cell count), alongside a Hoechst 33342 (total cell count) marker after treatment with CCT020312, with the inactive carboxy CCT, or with the DMSO vehicle control following the same treatment paradigm as used above (Figure 7A) indicated that compound concentrations produced similar cell death levels (mean ∼30%; Figure S5C). To test whether activation of lysosomal degradation by known autophagy activators could reduce PrP^PG14^ axonal endoggresomes, hippocampal neurons were treated with the established mTORC1 inhibitor rapamycin using the same treatment paradigm (Figure 7A). We observed an analogously reduced endoggresome burden in neurons expressing PrP^PG14^-mCh in the presence of rapamycin as in those treated with CCT020312, while there were no changes in PrP^WT^-mCh vesicle densities (Figure S5D). Altogether, these data highlight the efficacy of CCT020312 in reducing PrP^PG14^-mCh endoggresomes at low nanomolar concentrations, indicating that activating macroautophagy is a viable and generalized strategy to clear toxic axonal PrP^PG14^ endoggresomes from mammalian axons.

**Figure 7:**
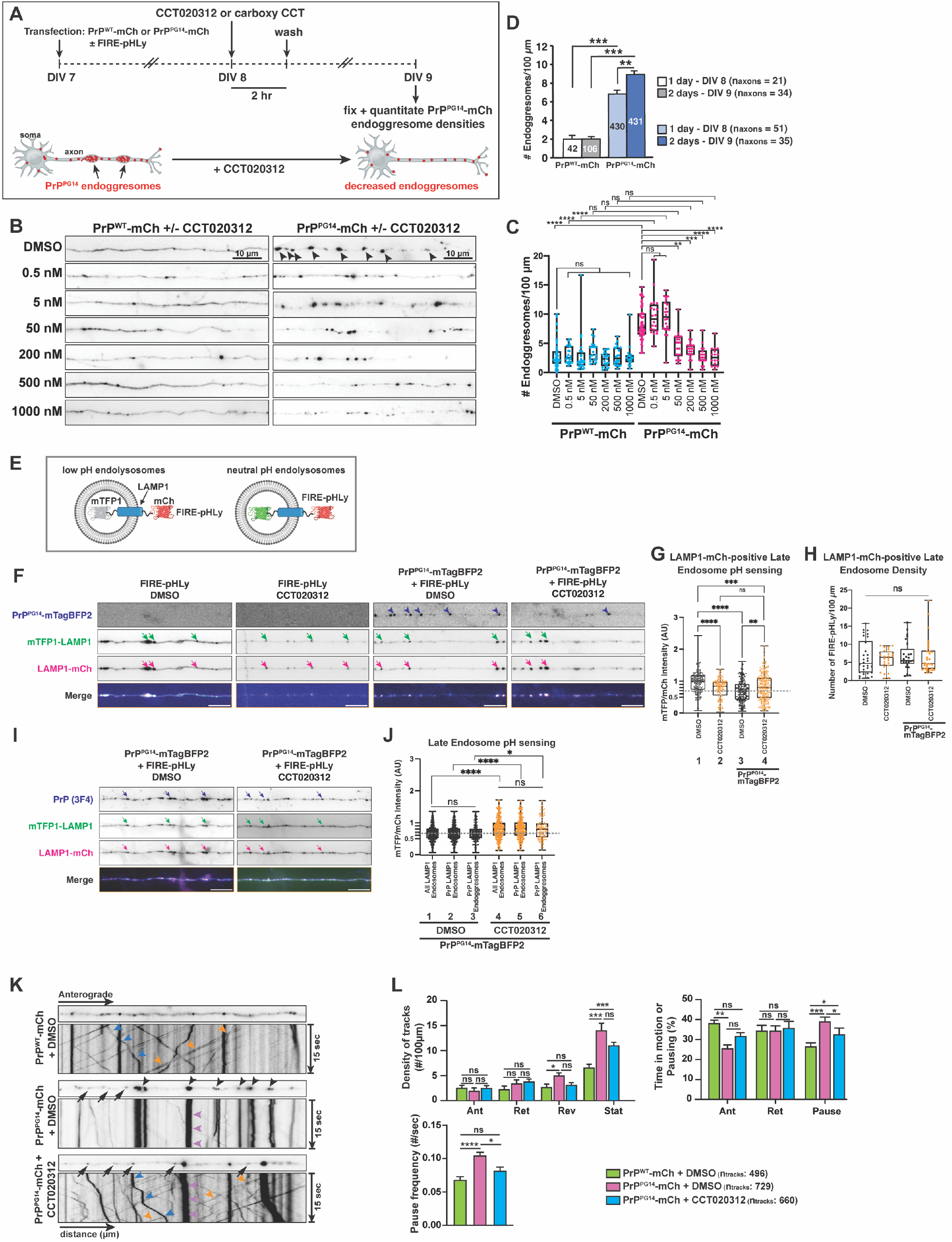
CCT020312 clears PrP^PG14^ endoggresomes from primary mouse hippocampal neurons. (**A**) Schematic diagram of experimental layout for transfection and treatment of neurons with PrP plasmids and CCT020312 or DMSO (vehicle), respectively. (**B**) Representative fluorescence images of axons of neurons expressing PrP^WT^-mCh or PrP^PG14^-mCh treated with DMSO (vehicle) or with increasing doses of CCT020312. Inverted images are shown. Arrowheads indicate endoggresomes. (**C**) PrP^WT^-mCh vesicle and PrP^PG14^-mCh endoggresome densities from (**B**), showing a statistically significant, dose-dependent decrease in axonal endoggresome densities starting at 50nM CCT020312. ** p < .01, *** p < .001, **** p < .0001, ns = not statistically significant. One-way ANOVA, Šidák correction. (**D**) PrP^WT^-mCh vesicle and PrP^PG14^-mCh endoggresome densities one and two days post-transfection of hippocampal neurons. (**E**) Schematic of FIRE-pHLy pH biosensor developed by Chin et al.{Chin, 2021 #819} (**F**) Representative fluorescence images of neutral endolysosomes (mTFP1-LAMP1) and total endolysosomes (LAMP1-mCh) in axons with or without PrP^PG14^-mTagBFP2 overexpression. Neurons were treated with DMSO or CCT020312. Inverted images are shown. Arrowheads indicate PrP^PG14^-mTagBFP2 endoggresomes, and arrows indicate LAMP1-positive endolysosomes. Scale bars, 10 μm. (**G**) Ratio of mTFP1 to mCh intensities of each LAMP1-positive endolysosome, and (**H**) density of endolysosomes in axons. Box plots represent median and interquartile range. **p < 0.01, ***p < 0.001, and ****p < 0.0001; Mann-Whitney test. AU, arbitrary units. (**I**) Representative fluorescence images of neutral endolysosomes (mTFP1-LAMP1) and total endolysosomes (LAMP1-mCh) in axons of PrP^PG14^-mTagBFP2-expressing neurons with DMSO or CCT020312 treatment. PrP^PG14^ endolysosomes were labeled with a 3F4 antibody. Arrowheads indicate LAMP1-positive PrP endoggresomes, and arrows indicate LAMP1-positive PrP^PG14^ endolysosomes that are not endoggresomes. Inverted images are shown. (**J**) Ratio of mTFP1 to mCh intensity of LAMP1-positive endolysosomes, LAMP1-positive PrP^PG14^ endolysosomes, and for LAMP1-positive PrP^PG14^ endoggresomes. Box plots represent median and interquartile range. *p < 0.05, and ****p < 0.0001; Mann-Whitney test. AU, arbitrary units. (**K**) Representative kymographs of axonal transport trajectories in neurons expressing PrP^WT^-mCh or PrP^PG14^-mCh treated with CCT020312 or DMSO. Arrowheads indicate anterograde (blue), retrograde (orange), or stationary (purple) trajectories of PrP-mCh vesicles or endoggresomes. Inverted images are shown. (**L**) Density of tracks, time spent in motion, and pause frequencies of PrP^WT^-mCh or PrP^PG14^-mCh vesicles in axons treated with CCT020312 or DMSO (vehicle). Dunn’s multiple comparisons test following Kruskal-Wallis * p < .05, ** p < .01, *** p < .001, **** p < .0001, ns = not statistically significant

### CCT020312 restores the acidification profile of endolysosomes within neurons challenged with prion protein mutations

Previous studies showed lower LysoTracker signal in axons of neurons expressing PrP^PG14^-mCh, suggesting decreased lysosomal degradation capacity.^167^ To further probe the nature of this lysosomal dysfunction and to investigate how treatment with CCT020312 clears these aggregate structures, we surveyed the pH acidification landscape of endolysosomes and endoggresomes in neurons expressing PrP^PG14^ (or not), followed by treatment with CCT020312 or vehicle (DMSO). Murine hippocampal neurons were co-transfected with PrP^PG14^-mTagBFP2 (blue fluorescent protein 2) and with Fluorescence Indicator REporting pH in Lysosomes (FIRE-pHLy), a recently developed ratiometric biosensor that reports on pH measurements of single endolysosomes.^168^ Co-transfections followed the same earlier scheme (Figure 7A). FIRE-pHLy is a dual mTFP1 (teal [cyan] FP)-human LAMP1-mCh reporter system that is targeted to endolysosomes via the LAMP1 endolysosomal protein (Figure 7E). mCh is in the cytosol and is acid-insensitive, while mTFP1 is acid-sensitive and localized within the lumen of endolysosomes (Figure 7E). An advantage of FIRE-pHLy over other pH reporters is its p*K*_a_ range from 3.5-6.0, allowing for the sensing of a wider range of pH values, including acidic ones.^168^ In this system, the mCh signal is used as a reference for the location of endolysosomes, while the mTFP1 changes signal intensity in response to the pH of the lumen of LAMP1 endolysosomes. The relative intensity differences between the fluorophores provides a ratiometric readout of the pH content of single endolysosomes.

We first imaged and quantitated mTFP1/mCh intensity ratios of individual LAMP1-mCh-positive endolysosomes in axons of neurons transfected with FIRE-pHLy alone (without PrP^PG14^-mTagBFP2), and treated with DMSO or with CCT020312 (200nM) for 2 h – a dose found to significantly clear axonal PrP^PG14^-mTagBFP2 endoggresomes (Figure 7B). We observed a significantly reduced mTFP1/mCh intensity ratio distribution of LAMP1-mCh-positive endolysosomes in axons compared to those in axons of DMSO-treated control neurons (Figure 7F, 7G). These data indicate an overall increased acidity in endolysosomes of neurons treated with CCT020312, consistent with a role of this molecule as an autophagy activator. Expression of PrP^PG14^-mTagBFP2 alone (+ DMSO but without CCT020312 treatment) resulted in a significantly lower mTFP1/mCh ratio (Figure 7G), indicating a hyperacidity distribution profile of LAMP1-mCh-positive endolysosomes compared to those in DMSO-treated neurons transfected with FIRE-pHLy alone (Figure 7F, 7G). Notably, treatment of neurons expressing PrP^PG14^-mTagBFP2 with CCT020312 rescued this phenotype, raising the pH distribution profile to similar levels as those of control neurons transfected with FIRE-pHLy alone and treated with CCT020312 (Figure 7F, 7G). Importantly, the densities of LAMP1-positive endolysosomes in axons were not significantly different across those conditions (Figure 7H).

To further determine whether the effect of CCT020312 in rescuing the pH profile of LAMP1-mCh endolysosomes extended to only PrP^PG14^-positive endolysosomes and endoggresomes, we quantitated mTFP1/mCh ratios in single PrP^PG14^-positive endolysosomes and endoggresomes in hippocampal neurons co-transfected with PrP^PG14^-mTagBFP2 and FIREpHLy (± DMSO or CCT020312). To clearly visualize PrP^PG14^ endolysosomes and endoggresomes, immunofluorescence was performed with an antibody specific to an introduced human 3F4 epitope in the PrP^PG14^-mTagBFP2 construct (Figure S5A). Consistent with the earlier results (Figure 7F, 7G), we detected an increased mTFP1/mCh intensity ratio distribution upon CCT020312 treatment in LAMP1-positive endolysosomes (Figure 7I, 7J, dot plots 1 vs 4). Importantly, this increase was also present in the sub-population of LAMP1-positive endolysosomes containing PrP^PG14^ (Figure 7J, dot plots 2 vs 5), and in LAMP1-positive PrP^PG14^ endoggresomes (Figure 7J, dot plots 3 vs 6). These data indicate that CCT020312 can modulate the acidity profile of PrP^PG14^ endolysosomes and endoggresomes in axons, suggesting that CCT020312 reduces PrP^PG14^ endoggresomes at least in part via modulating endolysosomal acidification in axons.

### CCT020312 restores axonal transport in a neuronal prion disease model

Axonal vesicular trafficking is central to neuronal function. In addition to the formation of PrP^PG14^ endoggresomes, live cell imaging revealed the movement of small PrP^PG14^-mCh vesicles within axons of neurons expressing PrP^PG14^-mCh (Figure 7K; top panel).^167^ Semi-automated single-particle high-resolution quantitative image analysis^169^ showed impaired movement of PrP^PG14^-mCh (+ DMSO vehicle) vesicles 2 d post-transfection (Figure 7K; middle panel). After entering the axon, the majority of PrP^PG14^-mCh vesicles remained stationary compared to those moving in axons of neurons expressing PrP^WT^-mCh (+ DMSO vehicle) (Figure 7K, 7L, Supplemental Movies 1 and 2). The remaining motile PrP^PG14^-mCh vesicles also displayed defective movement compared to their PrP^WT^-mCh counterparts, as they spent less time in motion, and paused more frequently during runs (Figure 7K, 7L). Moreover, PrP^PG14^-mCh vesicles moved significantly slower and their movement was less processive, as indicated by decreased velocity and run length cumulative distribution functions (CDFs), respectively (Figure 7K, S5E). Notably, treatment of neurons with 200nM CCT020312 resulted in a significant rescue of various transport parameters to those resembling the transport of PrP^WT^-mCh vesicles (Figure 7K, 7L, S5E, and Supplemental Movie 3), including enhanced anterograde segmental velocities of PrP^PG14^-mCh vesicles beyond those observed for PrP^WT^-mCh vesicles (Figure S5E, and Supplementary Movies 1 and 2). Our data indicate that treatment with CCT020312 at nM concentrations reduces PrP^PG14^ endoggresomes formed within mammalian axons, thus normalizing the anterograde and retrograde axonal transport of smaller PrP^PG14^-mCh vesicles–vesicular axonal movement critical for the normal function of neurons.

## Discussion

Autophagy is an important pathway for the maintenance of a healthy proteome.^170^ This hypothesis is strongly supported by the number of neurodegenerative diseases in which impaired autophagy has been implicated as a driver of pathology, including AD, Parkinson’s disease, ALS and others.^171–173^ While therapies for these diseases are beginning to emerge,^174–178^ we envision that autophagy activators, like CCT020312, could become useful drugs (upon medicinal chemistry optimization) to be used in combination with current regulatory agency approved drugs, based on their demonstrated efficacy.

The utility of the LD screen for identifying modulators of autophagy was demonstrated herein. This HTS can be done in a variety of cell types without any genetic manipulation. The OA supplemented LD assay (improves sensitivity) was used to screen 940,000 small molecules, the largest number of compounds screened to date for identifying autophagy activators by over three-fold.^23,179^ At least 40 autophagy activators were identified by this effort, as demonstrated by their activity in multiple macroautophagy flux assays, including the GFP-LC3-RFP-LC3ΔG autophagy effector assay carried out in RPE1 cells. Numerous mTORC1-inhibitor independent autophagy activators were discovered by the LD HTS, exemplified by CCT020312. These are being developed to treat various degenerative diseases and will be published subsequently.

Our data indicate that CCT020312 induces the TFEB transcriptional program by triggering TFEB nuclear localization through a target to be identified. TFEB activation has been shown to induce autophagy in several cell types.^125,180^ This is consistent with our findings that CCT020312 can activate macroautophagy in all cell types explored so far, including HeLa and RPE1 cells, and a variety of human and mouse neurons. Despite different basal levels of macroautophagy flux in various cell lines tested, CCT020312 activated macroautophagy on the timescale expected of a transcriptional program in all of them. While we discovered that CCT020312 could promote TFEB nuclear localization, we did not detect changes in the phosphorylation level of TFEB at Ser211, nor of other targets that are directly modulated by mTORC1. These data suggest that CCT020312 induces TFEB nuclear localization through a mechanism distinct from mTORC1 inhibition.^181^ It has been reported that other TFEB phosphorylation sites, probably regulated by other kinases, may modulate its activity, and some of these are via mTOR-independent mechanisms of action.^182^ Thus the target of CCT020312 could be a kinase or phosphatase. The EM data from HeLa cells reveals that CCT020312 treatment mediates a statistically significant increase in the number of electron-dense DGCs including, autolysosomes and lysosomes, compared to vehicle treatment over time (Figure 3B and Figure S3) showing that treatment with CCT020312 drives the remodeling of the membrane architecture required for macroautophagy, consistent with macroautophagy mediated by the TFEB transcriptional program.

CCT020312 also activates autophagy in several types of neurons based on numerous lines of evidence. In human cortical neurons derived from induced pluripotent stem cell (iPSC) differentiation,^143,144^ DALGreen and DAPRed probes demonstrate that neurons treated with CCT020312 exhibited a dose-dependent increase in the number of autolysosomes forming relative to the number of autophagosomes forming; Figure 4C, 4D), consistent with the EM data summarized above in a cancer cell. LysoTracker Red increased fluorescence intensity suggests larger and/or greater number of lysosomes in CCT020312-treated neurons (Figure 4E and 4F). Together, these results demonstrate that CCT020312 treatment enhances macroautophagy flux in human iPSC-derived cortical neurons above a concentration of 50nM. In primary mouse cortical neurons transfected with LC3-Dendra2 and subsequently irradiated with 405 nm light to irreversibly convert Dendra2 emission from green to red fluorescence, CCT020312 treatment revealed a dose-dependent reduction in the mean half-life of LC3-Dendra2 upon CCT020312 treatment above a concentration of 5nM, consistent with increased macroautophagy flux (Figure 4B). In an iPSC-derived Tau-P301L mutation harboring mixed population of neurons, CCT020312 administration mediated autophagy activation based on dose-dependent increases in LC3-II and upregulation of additional autophagy and lysosomal markers including LAMP1, p62 and CTSD. In murine primary neurons transfected with mutant prion protein, we observed an overall increased acidity in endolysosomes of neurons treated with 200 nM CCT020312, consistent with autophagy activation. That CCT020312 is potent at increasing the macroautophagy flux in mouse primary cortical neurons suggests that it could be useful in mouse models of neurodegenerative diseases, although we realize its metabolic stability and blood brain barrier permeability may have to be improved with medicinal chemistry efforts. That CCT020312 also activates macroautophagy in directly differentiated from fibroblasts collected from older AD patients indicates that CCT020312 could be useful in treating neurodegenerative diseases in the elderly who have epigenetic marks of aging.

The two major hypotheses underpinning the etiology of AD are (1) that amyloid beta (Aβ) and tau aggregation drive pathology^183,184^ along with (2) endolysosomal/lysosomal system dysfunction mediated by increased amyloid precursor protein *C*-terminal fragment levels^26,32^ and decreased functional PSEN1,^33^ which compromise lysosome acidification and autophagic function.^26,62,185,186^ Microglial cell overactivation may also become a major driver of AD in some patients as the disease progresses.^187^ The human genetics of AD and the Lecanemab pharmacology showing a 26% slowing of AD progression demonstrate that clearance of Aβ aggregates slows AD progression.^174–178^ Very recently, Ionis/Biogen announced slowing of cognitive decline in early stage Alzheimer’s disease patients by CNS administration of an antisense oligonucleotide that lowers Tau levels, probably diminishing Tau aggregation. Patients with other tauopathies, such as frontotemporal dementia (FTD), may also benefit by reducing Tau aggregation, especially if used in combination with drugs that would improve endolysosomal system function. Herein, we show that the mTOR-independent autophagy activator CCT020312 reduces the cellular levels of Aβ and Tau. Interestingly, in human *ex vivo* FTD-derived neurons, CCT020312 reduced tau burden and toxicity (Figure 5), including oligomeric insoluble tau species, for up to 16 days, suggesting favorable kinetics for therapeutic development since pulsatile exposure may lead to long-term reduction of tau burden even after the majority of CCT020312 has been cleared from the system. This is analogous to previously published data with mTORC1 inhibitors, demonstrating that they can also have a prolonged pharmacodynamic effect.

Axonal inclusions in neurons of neurodegenerative disease patients are common;^159,167^ thus, our demonstration in Figure 7 that we can clear axonal membrane-delimited mutant prion protein aggregates (endoggresomes) by treatment with CCT020312 while normalizing vesicular trafficking is encouraging for treating neurodgenerative diseases.

The identification of CCT020312, which appears to activate macroautophagy through a TFEB mechanism, should be useful to cell biologists and physician scientists, especially those studying degenerative diseases. The development of compounds that directly or indirectly inhibit mTORC1 has been the primary pharmacological strategy for inducing autophagy to prolong healthspan/extend lifespan and to ameliorate proteinopathies. CCT020312 may be the best understood pharmacological means of activating macroautophagy by an mTORC1-inhibition independent mechanism.

## Author Contributions

Conceptualization, L.Y., R.C.B., A.V., S.J.H., K.A.J., M.C.S., A.S., S.E.E., J.W.K.; methodology, L.Y., R.C.B., A.V., Y.W., C.M.C., E.P.T., L.A.M., P.S.-M., C.-C.C., J.X., L.P.E., K.L., S.R.L., G.M.K., Q.X., S.C.-F., C.A.C., A.T., W.C.H., C.W.-G., L.H.I., K.N., Y.K., W.R., O.L.L., H.P., S.J. ; validation, L.Y., R.C.B., A.V., Y.W., C.M.C., E.P.T., L.A.M., P.S.-M., C.-C.C., J.X., L.P.E., K.L., S.R.L., G.M.K., Q.X., D.R., S.C.-F., A.T., K.N., W.R.,; investigation, L.Y., R.C.B., A.V., Y.W., C.M.C., E.P.T., L.A.M., P.S.-M., C.-C.C., J.X., L.P.E., K.L., S.R.L., G.M.K., Q.X., D.R., S.C.-F., L.H.I., K.N., Y.K., O.L.L.,; writing – original draft, L.Y., R.C.B., A.V., C.M.C., E.P.T., L.A.M., P.S.-M., C.-C.C., K.L., S.R.L., G.M.K., K.A.J., M.C.S., A.S., S.E.E.; writing – review & editing, L.Y. R.C.B., A.V., Y.W., C.M.C., E.P.T., L.A.M., C.-C.C., J.X., L.P.E., K.L., S.R.L., G.M.K., Q.X., C.A.C., W.C.H., M.J.B., M.H., M.A.P., I.D., F.R., D.G., R.L.W., E.T.P., S.J.H., K.A.J., M.C.S., A.S., S.E.E., J.W.K.; supervision, J.W.K., S.E.E., A.S., M.C.S., K.A.J., S.J.H., J.F., E.T.P., R.L.W., D.G., I.D., D.F., S.F., M.H., M.J.B., S.A.L.; funding acquisition, J.W.K., S.E.E., A.S., M.C.S., K.A.J., S.J.H., J.F., R.L.W., I.D., D.F., S.F., M.H., M.J.B., S.A.L.

## Acknowledgements

We acknowledge the expert assistance of Scott Henderson, Kimberly Vanderpool, and Theresa Fassel of The Core Microscopy Facility at The Scripps Research Institute for transmission electron microscopy data. We acknowledge Anan Yu, Susan Fox and Richard I. Morimoto for helpful conversations and critical comments; Rebeccah Ryden and Kylee House for administrative assistance. L.Y. would like to acknowledge Julia M. Grandjean and Jessica D. Rosarda for training in qPCR, RNA-Seq data analysis, and the CellTiter Glo assay; Zainab Ahmed for assistance with RNA-Seq sample prep and TFEB nuclear localization immunofluorescence assay assistance; Maziar S. Ardejani for assistance with RNA-Seq data analysis; Chongyang Wu and Ivan Putra for harvesting neurons for macroautophagy flux assays; Alina Meneses and Luke T. Nelson for assistance with LC3 western blots; Brittany Sanchez, Quynh Nguyen and Jason S. Chen in the Automated Synthesis Facility at The Scripps Research Institute for compound purification and helpful conversations; Gabriel J. Brighty, Carlos Cosme Jr, Jolene K. Diedrich, Julia Jones, Chung-Yon Lin, Ryan J. Paxman, and Justin Wang for helpful conversations; and Marco Mravic for providing critical comments. R.C.B. acknowledges J. Wade Harper and Heeseon An for sharing valuable cell lines and expertise in autophagy. E. P. T. and M. H. acknowledge Nora Lyang for assistance with TFEB nuclear translocation immunofluorescence assay. Emily P. Bentley contributed to the writing and editing of this manuscript.

J.W.K. acknowledges financial support from RF1AG073418 and the Rainwater Charitable Foundation for the discovery of autophagy activators to ameliorate Alzheimer’s disease and related dementias. S.E.E. was supported by NIH/NIA R01AG049483 and R01AG076745 grants, and by NIH/NIA/NINDS R56NS131648 grant. This work was supported by NIA P01AG054407 (J.F., D.F. & J.W.K.) and the Michael J. Fox Foundation for Parkinson’s Research and the Aligning Science Across Parkinson’s (ASAP) initiative (J.F.). The Michael J Fox Foundation administers the grant ASAP-000282 on behalf of ASAP and itself. C.-C.C. was supported by the Glenn Foundation for Medical Research Postdoctoral Fellowship in Aging Research, the Life Sciences Research Foundation Postdoctoral Fellowship and the Alzheimer’s Disease Research program in BrightFocus Foundation Grant. A.V. was supported by a George E. Hewitt Foundation for Medical Research Postdoctoral Fellowship, and Y.W. was supported by Dorris Neuroscience Center Fellowship Awards. S.J.H and M.C.S. were supported by the Rainwater Charitable Foundation and from the Stuart and Suzanne Steele MGH Research Scholars Program. A.S. was funded by the Deutsche Forschungsgemeinschaft (DFG, German Research Foundation) – 259130777 (SFB1177); 515275293 and the Dr. Rolf M. Schwiete Stiftung (13/2017). I.D. was funded by the Deutsche Forschungsgemeinschaft (DFG, German Research Foundation) – 25913077 (SFB1177). F.R. is supported by a Novo Nordisk Foundation (0066384) grant. S.A.L. was supported in part by NIH grants R35AG071734, R01AG056259, R01DA048882, RF1AG057409, and R56AG065372. M. H. was funded by R01AG038664.

## Materials and Methods

### LD screen and initial macroautophagy flux assessments

#### Cell culture

SW1990, HEK293T, and HeLa cells were purchased from ATCC. HT-1080 cells were purchased from Sigma-Aldrich. U251 cells were a generous gift from Calibr. HeLa Difluor (RFP-GFP-LC3) cells were purchased from Invivogen. Cells were cultured at 37°C and 5% CO_2_ in DMEM, RPMI, or MEM supplemented with 10% fetal bovine serum (FBS) and penicillin-streptomycin, as recommended by vendor.

For DALGreen/DAPRed and LysoTracker studies, human induced pluripotent stem cell (iPSC)-derived neural progenitor cells (NPCs) corresponding to the WT/WT parental line 7889SA (NYSCF Repository) were acquired from the laboratory of Stuart Lipton. The hiPSCs were differentiated following previously published standard protocols.^143,144^ In brief, hiPSCs were induced via addition of an inhibitors cocktail containing Dorsomorphin, A83-01, and PNU74654 in DMEM/F12 medium with 20% knock out serum replacement (Invitrogen) for 6 d. Cells were then manually scraped to form PAX6+ neurospheres maintained in DMEM/F12 medium supplemented with N2 and B27 (Invitrogen) and 20 ng/ml of bFGF for 2 weeks. The neurospheres were then seeded onto p-ornithine/laminin-coated 6-well plates to form a monolayer of neural progenitor cells (NPCs) in neural medium supplemented with 20 ng/ml bFGF to mature, expand, and bank. Terminal NPC differentiation into cortical neurons was done by dissociating mature NPCs into 0.1% Polyethyleneimine (PEI)/4 µg/ml Laminin coated coverslips at a density of 150,000-400,000 NPCs/coverslip. The cells were maintained in neural medium supplemented first with 0.1µM Compound-E for 2 d, and thereafter with 20 ng/ml each of hBDNF and hGDNF. Finally, the basal medium was switched to BrainPhys at week 3 with the same hBDNF and hGDNF supplements. Experiments were conducted between weeks 5-7 of terminal differentiation and maturation. To confirm the quality of the differentiated neurons, they were stained for immunohistochemistry with anti-PAX6 and anti-NeuN antibodies (looking for the absence of the latter and the presence of the former, the mature neuronal marker). In the Haggarty lab, work with human iPSC and the derived NPC lines was executed under the Massachusetts General Hospital/Mass General Brigham-approved IRB Protocol #2010P001611/MGH. Patient-specific iPSCs and cortical-enriched NPCs were generated as previously described.^153,188^ In the mutant tau study, the NPC line used is designated MGH2046-RC1 and is derived from an individual in her 50s with FTD and carrier of the tau-P301L autosomal dominant mutation (c.C1907T, rs63751273).^31,188^ NPCs were cultured in 6-well (Fisher Scientific Corning) or black 96-well clear bottom (Fisher Scientific Corning) plates coated with poly-ornithine (20 μg/mL in water, Sigma) and laminin (5 μg/mL in PBS, Sigma), referenced to as POL-coated plates. NPC proliferation medium composition was as follows: 70% (v/v) DMEM (Gibco), 30% Ham’s-F12 (v/v) (Fisher Scientific Corning), 2% B27 (v/v) (Gibco), and 1% penicillin-streptomycin (v/v) (Gibco); freshly supplemented with 20 ng/mL EGF (Sigma), 20 ng/ml FGF (Stemgent) and 5 μg/ml heparin (Sigma) at the time of cell passaging, which was done using TrypLE reagent (Thermo Fisher).

Primary hippocampal neuron cultures were prepared as described previously.^189^ Briefly, hippocampi were dissected from 1 to 2-d-old mouse neonates in cold Hank’s buffer solution (HBSS) (Gibco), supplemented with 0.08% D-glucose (Sigma), 0.17% HEPES, and 1% Pen-Strep (Gibco), filter sterilized and adjusted to pH 7.3. Dissected hippocampi were washed twice with cold HBSS (Gibco) and individually incubated at 37°C for 15 to 20 min in a sterile solution of 45 U papain (Worthington), 0.01% DNase (Sigma), 1 mg DL-cysteine (Sigma), 1 mg BSA (Sigma), and 25 mg D-Glucose (Sigma) in PBS (Gibco). Hippocampi were washed twice with Dulbecco’s Modified Eagle Medium (DMEM) (Gibco), supplemented with 10% FBS (Gibco) (pre-heated to 37 °C), and disrupted by ten to twelve cycles of aspiration through a micropipette tip. Dissociated neurons were then resuspended in warm DMEM (Gibco) + 10% FBS (Gibco) and plated in 24-well plates containing 12-mm glass coverslips pretreated with 50 µg/ml poly-L-lysine (Sigma) in Borate buffer (1.24 g boric acid (Fisher), 1.90 g borax (Sigma), in 500 ml cell-culture grade water, adjusted to pH 8.5, filter sterilized). After 1 h, media was replaced with Neurobasal-A medium (NBA) (Gibco), supplemented with 2% B-27 (Gibco) and 0.25% GlutaMAX (Gibco).

For Aβ studies, human dermal fibroblasts were acquired from Stanford Alzheimer’s Disease Research Center (ADRC). Cells were cultured in DMEM medium with high glucose and GlutaMAX containing 10% FBS, 1% Penicillin-Streptomycin, MEM Non-Essential Amino Acids, 1% sodium pyruvate, and 0.1% beta-mercaptoethanol (ThermoFisher Scientific).

#### Semiautomated high-throughput LD clearance screen

SW1990 cells (350 per well) were plated in 8 µL of 10µM OA-supplemented phenol-red– free Roswell Park Memorial Institute (RPMI) complete growth media (Gibco) into black-walled clear-bottom 1536-well plates. The OA and BSA were purchased from Sigma and the complex was prepared using a published protocol,^190^ or the OA-BSA complex was purchased from Sigma. These cells were allowed to grow and adhere for 20 h. Cells were then treated individually with 4µM LOPAC library component compounds (https://www.sigmaaldrich.com/US/en/product/sigma/lo1280) or individually with 5µM screening library compounds (940,000 small molecules in total from Calibr, most of which are commercially available or have been publicly disclosed.) for 20 h. Cells were fixed and stained for at least 1 h at RT (≈ 25°C) with a 2µL addition of a 5X fixative/staining solution (comprising final concentrations of 4% paraformaldehyde (PFA), 1µM BODIPY 493/503, and 0.1 µg/ml Hoechst 33342 fluorophore). Images were collected with a 10X objective on a CellInsight CX5 high-content imager employing an LED solid-state light source and filter sets of excitation 485/emission 521 (BOPIDY 493/503) and ex 386/em 440 (Hoescht) and processed using the Spot Identification algorithm in Cellomics. The entire area of each well was visualized in a single image. Columns of cells were treated with the vehicle control DMSO (final concentration of DMSO in each well was <0.15%) and tamoxifen (15 µM) in DMSO was employed as the positive control. The threshold for primary hit compounds was a >50% reduction in total lipid area in comparison to vehicle-treated cells, with a ≤30% reduction in cell number. Cell number measurements were determined by counting the nuclei stained with Hoescht in a given image (∼230 nuclei were detected in vehicle control wells). All hit compounds were retested in triplicate at 5 µM and the same parameters for advancement were applied (>50% reduction in total lipid area, with ≤30% reduction in cell number). Confirmed hits were assessed for dose-dependent activity (10-point dilution ranging from 5µM to 9nM). Only compounds demonstrating dose-dependent reduction in cellular lipid load, in addition to a >50% reduction in total lipid area, with ≤30% reduction in cell number at maximum concentration, were progressed. The primary hit rates were 0.65% for the 940,000 HTS compound collection and 0.98% for the LOPAC constituents.

#### Counter screen for fluorescent artifacts and elimination undesirable chemical moieties

The 409 confirmed hits exhibiting dose-dependent activity as well as the other attributes described above for progression were subjected to counter screening. SW1990 cells (350 per well) were plated in black-walled clear-bottom 1536-well plates in 10µM OA supplemented phenol-red–free RPMI complete growth media and allowed to adhere for 40-45 h (total incubation period with lipid in primary screen). Cells were fixed via 2 µL addition of a 5X fixative/staining solution (final concentrations of 4% PFA, 1µM BODIPY 493/503, 0.1 µg/mL Hoechst 33342) for 2 h at RT (≈ 25°C). Cells were then treated individually with primary hit compounds at a concentration of 5µM for 1 h. A column of cells was treated with DMSO vehicle control, while a column of cells was fixed and stained (4% PFA, 0.1 µg/ml Hoechst 33342) without a lipid dye (i.e. no BODIPY 493/503) as a positive control. Images were collected with a 10X objective on a CellInsight CX5 high-content imager and processed using the same Spot Identification algorithm in Cellomics using Spot Total Area per Cell as previously described. All hits for which the residual activity (BODIPY 493/503 fluorescence area decrease in live cells during treatment minus artifactual BODIPY 493/503 fluorescence area decrease in fixed cells) was not >50% were eliminated. In total, 188 compounds were eliminated, leaving 221 compounds. These 221 compounds were manually curated to eliminate pan-assay interference compounds, those bearing significant metabolic liabilities, and compounds that were frequent hits in a wide variety of high-throughput screens, leaving 108 hits, of which 81 could be reordered as solid stocks.

#### Confirmation of re-ordered hits

SW1990, U251, and HT-1080 cells (1000 cells / well) were plated in 50 µL of 10µM OA supplemented RPMI complete growth media (Gibco) into black-walled clear-bottom 384-well plates. The OA and BSA were purchased from Sigma and the complex was prepared using a published protocol,^190^ or the OA-BSA complex was purchased from Sigma. These cells were allowed to grow and adhere for 20 h. Cells were then subjected to 10-point dose response curve (10µM to 0.01nM) of the aforementioned 81 hits for 20 h. Cells were fixed and stained via 2 µL addition of a 5X fixative/staining solution (final concentrations of 4% PFA, 1µM BODIPY 493/503, 0.1 µg/ml Hoechst 33342) for at least 1 h at RT. Plates were washed with DPBS using an automated Biotek automated plate washer. Cell images were collected with a 10X objective on a CellInsight CX5 high content imager, and LED solid-state light source and filter sets of ex 485/em 521 (BOPIDY 493/503) and ex 386/em 440 (Hoescht) and processed using the Spot Identification algorithm in Cellomics using Spot Total Area per Cell. Cellular lipid area was determined relative to lipid load of vehicle-treated cells. Compounds confirmation required a dose-dependent decrease in cellular lipid load as ascertained by a decrease in BODIPY 493/503 fluorescence.

#### Immunoblot assessment of autophagy

Adherent cells at 75% confluency (150,000 to 300,000 cells) in six-well plates were treated with small molecules for 20 h. (For data in Figure S1A, SW1990 cells were plated in RPMI containing 10% FBS, 1% PenStrep, and 10µM OA, and the next day compounds were spiked into this unchanged media.) At the conclusion of treatment, the medium and trypsin-aided detached cells were pelleted, washed with PBS, and lysed on ice in RIPA buffer for 30 min with intermittent vortexing (50mM Tris base, 150mM NaCl, 1% Triton X-100, 0.5% sodium deoxycholate, 0.1% SDS, pH 7.4, with a 1:100 dilution of Halt Protease Inhibitor EDTA-free, 1:100 dilution of Halt Phosphatase Inhibitor, 0.5 µL/mL benzonase). Lysates were clarified (10 min at 14,000 rpm, 4°C). Protein content was normalized by BCA Protein Assay reagent (Pierce). Samples were denatured, separated by SDS-PAGE (4−20%), and transferred to a PVDF membrane (Biorad Trans Turbo). Membranes were blocked with 5% milk in TBST, probed with primary antibodies LC3-I/II (Cell Signaling 2775), phospho S6K (Cell Signaling 9234), S6K (Cell Signaling 2708), and GAPDH (Cell Signaling 2118) overnight at 4°C in TBST containing 5% BSA, washed with TBST (two fast washes, two 10-min washes on gyrotary shaker) then imaged by chemiluminescence with a HRP-conjugated secondary antibody (Cell Signaling 7074).

#### GFP-LC3-RFP-LC3ΔG autophagy flux reporter assay

pMRX-IP-GFP-LC3-RFP-LC3ΔG was a gift from Noboru Mizushima (The University of Tokyo, Tokyo, Japan; Addgene plasmid #84572; https://www.addgene.org/84572/). 2000 RPE1 cells per well were seeded in 384-well plates in 50 µL DMEM media, supplemented with 10% FBS, 1% penicillin/streptomycin, and incubated for 24 h before treatment. 50 μL of media containing either 2x final concentration of indicated compounds in or control compounds (final conc.: 0.1% DMSO, 250 nM Torin-1, 200 ng/ml Bafilomycin A1) were added and images in three channels (fluorescence of GFP, RFP, as well as phase) were taken every 2 h over a period of 72 h using the IncuCyte® (Sartorius, Germany). General macroautophagy flux was calculated via the ratio of the total integrated fluorescence intensities of GFP/RFP. Cell confluence is represented as % of covered area by cells. Each data point represents the averaged ratio or confluence obtained from three individual wells of the plate.

To visualize the complete dataset of the screen, values for each individual compound and time point were normalized to the respective value of the negative control DMSO by subtracting the averaged ratio from the corresponding average ratio of the averaged DMSO control of the same 384-well plate. The resulting normalized data was put together sequentially and represented in a heatmap (Figure 1E). Data distribution was analyzed, and thresholds were selected by analyzing the data’s symmetry to represent the whole range within the dataset (red = higher autophagy flux than DMSO; green = lower autophagy flux than DMSO). Values of 20% and 80% within the data’s histogram with a significant frequency (outliers removed) were chosen as the maximum and minimum of the heatmaps’ range, respectively. In a second heatmap (Figure 1F), the slope of flux kinetics (direction of changes of the autophagy flux between time points) was presented. Each subsequent set of two timepoints was averaged to represent a meaningful trend. Data distribution was evaluated, and thresholds were selected to represent the whole range of the dataset as described above (red = lower slope than DMSO = higher autophagy flux induction; green = steeper slope than DMSO = lower autophagy flux). Hits were defined as compound reaching 20% or more of the positive control (Torin1) autophagy flux for three or more consecutive timepoints in any of the tested concentrations.

#### LC3B and LAMP2 immunofluorescence confocal microscopy

HeLa cells (2.0×10^4^, 400 µL) were seeded in 8-well Lab-Tek chambered coverglass (Thermo #155409) and treated with 0.1% DMSO, 250nM rapamycin (Alfa Aesar #J62473), 1µM CCT020312, or 2.5µM CCT020312 for 20 h. After treatment, cells were fixed and permeabilized with ice-cold methanol for 10 min in −20°C freezer. Cells were then washed three times with PBS and incubated with a rabbit primary antibody against LC3B (CST #3868) diluted 1:100 in PBS containing 10% FBS and 0.2% saponin overnight at 4°C. Then, LC3B primary antibody was removed and mouse antibody against LAMP2 (Abcam #H4B4) diluted 1:200 in PBS was directly added to the cells for 12 h at 4°C. After the removal of LAMP2 primary antibody, the cells were washed 3 times with PBS and a mixture of goat anti-mouse IgG(H+L) cross-absorbed secondary antibody conjugated to Alexa Fluor^TM^ 488 (Invitrogen #A-11001) and goat anti-rabbit IgG(H+L) cross-absorbed secondary antibody conjugated to Alexa Fluor^TM^ 546 (Invitrogen #A-11035), both diluted 1:500 in PBS, was added to the cells for 2 h at RT. After removal of secondary antibodies, cells were incubated with DAPI (1 µg/ml in PBS) for 5 min at RT, followed by three washes with PBS. After the washes, 200 µL PBS was added to each well and the chamber slides were subjected to imaging. Zeiss Confocal 780 was used for imaging with 63x oil objective lens, where laser lines 488 nm, 561 nm, and 405 nm were used for the excitation of Alexa Fluor^TM^ 488, Alexa Fluor^TM^ 546, and DAPI fluorophores. Images collected were further processed by ImageJ to overlap each channel to give the merged images. Colocalization of LAMP2 and LC3B was represented by Pearson’s Coefficient (1 indicates complete positive correlation and 0 indicates no correlation), which is derived from JACoP plugin in Fiji.^191^

#### Autophagy flux assay in primary mouse cortical neurons

pGW1-LC3-Dendra2 was transfected into cultured primary mouse cortical neurons. Turnover of LC3-Dendra2 was performed by optical pulse labeling. Dendra2 is a photoswitchable protein, which has been described previously.^141,192^ 2 h prior to photoswitching, compounds were added to the neuronal cultures (final concentration: 250nM rapamycin, 0.5nM, 5nM, 50nM, 500nM CCT020312 or carboxy CCT or equivalent volume of DMSO). The decay of photoswitched Dendra2 fluorescence was measured with the same microscope by following photoswitched neurons at approximately 4-h intervals over 24 h and quantitated as previously described.^141,192^

#### Human iPSC-derived neurons culture, treatment and autophagy flux

NPCs from Stuart Lipton’s laboratory were terminally differentiated into cerebrocortical neurons following a standard protocol.^143^ In brief, 300,000 to 400,000 NPCs seeded into microscope coverslips were treated with 100nM compound E (EMD Millipore, Temecula, CA) in BrainPhys medium (StemCell Technologies) for 48 h and then maintained thereafter in BrainPhys medium supplemented with BDNF and GDNF. Upon 7 weeks of differentiation and maturation, quality control histological stainings against NeuN and Ca^2+^ imaging neural activity measurements were done to confirm robust mature neuronal signatures before beginning treatment experiments. The small molecules (250nM rapamycin, 50nM CCT020312, 200nM CCT020312, or vehicle control) were added for 24 h with DMSO normalized in all conditions to 1% (v/v). Cells were treated with 75nM LysoTracker Red (Thermo) for final 3 h of treatment or with 1µM DALGreen (Dojindo D675) and 200nM DAPRed (Dojindo D677) according to manufacturer’s instructions (https://www.dojindo.com/products/manual/D677e.pdf) for final hour of treatment. For the final 30 min, Hoechst dye was added as nuclear stain. A full media change was performed at the 24-h time point after washing right before imaging in a confocal fluorescence microscope (Nikon C2) at a 10x magnification. 3 to 5 different fields were randomly selected per coverslip replicate. Images were processed and analyzed with ImageJ. DALGreen-DAPRed colocalization, background subtraction and thresholding were used to define a population of Green/Red puncta, which the resulting intensity ratio was calculated and used to measure the macroautophagy flux (modified from manufacturer’s protocol).

#### Assessment of the autophagy flux by transmission electron microscopy

700,000 HeLa Cells were plated in Nunc^TM^ Permanox^TM^ (Cat #174888) petri dishes, treated the next day (for 4, 11, 15.5 or 24h) with fresh DMEM (supplemented with 10% FBS and 1% PenStrep) containing 0.1% DMSO (vehicle control), 2.5µM CCT020312, 250nM Torin-1, 100nM Bafilomycin A1 or left untreated with only a media change and washed with serum and antibiotic-free DMEM and fixed in oxygenated 4% PFA + 2.5% glutaraldehyde^193^ in 0.1M sodium cacodylate buffer (pH 7.3) for 5 min at 37°C, then transferred to ice and fixed for 1 h. Cells were washed with buffer, and post-fixed in buffered 1% osmium tetroxide + 1.5% potassium ferrocyanide for 1 h on ice. The samples were stained en bloc with aqueous 0.5% uranyl acetate overnight at 4°C. Dehydration was carried out using a graded ethanol series followed by acetone. The cells were embedded in LX-112 (Ladd Cat #21210) epoxy resin and polymerized at 60°C. Ultrathin sections (silver) were cut using a Leica UC7 ultramicrotome and imaged at 80 kV with a ThermoFisher Talos L120C transmission electron microscope and images were acquired with a CETA 16-M CMOS camera. The average number of DGCs (amphisomes, lysosomes and autolysosomes) per cell section was determined by counting these compartments through 50 cell sections per condition, randomly selected from two biological replicate experiments.

#### Measurement of UPR and HSF1 activity using luciferase reporters

HEK293T-Rex cells expressing the ATF4.FLuc, ERSE.FLuc,^194^ XBP1s.RLuc,^195^ or HSF1.FLuc reporter were plated at 80 µL/well from suspensions of 160,000 cells/mL (XBP1s.RLuc and HSF1.FLuc) or 170,000 cells/mL (ATF4.FLuc) or 100 µL/well from suspensions of 200,000 cells/mL (ERSE.FLuc) in white clear-bottom 96-well plates (Corning) and incubated at 37°C overnight. The following day, cells were treated with 20 µL (ATF4.FLuc, XBP1s.RLuc and HSF1.FLuc) or 25 µL (ERSE.FLuc) of compound-containing media to give final concentration as described before incubating for 18 h at 37°C. The plates were equilibrated to RT, then either 100 μL Firefly luciferase assay reagent-1 (ATF4.FLuc and HSF1.FLuc), 125 µL Firefly luciferase reagent-1 (ERSE.FLuc) or 100 µL Renilla luciferase assay reagent-1 (XBP1s.RLuc) (Targeting Systems) were added to each well. Samples were dark adapted for 10 min to stabilize signals. Luminescence was then measured in an Infinite F200 PRO plate reader (Tecan) and corrected for background signal (integration time 500 ms). All measurements were performed in biological triplicate.

#### ARE-LUC reporter assay

IMR32 cells were plated at 5000 cells per well in white, clear bottom 384-well plates in 40 µL growth medium. The next day, 100 ng pTI-ARE-LUC reporter plasmid in 10 µL OptiMEM (Gibco) was transected into each well using Fugene HD at a dilution of 1 µg plasmid DNA:4 µL Fugene. After 24 h, compounds were transferred using a 100 nL pintool head affixed to PerkinElmer FX instrument. After 24 h of compound incubation, 30 µL BrightGlo reagent solution (Promega) diluted 1:3 in water was added to each well and then ARE-LUC luminescence values were recorded on an Envision instrument (PerkinElmer).

#### GSH/GSSG-Glo^TM^ assay

HEK293T cells were plated in sextuplet per planned treatment condition at 10,000 cells per well in white, clear bottom 96-well plates (Corning Cat #3903) in 100 µL DMEM (Gibco) with 10% FBS and 1% Pen-Strep. The next day, the media was replaced with fresh media containing 0.1% DMSO, 40µM positive control radical oxidative stress inducer menadione or CCT020312 (100nM, 1µM, 5µM or 10µM). After 2 h of compound incubation, 50 µL Total Glutathione Lysis Reagent (Promega Cat #TM344) was added to 3 of the wells per treatment condition and 50 µL Oxidized Glutathione Lysis Reagent (Promega Cat #TM344) was added to the other 3 wells per treatment condition. 50 µL of the two Lysis Reagent buffers were also added to 2 empty wells per buffer not containing any cells (background control). The plates were shaken at RT for 5 min. After shaking, 50 µL Luciferin Generation Reagent (Promega Cat #TM344) was added to all wells. Plates were shaken for 30 s at RT followed by 30-min RT incubation with no shaking in the dark. Lastly, 100 µL Luciferin Detection Reagent (Promega Cat #TM344) was added to all wells, plates were shaken for 30 s at RT and incubated at RT for 15 min in the dark with no shaking. Then luminescence values were recorded on a Pherastar plate reader (BMG). The ratio of GSH/GSSG was calculated (“Calculating the Ratio from Net RLU”) and background correction was performed according to the manufacturer’s protocol.

#### RNA-seq analysis

HeLa or HEK293T cells were lysed and total RNA collected using the Quick-RNA Miniprep kit from Zymo Research (R1055) according to manufacturer’s instructions. RNA concentration was then quantified by NanoDrop. Whole transcriptome RNA was then prepared and sequenced by BGI Americas on the BGI Proprietary platform, which provided paired-end 50 bp reads at 20 million reads per sample. Each condition was performed in triplicate. RNA-seq reads were aligned using DNAstar Lasergene SeqManPro to the Homo_sapiens-GRCh38.p7 human genome reference assembly, and assembly data were imported into ArrayStar 12.2 with QSeq (DNAStar Inc.) to quantify the gene expression levels and normalization to reads per kilobase per million. Differential expression analysis was assessed using DESeq2 in R, which also calculated statistical significance calculations of treated cells compared to vehicle-treated cells using a standard negative binomial fit of the reads per kilobase per million data to generate fold-change quantifications.

#### Analysis with DAVID bioinformatics tool

Using the significant genes (adjusted p-value < 0.05) from RNA-seq analysis with DESeq2, the genes with fold change > 1 (versus DMSO) served as input for the DAVID bioinformatics tool. GO terms for Molecular Function, Biological Process and Cellular Component were sorted by lowest adjusted p-value (DAVID false discovery rate), then by highest-to-lowest fold enrichment (compared to background genome), and the top 4 terms from each category were plotted by fold enrichment in Figure 3F.

#### TFEB immunofluorescence confocal microscopy for measurement of nuclear localization

HeLa cells (2.5×10^5^) were seeded in 24-well plates containing 12-mm glass coverslips and treated with 0.1% DMSO or 1µM CCT020312 for 20 h. After treatment, cells were fixed with 4% PFA in PBS for 20 min at RT for TFEB staining. Fixed cells were first permeabilized using 0.1% Triton X-100 in PBS for 5 min at RT. Then, cells were incubated with a primary antibody against TFEB (CST #4240)^196^ diluted in PBS containing 10% FBS and 0.2% saponin for 2 h at RT. Stained cells were then washed three times with PBS and incubated with Alexa Fluor 488–labeled secondary antibodies (Thermo Fisher) diluted at 1:1000 in 10% FBS and 0.2% saponin in PBS for 1 h at RT. Cells were washed three times in PBS, incubated with 4′,6-diamidino-2-phenylindole (DAPI) to label nuclei, and mounted with Prolong® Gold Antifade Reagent. Imaging was performed using a Zeiss ApoTome. For each condition, 15 to 20 images were acquired in randomly selected fields with a 63X oil objective lens. An average of 10 to 15 stacks of 0.3 µm thickness were acquired.

### Functional rescue of prionopathy disease model

#### Primary neuronal culture

Primary hippocampal neuron cultures were prepared as described previously.^189^ Briefly, hippocampi were dissected from 1-to 2-d-old mouse neonates in cold Hank’s buffer solution (HBSS) (Gibco) supplemented with 0.08% D-glucose (Sigma), 0.17% HEPES, and 1% Pen-Strep (Gibco); filter sterilized; and adjusted to pH 7.3. Dissected hippocampi were washed twice with cold HBSS (Gibco) and individually incubated at 37 °C for 15 to 20 min in a sterile solution of 45 U papain (Worthington), 0.01% DNase (Sigma), 1 mg DL-cysteine (Sigma), 1 mg BSA (Sigma), and 25 mg D-Glucose (Sigma) in PBS (Gibco). Hippocampi were washed twice with Dulbecco’s Modified Eagle Medium (DMEM) (Gibco) supplemented with 10% FBS (Gibco) (preheated to 37 °C) and disrupted by ten to twelve cycles of aspiration through a micropipette tip. Dissociated neurons were then resuspended in warm DMEM (Gibco) + 10% FBS (Gibco) and plated in 24-well plates containing 12-mm glass coverslips pretreated with 50 µg/ml poly-L-lysine (Sigma) in Borate buffer (1.24 g boric acid [Fisher], 1.90 g borax [Sigma] in 500 mL cell-culture grade water, adjusted to pH 8.5, filter sterilized). After 1 h, media was replaced with Neurobasal-A medium (NBA) (Gibco) supplemented with 2% B-27 (Gibco) and 0.25% GlutaMAX (Gibco).

#### Prion Degradation Assay

At 7 d in vitro (DIV), hippocampal neurons grown on a 12-mm coverslip in 24-well plates were transfected using 2 µL of Lipofectamine and 0.8 µg of MoPrP^WT^- or MoPrP^PG14^-mCherry DNA per well. 24 h after transfection, small molecule CCT020312, carboxy CCT, rapamycin or vehicle was applied at a final concentration as indicated in Figures 7B, 7C, S7B, S7D and incubated for 2 h, after which the media was replaced with fresh Neurobasal-A medium (Gibco) supplemented with 2% B-27 (Gibco) and 0.25% GlutaMAX (Gibco). At 9 DIV, 48 h post transfection, neurons were fixed with 4% PFA (Electron Microscopy Services) containing 4% sucrose (Sigma) for 30 min at 37°C. Cells were washed once in 50mM Glycine in PBS, 3 more times in PBS, and once in H_2_O before being mounted in ProLong Diamond antifade reagent (Thermo Fisher). Fixed cells were imaged using a Nikon Ti-E Perfect Focus inverted microscope equipped with a total internal reflection fluorescence microscopy (TIRFM) setup, with an Andor iXon + DU897 EM camera, and a 100×/1.49 numerical aperture (NA) oil objective. Solid-state 561-nm lasers were used for detecting mCh. For all images acquired, the exposure time was set to 100 ms. Pixel size was 0.16 µm. Images were taken at 3 field of view away from the cell body. Transfection rates were ∼1%, which enabled imaging of individual neurons.

#### Image quantification of particle density in axons

For PrP^PG14^ aggregate density quantification from still images, we calculated the average axonal background greyscale value. We defined aggregates as those puncta with a greyscale value of at least 2 times the background. Axon length was measured using the polyline tool of ImageJ. Aggregate density was expressed as the average number of aggregates per 100 µm of axon length.

#### Axonal endolysosomal pH assessment by FIRE-pHLy

For characterizing the effect of CCT020312 on modulating lysosomal acidification in axons, hippocampal neurons were transfected with pFUGW-FIRE-pHLy with or without co-transfection of PrP^PG14^-mTagBFP2. One d after transfection, neurons were treated with DMSO or 200 nM CCT020312 for 2 h. Two d after transfection, live imaging was performed using a Nikon Ti-E Perfect Focus inverted microscope equipped with a total internal reflection fluorescence microscopy (TIRFM) setup, with an Andor iXon + DU897 EM camera, and a 100×/1.49 numerical aperture (NA) oil objective. Epifluorescence excitation was provided by a metal halide–doped mercury arc lamp (Nikon Intensilight). mTagBFP2, mTFP1 and mCh were collected using UV-2E/C [AT350/50x, T400lp, ET460/50m], ET-GFP [ET470/40x, T495lpxr, ET525/50m], and ET-DsRed [ET545/30x, T570lp, ET620/60m] Chroma filters, respectively.

For measuring acidity in PrP^PG14^ endolysosomes in axons, hippocampal neurons were co-transfected with pFUGW-FIRE-pHLy and PrP^PG14^-mTagBFP2. 1 d after transfection, neurons were treated with DMSO or 200 nM of CCT020312 for 2 h. 2 d after transfection, neurons were fixed with 4% PFA (Electron Microscopy Services) containing 4% sucrose (Sigma-Aldrich) for 30 min at 37°C, followed by three washes with PBS. Cells were then permeabilized by incubating the coverslips in 0.1% Triton X-100 (Sigma-Aldrich) in PBS for 5 min followed by three washes with PBS. Coverslips were incubated in blocking solution [3% immunoglobulin G (IgG)-free BSA and 10% donkey serum in PBS; Jackson ImmunoResearch] for 30 min at RT, followed by incubation with primary antibody against PrP (mouse anti-PrP^Sc^ (3F4), Novus, 1:250) in blocking solution overnight at 4°C. After three washes in PBS, neurons were incubated with secondary antibodies in blocking solution for 1 h at RT. After three washes in PBS, coverslips were mounted in ProLong Diamond antifade reagent (Thermo Fisher Scientific). Coverslips were imaged using the aforementioned Nikon Ti-E Perfect Focus inverted microscope. mTFP1, mCh and PrP labelled with Alexa 647 were collected using ET-GFP [ET470/40x, T495lpxr, ET525/50m], ET-DsRed [ET545/30x, T570lp, ET620/60m], and Cy5 HYQ [ET620/60x, T660lp, ET700/75m] Chroma filters, respectively.

All images were processed in ImageJ. ROIs of axons were made from each image using the polyline tool of ImageJ. Rolling-ball background fluorescence correction was applied to all ROIs. To quantify FIRE-pHLy fluorescence, the mCh channel was used to create an object “mask” for endolysosomes or endoggresomes. The “mask” was applied to both mTFP1 and mCh channels, and the mean fluorescence intensities of each lysosome in the mCh and mTFP1 channels were generated, and mTFP1/mCh intensity ratio was calculated. PrP endolysosomes and endoggresomes were identified with the 3F4 (Alexa 647) channel using the polyline tool, and mTFP1/mCh intensity ratio was calculated for LAMP1-positive PrP endolysosomes and endoggresomes.

The data were first tested for normality using the Prism D’Agostino-Pearson normality test, and were identified as nonnormally distributed. Mann-Whitney test was performed, and box plot showing median, interquartile range and all data points was plotted.

#### Live cell imaging for axonal transport measurements

Live imaging of axons of hippocampal neurons was done at d 9, 2 d post transfection. Coverslips (12 mm) were transferred and flipped onto 35-mm glass-bottom dishes (MatTek) containing 2 ml of Neurobasal-A medium (Gibco), containing 2% B-27 (Gibco) and 0.25% GlutaMAX (Gibco) medium. Neurons were imaged using a Nikon Ti-E Perfect Focus inverted microscope equipped with a total internal reflection fluorescence microscopy (TIRFM) setup, with an Andor iXon + DU897 EM camera, and a 100×/1.49 numerical aperture (NA) oil objective. Solid-state 561-nm lasers were used for detecting mCh. Lasers were positioned at varying angles for pseudo-TIRFM (pTIRFM) acquisition. Transfection rates were ∼1%, which enabled imaging of individual neurons.

Time-lapse movies of fast vesicular transport in axons were 15 s long and collected at a frame rate of 10 frames/s (10 Hz). For all movies acquired, the exposure time was set to 100 ms. Pixel size was 0.16 µm. Plates with cultured neurons were maintained at 37°C and 5.5% CO_2_ throughout the total imaging period. All axonal transport movies were taken in a central region of the axon, at least 150 µm away from both the soma and the axon tip.

#### Analysis of axonal transport dynamics from time-lapse movies

Axonal transport analysis was performed using the custom-made and previously validated KymoAnalyzer ImageJ package of macros.^169^ KymoAnalyzer is freely available for download from our laboratory website (www.encalada.scripps.edu/kymoanalyzer), and a detailed description of the software and the definition of all parameters calculated are available in the website and in our publication (Neumann et al., 2017). Briefly, kymographs were generated from time-lapse movies. Particle trajectories were manually assigned from the kymograph images. Track and segment-related parameters were automatically calculated by KymoAnalyzer.

### Functional rescue of tauopathy frontotemporal dementia disease model

#### Human FTD neural progenitor cell lines

Work with human induced pluripotent stem cells (iPSC) and the derived neural progenitor cell (NPC) lines was executed under the Massachusetts General Hospital/Mass General Brigham-approved IRB Protocol #2010P001611/MGH. Patient-specific iPSCs and cortical-enriched NPCs were generated as previously described.^153,188^ In this study, the NPC line used is designated MGH2046-RC1 and is derived from an individual in her 50’s with FTD and carrier of the tau-P301L autosomal dominant mutation (c.C1907T, rs63751273).^153,188^

#### NPC culture, differentiation and compound treatment

NPCs were cultured in 6-well (Fisher Scientific Corning) or black 96-well clear bottom (Fisher Scientific Corning) plates coated with poly-ornithine (20 μg/mL in water, Sigma) and laminin (5 μg/mL in PBS, Sigma), referenced to as POL-coated plates. NPC proliferation medium composition was as follows: 70% (v/v) DMEM (Gibco), 30% Ham’s-F12 (v/v) (Fisher Scientific Corning), 2% B27 (v/v) (Gibco), and 1% penicillin-streptomycin (v/v) (Gibco); freshly supplemented with 20 ng/mL EGF (Sigma), 20 ng/mL FGF (Stemgent), and 5 μg/mL heparin (Sigma) at the time of cell passaging, which was done using TrypLE reagent (Thermo Fisher). For neural differentiation over the course of six to eight weeks, cells were plated at an average density of 80,000 cells/cm^2^ in 6-well (in 2 mL medium volume) or 96-well (in 200 μL medium volume) plates POL-coated, in the same NPC media but without growth factors. Neuronal culture feeding consisted of half-medium volume changes twice a week.^153^ Compound treatment in 6-well plate format was performed by removing 1 mL of medium from the culture and adding 1 mL of new medium pre-mixed with the compound at the appropriate 2X final concentration (final volume 2 mL), followed by incubation at 37°C for the designated time. For compound treatment in 96-well plates, cells were in 100 μL media, and compound was added directly to the medium at 1X final concentration. CCT020312 synthesized in the Kelly lab was used. As a control, neurons were treated with vehicle alone (DMSO 0.1% (v/v)).

#### Neuronal cell lysis and western blot analysis

Neurons differentiated and treated in 6-well plates were washed with DPBS (Corning), detached from the well into suspension by scrapping, transferred to Eppendorf tubes, and centrifuged at 3,000 *x g* for 5 min. Cell pellets were lysed directly in 8X the cell pellet volume with 2% SDS blue loading buffer (New England Biolabs), containing DTT, incubated at RT for 15 min, and boiled for 15 min prior to SDS-PAGE. Electrophoresis was performed with 10 μL of sample per well using the Novex NuPAGE SDS-PAGE Gel System (Invitrogen). Proteins were transferred from the gel onto PVDF membranes (EMD Millipore) using standard methods. Membranes were blocked in 5% (w/v) BSA (Sigma) in Tris-buffered saline with Tween-20 (TBST, Boston Bio-Products) and incubated overnight in primary antibody prepared in 5% (w/v) BSA-TBST at 4°C. After membrane washing in TBST, these were incubated with the corresponding HRP-linked secondary antibody at 1:4000 dilution (Cell Signaling Technology). Membranes were incubated with SuperSignal West Pico Chemiluminescent Substrate (Thermo Fisher) according to manufacturer’s instructions and exposed to autoradiographic films (LabScientific by Thermo Fisher). Image capture was done with an Epson Perfection V800 Photo Scanner. Protein bands’ densitometry quantification (pixel mean intensity in arbitrary units, a.u. or OD) was done with Adobe Photoshop 2021 Histogram function and normalized to the respective internal control (β-Actin or GAPDH) band. Calculations were performed in Microsoft Excel (v.16.59), and graphs were plotted in GraphPad Prism 9.

The antibodies used were as follows: total tau TAU5 from Invitrogen (AHB0042), P-tau Ser202/Thr205 or AT8 from Thermo Scientific (MN1020), P-tau Ser396 from Invitrogen (44752G), β-III-tubulin from Sigma (T-8660), GAPDH from Abcam (ab8245), and β-ACTIN from Sigma (A1978).

#### Protein solubility analysis

Cell lysis and fractionation based on protein differential solubility to detergents Triton-X100 and SDS was performed as previously described.^151–153^ Briefly, cell pellets corresponding to two wells of a 6-well plate (double-well pellet with same treatment conditions) were lysed in 1% (v/v) TritonX-100 buffer (Fisher Scientific) in DPBS supplemented with 1% (v/v) Halt Protease/Phosphatase inhibitors (Thermo Fisher Scientific), 1:5000 Benzonase (Sigma), and 10mM DTT (New England BioLabs). Lysates were centrifuged at 14,000 *x g* for 10 min at 4°C, and the supernatants containing higher solubility proteins (S fractions) were transferred to new tubes. The pellets were resuspended in 5% (v/v) SDS (Sigma) RIPA buffer supplemented with 1% (v/v) Halt Protease/Phosphatase inhibitors (Thermo Fisher Scientific), 1:5000 Benzonase (Sigma), and 10mM DTT (New England BioLabs) and centrifuged at 20,000 *x g* for 2 min at 20°C. The new supernatants contain protein of lower solubility or insoluble protein (P fractions). SDS-PAGE western blot was performed by loading 20 μg of each S-fraction and equal volume of the P-fraction onto precast Tris-Acetate SDS-PAGE (Novex, Invitrogen). Western blots were performed as before. Densitometry quantification (pixel mean intensity in arbitrary units, a.u. or OD) was done with the Histogram function of Adobe Photoshop 2021 and normalized to the respective GAPDH intensity in the S-fraction. Calculations were done in Microsoft Excel (v.16.59) and graphs were plotted in GraphPad Prism 9.

#### Time-course analysis of tau levels

Neurons were differentiated for six weeks and on d 0, vehicle alone (DMSO) or compound CCT020312 was added to the neuronal cultures and incubated for 24 h at 37°C. On d 1, neuronal medium was replaced with new medium without compound (compound washout). At each time point after the 24-h treatment and up to 16 d post-treatment, one well of cells was sacrificed for direct lysis in 2% SDS blue loading buffer (New England Biolabs) and protein analysis of tau and β-III-tubulin (Sigma T-8660) by western blot as described above.

#### Stress vulnerability assays

Performed as previously described.^153^ Briefly, NPCs were plated at an average density of 90,000 cells/cm^2^ in 96-well plates (black, clear flat bottom, Corning) and differentiated for eight weeks. Compound CCT020312 (1µM or 10µM) or DMSO (Sigma) were added directly onto the media and incubated for 8 h at 37°C. Then, wells were treated with either 25µM Aβ(1-42) (Abcam), 400µM NMDA (Enzo LifeSciences), 5µM rotenone (Sigma), or vehicle/DMSO alone for an additional 16 h. At 24 h, viability was measured as described above, with alamarBlue HS cell viability reagent (Thermo Fisher) at 1:10 dilution with 3 h incubation at 37°C, according to manufacturer’s instructions. Calculations and statistics were done in Microsoft Excel version 16.59; graphs were plotted in GraphPad Prism 9.

#### Immunofluorescence of neuronal cells

NPCs were plated at a starting density of 90,000 cells/cm^2^ in black POL-coated 96-well clear-bottom plates (Corning), in DMEM/F12-B27 media and differentiated for six weeks, followed by compound treatment for 24 h. For LysoTracker live-cell staining, cells were incubated with LysoTracker Red DND-99 (Life Technologies L-7528) at 500nM for 45 min at 37°C, according to manufacturer’s protocols. Then, all neurons were fixed with 4% (v/v) formaldehyde-PBS (Tousimis) for 30 min, washed in DPBS (Corning), and incubated for 2 h with permeabilization/blocking buffer with the following composition: 10 mg/mL BSA (Sigma), 0.05% (v/v) Tween-20 (Bio-Rad), 2% (v/v) goat serum (Life Technologies), 0.1% Triton X-100 (Bio-Rad) in PBS. Cells were then incubated with primary antibodies overnight at 4°C: P-Tau PHF1 (kind gift from Dr. Peter Davies), MAP2 (Millipore AB5543), LAMP1 (Cell Signaling Technology 9091), and Hoechst 33342 (Thermo Fisher). Afterward, cells were washed with PBS and incubated with the corresponding AlexaFluor-conjugated secondary antibodies (Life Technologies). Image acquisition was done with the InCell Analyzer 6000 Cell Imaging System (GE Healthcare Life Sciences) and micrographs were assembled using Adobe Photoshop 2022.

#### Statistical information

Graphed data represent mean values and error bars represent SD (standard deviation) or SEM (standard error of the mean), calculated using Microsoft Excel and GraphPad Prism. *P* value < 0.05 was considered the threshold for statistical significance. *P* value significance intervals (*) are provided within each figure legend, together with the statistical test performed for each experiment. *N* values are indicated within figure legends and refer to biological replicates (NPC differentiation cultures independent setup and analysis, at different times), whereas technical replicates refer to the repeated analysis of the same samples. Derived statistics correspond to analysis of averaged values across biological replicates.

### Functional rescue of AD model

#### Human dermal fibroblast cell culture and direct neuronal differentiation

Human dermal fibroblasts were acquired from Stanford Alzheimer’s Disease Research Center (ADRC). Cells were cultured in DMEM medium with high glucose and GlutaMAX containing 10% FBS, 1% penicillin/streptomycin, MEM Non-Essential Amino Acids, 1% sodium pyruvate, and 0.1% beta-mercaptoethanol (Thermo Fisher Scientific).

To generate directly induced neurons (iNs), fibroblasts were transduced with lentiviral vectors of FUW-M2rtTA and doxycycline-inducible Ngn2-puro, Ascl1, Brn2, and Myt1l.

The transduced cells were selected by 0.5 µg/mL puromycin (Thermo Fisher Scientific) for 2 d and then received magnetic cell sorting to enrich the PSA-NCAM-positive cells. These PSA-NCAM-positive cells were replated onto the poly-L-ornithine/laminin–coated coverslips and cultured in the Reprogramming Medium: DMEM/F-12 and neurobasal medium supplemented with 0.25% GlutaMAX, 1% N-2, 2% B-27, and 1% Penicillin-Streptomycin (Thermo Fisher Scientific) for 3 weeks. The Reprogramming Medium was added with 5µM Forskolin (Sigma-Aldrich), 2µM Dorsomorphin (Tocris), 10µM SB-431542 (Tocris), 1 µg/mL doxycycline (Cayman), 10 ng/mL BDNF (Peprotech), and 10 ng/mL NT-3 (Peprotech) to promote neuronal differentiation. Then, the cells were switched to Neuronal Medium: BrainPhys neuronal medium (STEMCELL Technologies) supplemented with 0.25% GlutaMAX, 1% N-2, 2% B-27, 1% Penicillin-Streptomycin (Thermo Fisher Scientific), 5µM Forskolin (Sigma-Aldrich), 2 µM Dorsomorphin (Tocris), 10 µM SB-431542 (Tocris), 1 µg/mL doxycycline (Cayman), 10 ng/mL BDNF (Peprotech), and 10 ng/mL NT-3 (Peprotech) to promote neuronal maturation, and cultured for additional 2 weeks. Half of the culture medium was changed every 2 to 3 d.

#### iNS immunofluorescence

The iNs were plated at a starting density of 5×10^4^ cells in 24-well plates in neuronal media and treated with either 0.1% DMSO, 1µM BACE-1 inhibitor (β-Secretase Inhibitor IV, Sigma-Aldrich #565788), 250nM rapamycin, or CCT020312 for 48 h, all at a final DMSO concentration of 0.1%. Then, all neurons were fixed with 4% PFA for 15 min, washed in PBS, and incubated for 5 min with 0.3% Triton X-100 for permeabilization and for 45 min with 5% BSA for blocking. Cells were then incubated with primary antibodies overnight at 4°C: Amyloid β (D54D2) (Cell Signaling) at 1:100 dilution, Amyloid β A4 (CT, 1-42) (8G7) (Enzo Life Sciences) at 1:200 dilution, and Tubulin β-III (Tuj1) (Neuromics) at 1:1000 dilution. Afterwards, cells were washed with PBS and incubated with Alexa Fluor 488-, Alexa Fluor 555-, and Alexa Fluor 647-conjugated secondary antibodies at 1:500 dilution for 1 at RT. Image acquisition was done with the Zeiss LSM 700 confocal fluorescence microscope. Images were analyzed by calculating mean fluorescence intensity of total Aβ and Aβ(1-42) in the cell body of individual neurons. The total cell numbers used for data analysis in each condition range from 88 to 142 for three independent experiments.

## References

1. Tsukada, M., and Ohsumi, Y. (1993). ISOLATION AND CHARACTERIZATION OF AUTOPHAGY-DEFECTIVE MUTANTS OF SACCHAROMYCES-CEREVISIAE. Febs Letters 333, 169–174. 10.1016/0014-5793(93)80398-e.

2. Ichimura, Y., Kirisako, T., Takao, T., Satomi, Y., Shimonishi, Y., Ishihara, N., Mizushima, N., Tanida, I., Kominami, E., Ohsumi, M., et al. (2000). A ubiquitin-like system mediates protein lipidation. Nature 408, 488–492. 10.1038/35044114.

3. Kabeya, Y., Mizushima, N., Uero, T., Yamamoto, A., Kirisako, T., Noda, T., Kominami, E., Ohsumi, Y., and Yoshimori, T. (2000). LC3, a mammalian homologue of yeast Apg8p, is localized in autophagosome membranes after processing. Embo Journal 19, 5720–5728. 10.1093/emboj/19.21.5720.

4. Lazarou, M., Sliter, D.A., Kane, L.A., Sarraf, S.A., Wang, C.X., Burman, J.L., Sideris, D.P., Fogel, A.I., and Youle, R.J. (2015). The ubiquitin kinase PINK1 recruits autophagy receptors to induce mitophagy. Nature 524, 309-+. 10.1038/nature14893.

5. Goodall, E.A., Kraus, F., and Harper, J.W. (2022). Mechanisms underlying ubiquitin-driven selective mitochondrial and bacterial autophagy. Molecular Cell 82, 1501–1513. 10.1016/j.molcel.2022.03.012.

6. Novak, I., Kirkin, V., McEwan, D.G., Zhang, J., Wild, P., Rozenknop, A., Rogov, V., Lohr, F., Popovic, D., Occhipinti, A., et al. (2010). Nix is a selective autophagy receptor for mitochondrial clearance. Embo Reports 11, 45–51. 10.1038/embor.2009.256.

7. An, H., Ordureau, A., Paulo, J.A., Shoemaker, C.J., Denic, V., and Harper, J.W. (2019). TEX264 Is an Endoplasmic Reticulum-Resident ATG8-Interacting Protein Critical for ER Remodeling during Nutrient Stress. Molecular Cell 74, 891-+. 10.1016/j.molcel.2019.03.034.

8. Khaminets, A., Heinrich, T., Mari, M., Grumati, P., Huebner, A.K., Akutsu, M., Liebmann, L., Stolz, A., Nietzsche, S., Koch, N., et al. (2015). Regulation of endoplasmic reticulum turnover by selective autophagy. Nature 522, 354-+. 10.1038/nature14498.

9. Eapen, V.V., Swarup, S., Hoyer, M.J., Paulo, J.A., and Harper, J.W. (2021). Quantitative proteomics reveals the selectivity of ubiquitin-binding autophagy receptors in the turnover of damaged lysosomes by lysophagy. eLife 10, 36, e72328. 10.7554/eLife.72328; 10.7554/eLife.72328.sa1; 10.7554/eLife.72328.sa2.

10. Jin, S.M., and Youle, R.J. (2012). PINK1- and Parkin-mediated mitophagy at a glance. J Cell Sci 125, 795–799. 10.1242/jcs.093849.

11. Lapierre, L.R., Silvestrini, M.J., Nunez, L., Ames, K., Wong, S., Le, T.T., Hansen, M., and Melendez, A. (2013). Autophagy genes are required for normal lipid levels in C. elegans. Autophagy 9, 278–286. 10.4161/auto.22930.

12. Singh, R., Kaushik, S., Wang, Y.J., Xiang, Y.Q., Novak, I., Komatsu, M., Tanaka, K., Cuervo, A.M., and Czaja, M.J. (2009). Autophagy regulates lipid metabolism. Nature 458, 1131–U1164. 10.1038/nature07976.

13. Sarraf, S.A., Shah, H.V., Kanfer, G., Pickrell, A.M., Holtzclaw, L.A., Ward, M.E., and Youle, R.J. (2020). Loss of TAX1BP1-Directed Autophagy Results in Protein Aggregate Accumulation in the Brain. Molecular Cell 80, 779–795.e710. 10.1016/j.molcel.2020.10.041.

14. Kirkin, V., McEwan, D.G., Novak, I., and Dikic, I. (2009). A Role for Ubiquitin in Selective Autophagy. Molecular Cell 34, 259–269. 10.1016/j.molcel.2009.04.026.

15. Ravikumar, B., Duden, R., and Rubinsztein, D.C. (2002). Aggregate-prone proteins with polyglutamine and polyalanine expansions are degraded by autophagy. Hum Mol Genet 11, 1107–1117.

16. An, H., and Harper, J.W. (2018). Systematic analysis of ribophagy in human cells reveals bystander flux during selective autophagy. Nature Cell Biology 20, 135-+. 10.1038/s41556-017-0007-x.

17. Kroemer, G., Mariño, G., and Levine, B. (2010). Autophagy and the Integrated Stress Response. Molecular Cell 40, 280–293. 10.1016/j.molcel.2010.09.023.

18. Shang, L., Chen, S., Du, F., Li, S., Zhao, L., and Wang, X. (2011). Nutrient starvation elicits an acute autophagic response mediated by Ulk1 dephosphorylation and its subsequent dissociation from AMPK. Proc Natl Acad Sci U S A 108, 4788–4793. 10.1073/pnas.1100844108.

19. Rubinsztein, D.C., Gestwicki, J.E., Murphy, L.O., and Klionsky, D.J. (2007). Potential therapeutic applications of autophagy. Nature Reviews Drug Discovery 6, 304–312. 10.1038/nrd2272.

20. Kaizuka, T., Morishita, H., Hama, Y., Tsukamoto, S., Matsui, T., Toyota, Y., Kodama, A., Ishihara, T., Mizushima, T., and Mizushima, N. (2016). An Autophagic Flux Probe that Releases an Internal Control. Molecular Cell 64, 835–849. 10.1016/j.molcel.2016.09.037.

21. Lipinski, M.M., Hoffman, G., Ng, A., Zhou, W., Py, B.F., Hsu, E., Liu, X.X., Eisenberg, J., Liu, J., Blenis, J., et al. (2010). A Genome-Wide siRNA Screen Reveals Multiple mTORC1 Independent Signaling Pathways Regulating Autophagy under Normal Nutritional Conditions. Developmental Cell 18, 1041–1052. 10.1016/j.devcel.2010.05.005.

22. Zhang, L.H., Yu, J., Pan, H.L., Hu, P., Hao, Y., Cai, W.Q., Zhu, H., Yu, A.D., Xie, X., Ma, D.W., and Yuan, J.Y. (2007). Small molecule regulators of autophagy identified by an image-based high-throughput screen. Proceedings of the National Academy of Sciences of the United States of America 104, 19023–19028. 10.1073/pnas.0709695104.

23. Panda, P.K., Fahmer, A., Vats, S., Seranova, E., Sharma, V., Chipara, M., Desai, P., Torresi, J., Rosenstock, T., Kumar, D., and Sarkar, S. (2019). Chemical Screening Approaches Enabling Drug Discovery of Autophagy Modulators for Biomedical Applications in Human Diseases. Front. Cell. Dev. Biol. 7, 19, 38. 10.3389/fcell.2019.00038.

24. Stolz, A., Ernst, A., and Dikic, I. (2014). Cargo recognition and trafficking in selective autophagy. Nature Cell Biology 16, 495–501. 10.1038/ncb2979.

25. Klionsky, D.J., Abdelmohsen, K., Abe, A., Abedin, M.J., Abeliovich, H., Acevedo Arozena, A., Adachi, H., Adams, C.M., Adams, P.D., Adeli, K., et al. (2016). Guidelines for the use and interpretation of assays for monitoring autophagy (3rd edition). Autophagy 12, 1–222. 10.1080/15548627.2015.1100356.

26. Lee, J.H., Yang, D.S., Goulbourne, C.N., Im, E., Stavrides, P., Pensalfini, A., Chan, H., Bouchet-Marquis, C., Bleiwas, C., Berg, M.J., et al. (2022). Faulty autolysosome acidification in Alzheimer’s disease mouse models induces autophagic build-up of A beta in neurons, yielding senile plaques. Nature Neuroscience 25, 688-+. 10.1038/s41593-022-01084-8.

27. Rubinsztein, D.C., Marino, G., and Kroemer, G. (2011). Autophagy and Aging. Cell 146, 682–695. 10.1016/j.cell.2011.07.030.

28. Yang, D.S., Stavrides, P., Saito, M., Kumar, A., Rodriguez-Navarro, J.A., Pawlik, M., Huo, C.F., Walkley, S.U., Saito, M., Cuervo, A.M., and Nixon, R.A. (2014). Defective macroautophagic turnover of brain lipids in the TgCRND8 Alzheimer mouse model: prevention by correcting lysosomal proteolytic deficits. Brain 137, 3300–3318. 10.1093/brain/awu278.

29. Kao, A.W., McKay, A., Singh, P.P., Brunet, A., and Huang, E.J. (2017). Progranulin, lysosomal regulation and neurodegenerative disease. Nature Reviews Neuroscience 18, 325–333. 10.1038/nrn.2017.36.

30. Butlers, V.J., Cortopassi, W.A., Argouarch, A.R., Ivry, S.L., Craik, C.S., Jacobson, M.P., and Kao, A.W. (2019). Progranulin Stimulates the In Vitro Maturation of Pro-Cathepsin D at Acidic pH. Journal of Molecular Biology 431, 1038–1047. 10.1016/j.jmb.2019.01.027.

31. Harold, D., Abraham, R., Hollingworth, P., Sims, R., Gerrish, A., Hamshere, M.L., Pahwa, J.S., Moskvina, V., Dowzell, K., Williams, A., et al. (2009). Genome-wide association study identifies variants at CLU and PICALM associated with Alzheimer’s disease. Nature Genetics 41, 1088–U1061. 10.1038/ng.440.

32. Jiang, Y., Sato, Y., Im, E., Berg, M., Bordi, M., Darji, S., Kumar, A., Mohan, P.S., Bandyopadhyay, U., Diaz, A., et al. (2019). Lysosomal Dysfunction in Down Syndrome Is APP-Dependent and Mediated by APP-beta CTF (C99). Journal of Neuroscience 39, 5255–5268. 10.1523/jneurosci.0578-19.2019.

33. Lee, J.H., Yu, W.H., Kumar, A., Lee, S., Mohan, P.S., Peterhoff, C.M., Wolfe, D.M., Martinez-Vicente, M., Massey, A.C., Sovak, G., et al. (2010). Lysosomal Proteolysis and Autophagy Require Presenilin 1 and Are Disrupted by Alzheimer-Related PS1 Mutations. Cell 141, 1146–U1191. 10.1016/j.cell.2010.05.008.

34. Bourdenx, M., Martín-Segura, A., Scrivo, A., Rodriguez-Navarro, J.A., Kaushik, S., Tasset, I., Diaz, A., Storm, N.J., Xin, Q., Juste, Y.R., et al. (2021). Chaperone-mediated autophagy prevents collapse of the neuronal metastable proteome. Cell 184, 2696–2714.e2625. 10.1016/j.cell.2021.03.048.

35. Cotman, S.L., Karaa, A., Staropoli, J.F., and Sims, K.B. (2013). Neuronal Ceroid Lipofuscinosis: Impact of Recent Genetic Advances and Expansion of the Clinicopathologic Spectrum. Curr. Neurol. Neurosci. Rep. 13, 11, 366. 10.1007/s11910-013-0366-z.

36. von Kleist, L., Ariunbat, K., Braren, I., Stauber, T., Storch, S., and Danyukova, T. (2019). A newly generated neuronal cell model of CLN7 disease reveals aberrant lysosome motility and impaired cell survival. Mol. Genet. Metab. 126, 196–205. 10.1016/j.ymgme.2018.09.009.

37. Brady, O.A., Zheng, Y.Q., Murphy, K., Huang, M., and Hu, F.H. (2013). The frontotemporal lobar degeneration risk factor, TMEM106B, regulates lysosomal morphology and function. Human Molecular Genetics 22, 685–695. 10.1093/hmg/dds475.

38. Alquezar, C., Schoch, K.M., Geier, E.G., Ramos, E.M., Scrivo, A., Li, K.H., Argouarch, A.R., Mlynarski, E.E., Dombroski, B., DeTure, M., et al. (2021). TSC1 loss increases risk for tauopathy by inducing tau acetylation and preventing tau clearance via chaperone-mediated autophagy. Sci. Adv. 7, 15, eabg3897. 10.1126/sciadv.abg3897.

39. Ramirez, A., Heimbach, A., Gruendemann, J., Stiller, B., Hampshire, D., Cid, L.P., Goebel, I., Mubaidin, A.F., Wriekat, A.L., Roeper, J., et al. (2006). Hereditary parkinsonism with dementia is caused by mutations in ATP13A2, encoding a lysosomal type 5 P-type ATPase. Nature Genetics 38, 1184–1191. 10.1038/ng1884.

40. Papassotiropoulos, A., Bagli, M., Kurz, A., Kornhuber, J., Forstl, H., Maier, W., Pauls, J., Lautenschlager, N., and Heun, R. (2000). A genetic variation of cathepsin D is a major risk factor for Alzheimer’s disease. Annals of Neurology 47, 399–403. 10.1002/1531-8249(200003)47:3<399::Aid-ana22>3.0.Co;2-5.

41. Sidransky, E., Nalls, M.A., Aasly, J.O., Aharon-Peretz, J., Annesi, G., Barbosa, E.R., Bar-Shira, A., Berg, D., Bras, J., Brice, A., et al. (2009). Multicenter analysis of glucocerebrosidase mutations in Parkinson’s disease. N Engl J Med 361, 1651–1661. 10.1056/NEJMoa0901281.

42. Zavodszky, E., Seaman, M.N.J., Moreau, K., Jimenez-Sanchez, M., Breusegem, S.Y., Harbour, M.E., and Rubinsztein, D.C. (2014). Mutation in VPS35 associated with Parkinson’s disease impairs WASH complex association and inhibits autophagy. Nature Communications 5, 3828. 10.1038/ncomms4828.

43. Clayton, E.L., Mizielinska, S., Edgar, J.R., Nielsen, T.T., Marshall, S., Norona, F.E., Robbins, M., Damirji, H., Holm, I.E., Johannsen, P., et al. (2015). Frontotemporal dementia caused by CHMP2B mutation is characterised by neuronal lysosomal storage pathology. Acta Neuropathologica 130, 511–523. 10.1007/s00401-015-1475-3.

44. Bucci, C., Thomsen, P., Nicoziani, P., McCarthy, J., and van Deurs, B. (2000). Rab7: a key to lysosome biogenesis. Mol Biol Cell 11, 467–480. 10.1091/mbc.11.2.467.

45. Cherry, S., Jin, E.J., Ozel, M.N., Lu, Z., Agi, E., Wang, D., Jung, W.H., Epstein, D., Meinertzhagen, I.A., Chan, C.C., and Hiesinger, P.R. (2013). Charcot-Marie-Tooth 2B mutations in rab7 cause dosage-dependent neurodegeneration due to partial loss of function. eLife 2, e01064. 10.7554/eLife.01064.

46. Wan, H., Wang, Q., Chen, X., Zeng, Q., Shao, Y., Fang, H., Liao, X., Li, H.S., Liu, M.G., Xu, T.L., et al. (2020). WDR45 contributes to neurodegeneration through regulation of ER homeostasis and neuronal death. Autophagy 16, 531–547. 10.1080/15548627.2019.1630224.

47. Yang, Y., Hentati, A., Deng, H.-X., Dabbagh, O., Sasaki, T., Hirano, M., Hung, W.-Y., Ouahchi, K., Yan, J., Azim, A.C., et al. (2001). The gene encoding alsin, a protein with three guanine-nucleotide exchange factor domains, is mutated in a form of recessive amyotrophic lateral sclerosis. Nature Genetics 29, 160–165. 10.1038/ng1001-160.

48. Lai, C., Xie, C., Shim, H., Chandran, J., Howell, B.W., and Cai, H. (2009). Regulation of endosomal motility and degradation by amyotrophic lateral sclerosis 2/alsin. Mol Brain 2, 23. 10.1186/1756-6606-2-23.

49. Metzger, S., Saukko, M., Van Che, H., Tong, L., Puder, Y., Riess, O., and Nguyen, H.P. (2010). Age at onset in Huntington’s disease is modified by the autophagy pathway: implication of the V471A polymorphism in Atg7. Hum Genet 128, 453–459. 10.1007/s00439-010-0873-9.

50. Chang, J., Lee, S., and Blackstone, C. (2014). Spastic paraplegia proteins spastizin and spatacsin mediate autophagic lysosome reformation. Journal of Clinical Investigation 124, 5249–5262. 10.1172/jci77598.

51. Tian, Y., Chang, J.C., Fan, E.Y., Flajolet, M., and Greengard, P. (2013). Adaptor complex AP2/PICALM, through interaction with LC3, targets Alzheimer’s APP-CTF for terminal degradation via autophagy. Proceedings of the National Academy of Sciences of the United States of America 110, 17071–17076. 10.1073/pnas.1315110110.

52. Freischmidt, A., Wieland, T., Richter, B., Ruf, W., Schaeffer, V., Müller, K., Marroquin, N., Nordin, F., Hübers, A., Weydt, P., et al. (2015). Haploinsufficiency of TBK1 causes familial ALS and fronto-temporal dementia. Nature Neuroscience 18, 631–636. 10.1038/nn.4000.

53. Oakes, J.A., Davies, M.C., and Collins, M.O. (2017). TBK1: a new player in ALS linking autophagy and neuroinflammation. Molecular Brain 10, 5. 10.1186/s13041-017-0287-x.

54. Ochaba, J., Lukacsovich, T., Csikos, G., Zheng, S., Margulis, J., Salazar, L., Mao, K., Lau, A.L., Yeung, S.Y., Humbert, S., et al. (2014). Potential function for the Huntingtin protein as a scaffold for selective autophagy. Proceedings of the National Academy of Sciences 111, 16889–16894. doi:10.1073/pnas.1420103111.

55. Watanabe, S., Ilieva, H., Tamada, H., Nomura, H., Komine, O., Endo, F., Jin, S., Mancias, P., Kiyama, H., and Yamanaka, K. (2016). Mitochondria-associated membrane collapse is a common pathomechanism in SIGMAR1- and SOD1-linked ALS. EMBO Molecular Medicine 8, 1421–1437. 10.15252/emmm.201606403.

56. Hara, T., Nakamura, K., Matsui, M., Yamamoto, A., Nakahara, Y., Suzuki-Migishima, R., Yokoyama, M., Mishima, K., Saito, I., Okano, H., and Mizushima, N. (2006). Suppression of basal autophagy in neural cells causes neurodegenerative disease in mice. Nature 441, 885–889. 10.1038/nature04724.

57. Komatsu, M., Waguri, S., Chiba, T., Murata, S., Iwata, J., Tanida, I., Ueno, T., Koike, M., Uchiyama, Y., Kominami, E., and Tanaka, K. (2006). Loss of autophagy in the central nervous system causes neurodegeneration in mice. Nature 441, 880–884. 10.1038/nature04723.

58. Metzger, S., Saukko, M., Che, H.V., Tong, L.A., Puder, Y., Riess, O., and Nguyen, H.P. (2010). Age at onset in Huntington’s disease is modified by the autophagy pathway: implication of the V471A polymorphism in Atg7. Human Genetics 128, 453–459. 10.1007/s00439-010-0873-9.

59. Hou, Y., Dan, X., Babbar, M., Wei, Y., Hasselbalch, S.G., Croteau, D.L., and Bohr, V.A. (2019). Ageing as a risk factor for neurodegenerative disease. Nat Rev Neurol 15, 565–581. 10.1038/s41582-019-0244-7.

60. Cherra, S.J., 3rd, and Chu, C.T. (2008). Autophagy in neuroprotection and neurodegeneration: A question of balance. Future Neurol 3, 309–323. 10.2217/14796708.3.3.309.

61. Rubinsztein, D.C., Codogno, P., and Levine, B. (2012). Autophagy modulation as a potential therapeutic target for diverse diseases. Nature Reviews Drug Discovery 11, 709–U784.

62. Nixon, R.A. (2013). The role of autophagy in neurodegenerative disease. Nature Medicine 19, 983–997. 10.1038/nm.3232.

63. Duan, L.S., Hu, M.F., Tamm, J.A., Grinberg, Y.Y., Shen, F., Chai, Y.T., Xi, H.L., Gibilisco, L., Ravikumar, B., Gautam, V., et al. (2021). Arrayed CRISPR reveals genetic regulators of tau aggregation, autophagy and mitochondria in Alzheimer’s disease model. Scientific Reports 11, 17, 2879. 10.1038/s41598-021-82658-7.

64. Silva, M.C., Nandi, G.A., Tentarelli, S., Gurrell, I.K., Jamier, T., Lucente, D., Dickerson, B.C., Brown, D.G., Brandon, N.J., and Haggarty, S.J. (2020). Prolonged tau clearance and stress vulnerability rescue by pharmacological activation of autophagy in tauopathy neurons. Nature Communications 11. 10.1038/s41467-020-16984-1.

65. Thellung, S., Corsaro, A., Nizzari, M., Barbieri, F., and Florio, T. (2019). Autophagy Activator Drugs: A New Opportunity in Neuroprotection from Misfolded Protein Toxicity. Int J Mol Sci 20. 10.3390/ijms20040901.

66. Wang, C., Xiong, M.N.C., Gratuze, M., Bao, X., Shi, Y., Andhey, P.S., Manis, M., Schroeder, C., Yin, Z.R., Madore, C., et al. (2021). Selective removal of astrocytic APOE4 strongly protects against tau-mediated neurodegeneration and decreases synaptic phagocytosis by microglia. Neuron 109, 1657-+. 10.1016/j.neuron.2021.03.024.

67. Crother, T.R., Porritt, R.A., Dagvadorj, J., Tumurkhuu, G., Slepenkin, A.V., Peterson, E.M., Chen, S., Shimada, K., and Arditi, M. (2019). Autophagy Limits Inflammasome During Chlamydia pneumoniae Infection. Frontiers in Immunology 10, 11, 754. 10.3389/fimmu.2019.00754.

68. Zhong, Z.Y., Umemura, A., Sanchez-Lopez, E., Liang, S., Shalapour, S., Wong, J., He, F., Boassa, D., Perkins, G., Ali, S.R., et al. (2016). NF-kappa B Restricts Inflammasome Activation via Elimination of Damaged Mitochondria. Cell 164, 896–910. 10.1016/j.cell.2015.12.057.

69. Yamanaka, K., Chun, S.J., Boillee, S., Fujimori-Tonou, N., Yamashita, H., Gutmann, D.H., Takahashi, R., Misawa, H., and Cleveland, D.W. (2008). Astrocytes as determinants of disease progression in inherited amyotrophic lateral sclerosis. Nat Neurosci 11, 251–253. 10.1038/nn2047.

70. Thoreen, C.C., Kang, S.A., Chang, J.W., Liu, Q., Zhang, J., Gao, Y., Reichling, L.J., Sim, T., Sabatini, D.M., and Gray, N.S. (2009). An ATP-competitive mammalian target of rapamycin inhibitor reveals rapamycin-resistant functions of mTORC1. J Biol Chem 284, 8023–8032. 10.1074/jbc.M900301200.

71. Chin, M.Y., Ang, K.H., Davies, J., Alquezar, C., Garda, V.G., Rooney, B., Leng, K., Kampmann, M., Arkin, M.R., and Kao, A.W. (2022). Phenotypic Screening Using High-Content Imaging to Identify Lysosomal pH Modulators in a Neuronal Cell Model. ACS Chem. Neurosci. 13, 1505–1516. 10.1021/acschemneuro.1c00804.

72. Safren, N., Tank, E.M., Malik, A.M., Chua, J.P., Santoro, N., and Barmada, S.J. (2021). Development of a specific live-cell assay for native autophagic flux. Journal of Biological Chemistry 297. 10.1016/j.jbc.2021.101003.

73. Balgi, A.D., Fonseca, B.D., Donohue, E., Tsang, T.C.F., Lajoie, P., Proud, C.G., Nabi, I.R., and Roberge, M. (2009). Screen for Chemical Modulators of Autophagy Reveals Novel Therapeutic Inhibitors of mTORC1 Signaling. Plos One 4, e7124. 10.1371/journal.pone.0007124.

74. Chauhan, S., Ahmed, Z., Bradfute, S.B., Arko-Mensah, J., Mandell, M.A., Choi, S.W., Kimura, T., Blanchet, F., Waller, A., Mudd, M.H., et al. (2015). Pharmaceutical screen identifies novel target processes for activation of autophagy with a broad translational potential. Nature Communications 6, 8620. 10.1038/ncomms9620.

75. Sarkar, S., Ravikumar, B., Floto, R.A., and Rubinsztein, D.C. (2009). Rapamycin and mTOR-independent autophagy inducers ameliorate toxicity of polyglutamine-expanded huntingtin and related proteinopathies. Cell Death Differ 16, 46–56. 10.1038/cdd.2008.110.

76. Laplante, M., and Sabatini, David M. (2012). mTOR Signaling in Growth Control and Disease. Cell 149, 274–293. 10.1016/j.cell.2012.03.017.

77. Saxton, R.A., and Sabatini, D.M. (2017). mTOR Signaling in Growth, Metabolism, and Disease. Cell 168, 960–976. 10.1016/j.cell.2017.02.004.

78. Thomson, A.W., Turnquist, H.R., and Raimondi, G. (2009). Immunoregulatory functions of mTOR inhibition. Nat Rev Immunol 9, 324–337.

79. Hua, H., Kong, Q., Zhang, H., Wang, J., Luo, T., and Jiang, Y. (2019). Targeting mTOR for cancer therapy. Journal of Hematology & Oncology 12, 71. 10.1186/s13045-019-0754-1.

80. Sarkar, S., Davies, J.E., Huang, Z.B., Tunnacliffe, A., and Rubinsztein, D.C. (2007). Trehalose, a novel mTOR-independent autophagy enhancer, accelerates the clearance of mutant huntingtin and alpha-synuclein. Journal of Biological Chemistry 282, 5641–5652. 10.1074/jbc.M609532200.

81. Sarkar, S., Floto, R.A., Berger, Z., Imarisio, S., Cordenier, A., Pasco, M., Cook, L.J., and Rubinsztein, D.C. (2005). Lithium induces autophagy by inhibiting inositol monophosphatase. Journal of Cell Biology 170, 1101–1111. 10.1083/jcb.200504035.

82. Lee, S., Kim, E., and Park, S.B. (2013). Discovery of autophagy modulators through the construction of a high-content screening platform via monitoring of lipid droplets. Chemical Science 4, 3282–3287. 10.1039/c3sc51344k.

83. Hofer, S.J., Simon, A.K., Bergmann, M., Eisenberg, T., Kroemer, G., and Madeo, F. (2022). Mechanisms of spermidine-induced autophagy and geroprotection. Nature Aging 2, 1112–1129. 10.1038/s43587-022-00322-9.

84. Shoji-Kawata, S., Sumpter, R., Leveno, M., Campbell, G.R., Zou, Z., Kinch, L., Wilkins, A.D., Sun, Q., Pallauf, K., MacDuff, D., et al. (2013). Identification of a candidate therapeutic autophagy-inducing peptide. Nature 494, 201–206. 10.1038/nature11866.

85. Rusmini, P., Cortese, K., Crippa, V., Cristofani, R., Cicardi, M.E., Ferrari, V., Vezzoli, G., Tedesco, B., Meroni, M., Messi, E., et al. (2019). Trehalose induces autophagy via lysosomal-mediated TFEB activation in models of motoneuron degeneration. Autophagy 15, 631–651. 10.1080/15548627.2018.1535292.

86. Pupyshev, A.B., Belichenko, V.M., Tenditnik, M.V., Bashirzade, A.A., Dubrovina, N.I., Ovsyukova, M.V., Akopyan, A.A., Fedoseeva, L.A., Korolenko, T.A., Amstislavskaya, T.G., and Tikhonova, M.A. (2022). Combined induction of mTOR-dependent and mTOR-independent pathways of autophagy activation as an experimental therapy for Alzheimer’s disease-like pathology in a mouse model. Pharmacol Biochem Behav 217, 173406. 10.1016/j.pbb.2022.173406.

87. Hamilton, L.K., Dufresne, M., Joppe, S.E., Petryszyn, S., Aumont, A., Calon, F., Barnabe-Heider, F., Furtos, A., Parent, M., Chaurand, P., and Fernandes, K.J. (2015). Aberrant Lipid Metabolism in the Forebrain Niche Suppresses Adult Neural Stem Cell Proliferation in an Animal Model of Alzheimer’s Disease. Cell Stem Cell 17, 397–411. 10.1016/j.stem.2015.08.001.

88. Martin, S., and Parton, R.G. (2006). Lipid droplets: a unified view of a dynamic organelle. Nature Reviews Molecular Cell Biology 7, 373–378. 10.1038/nrm1912.

89. Murphy, D.J. (2001). The biogenesis and functions of lipid bodies in animals, plants and microorganisms. Progress in Lipid Research 40, 325–438. 10.1016/s0163-7827(01)00013-3.

90. Olzmann, J.A., and Carvalho, P. (2019). Dynamics and functions of lipid droplets. Nat Rev Mol Cell Biol 20, 137–155. 10.1038/s41580-018-0085-z.

91. Renne, M.F., and Hariri, H. (2021). Lipid Droplet-Organelle Contact Sites as Hubs for Fatty Acid Metabolism, Trafficking, and Metabolic Channeling. Front Cell Dev Biol 9, 726261. 10.3389/fcell.2021.726261.

92. Walther, T.C., Chung, J., and Farese, R.V., Jr. (2017). Lipid Droplet Biogenesis. Annu Rev Cell Dev Biol 33, 491–510. 10.1146/annurev-cellbio-100616-060608.

93. Spangenburg, E.E., Pratt, S.J.P., Wohlers, L.M., and Lovering, R.M. (2011). Use of BODIPY (493/503) to visualize intramuscular lipid droplets in skeletal muscle. J Biomed Biotechnol 2011, 598358. 10.1155/2011/598358.

94. Ranall, M.V., Gabrielli, B.G., and Gonda, T.J. (2011). High-content imaging of neutral lipid droplets with 1,6-diphenylhexatriene. Biotechniques 51, 35–42. 10.2144/000113702.

95. Qiu, B., and Simon, M.C. (2016). BODIPY 493/503 Staining of Neutral Lipid Droplets for Microscopy and Quantification by Flow Cytometry. Bio Protoc 6. 10.21769/BioProtoc.1912.

96. Corrêa, R., Silva, L.F.F., Ribeiro, D.J.S., Almeida, R.d.N., Santos, I.d.O., Corrêa, L.H., de Sant’Ana, L.P., Assunção, L.S., Bozza, P.T., and Magalhães, K.G. (2020). Lysophosphatidylcholine Induces NLRP3 Inflammasome-Mediated Foam Cell Formation and Pyroptosis in Human Monocytes and Endothelial Cells. Frontiers in Immunology 10. 10.3389/fimmu.2019.02927.

97. Lass, A., Zimmermann, R., Haemmerle, G., Riederer, M., Schoiswohl, G., Schweiger, M., Kienesberger, P., Strauss, J.G., Gorkiewicz, G., and Zechner, R. (2006). Adipose triglyceride lipase-mediated lipolysis of cellular fat stores is activated by CGI-58 and defective in Chanarin-Dorfman Syndrome. Cell Metab 3, 309–319. 10.1016/j.cmet.2006.03.005.

98. Cingolani, F., and Czaja, M.J. (2016). Regulation and Functions of Autophagic Lipolysis. Trends Endocrinol Metab 27, 696–705. 10.1016/j.tem.2016.06.003.

99. Roberts, M.A., and Olzmann, J.A. (2020). Protein Quality Control and Lipid Droplet Metabolism. Annu Rev Cell Dev Biol 36, 115–139. 10.1146/annurev-cellbio-031320-101827.

100. Jiang, L., Wang, W., He, Q., Wu, Y., Lu, Z., Sun, J., Liu, Z., Shao, Y., and Wang, A. (2017). Oleic acid induces apoptosis and autophagy in the treatment of Tongue Squamous cell carcinomas. Sci Rep 7, 11277. 10.1038/s41598-017-11842-5.

101. Klionsky, D.J., Abdel-Aziz, A.K., Abdelfatah, S., Abdellatif, M., Abdoli, A., Abel, S., Abeliovich, H., Abildgaard, M.H., Abudu, Y.P., Acevedo-Arozena, A., et al. (2021). Guidelines for the use and interpretation of assays for monitoring autophagy (4th edition)(1). Autophagy 17, 1–382. 10.1080/15548627.2020.1797280.

102. Towers, C.G. (2018). Measuring Autophagic Flux with LC3 protein levels: The do’s and don’ts. https://www.novusbio.com/antibody-news/measuring-autophagic-flux-with-lc3-protein-levels-the-dos-and-donts.

103. Kim, E., Lee, S., and Park, S.B. (2012). A Seoul-Fluor-based bioprobe for lipid droplets and its application in image-based high throughput screening. Chemical Communications 48, 2331–2333. 10.1039/c2cc17496k.

104. Grandl, M., and Schmitz, G. (2010). Fluorescent High-Content Imaging Allows the Discrimination and Quantitation of E-LDL-Induced Lipid Droplets and Ox-LDL-Generated Phospholipidosis in Human Macrophages. Cytometry Part A 77A, 231–242. 10.1002/cyto.a.20828.

105. Rodrigues, A.C., Prímola-Gomes, T.N., Peluzio, M.C.G., Hermsdorff, H.H.M., and Natali, A.J. (2021). Aerobic exercise and lipolysis: A review of the β-adrenergic signaling pathways in adipose tissue. Science & Sports 36, 16–26. 10.1016/j.scispo.2020.04.006.

106. Baell, J.B., and Nissink, J.W.M. (2018). Seven Year Itch: Pan-Assay Interference Compounds (PAINS) in 2017—Utility and Limitations. ACS Chemical Biology 13, 36–44. 10.1021/acschembio.7b00903.

107. Stockwell, S.R., Platt, G., Barrie, S.E., Zoumpoulidou, G., Poele, R.H.T., Aherne, G.W., Wilson, S.C., Sheldrake, P., McDonald, E., Venet, M., et al. (2012). Mechanism-Based Screen for G1/S Checkpoint Activators Identifies a Selective Activator of EIF2AK3/PERK Signalling. Plos One 7, 16, e28568. 10.1371/journal.pone.0028568.

108. Luhr, M., Torgersen, M.L., Szalai, P., Hashim, A., Brech, A., Staerk, J., and Engedal, N. (2019). The kinase PERK and the transcription factor ATF4 play distinct and essential roles in autophagy resulting from tunicamycin-induced ER stress. J Biol Chem 294, 8197–8217. 10.1074/jbc.RA118.002829.

109. Owen, S.C., Doak, A.K., Wassam, P., Shoichet, M.S., and Shoichet, B.K. (2012). Colloidal aggregation affects the efficacy of anticancer drugs in cell culture. ACS Chem Biol 7, 1429–1435. 10.1021/cb300189b.

110. Jung, M., Choi, H., and Mun, J.Y. (2019). The autophagy research in electron microscopy. Appl Microsc 49, 11. 10.1186/s42649-019-0012-6.

111. Neikirk, K., Vue, Z., Katti, P., Rodriguez, B.I., Omer, S., Shao, J., Christensen, T., Garza Lopez, E., Marshall, A., Palavicino-Maggio, C.B., et al. (2023). Systematic Transmission Electron Microscopy-Based Identification and 3D Reconstruction of Cellular Degradation Machinery. Adv Biol (Weinh) 7, e2200221. 10.1002/adbi.202200221.

112. Mauthe, M., Orhon, I., Rocchi, C., Zhou, X., Luhr, M., Hijlkema, K.-J., Coppes, R.P., Engedal, N., Mari, M., and Reggiori, F. (2018). Chloroquine inhibits autophagic flux by decreasing autophagosome-lysosome fusion. Autophagy 14, 1435–1455. 10.1080/15548627.2018.1474314.

113. Eskelinen, E.-L., and Kovács, A.L. (2011). Double membranes vs. lipid bilayers, and their significance for correct identification of macroautophagic structures. Autophagy 7, 931–932. 10.4161/auto.7.9.16679.

114. Eskelinen, E.-L. (2008). To be or not to be? Examples of incorrect identification of autophagic compartments in conventional transmission electron microscopy of mammalian cells. Autophagy 4, 257–260. 10.4161/auto.5179.

115. Stockwell, S.R., and Mittnacht, S. (2014). Workflow for High-content, Individual Cell Quantification of Fluorescent Markers from Universal Microscope Data, Supported by Open Source Software. Jove-Journal of Visualized Experiments, e51882. 10.3791/51882.

116. Chen, T., Ozel, D., Qiao, Y., Harbinski, F., Chen, L., Denoyelle, S., He, X., Zvereva, N., Supko, J.G., Chorev, M., et al. (2011). Chemical genetics identify eIF2α kinase heme-regulated inhibitor as an anticancer target. Nature Chemical Biology 7, 610–616. 10.1038/nchembio.613.

117. Axten, J.M., Romeril, S.P., Shu, A., Ralph, J., Medina, J.R., Feng, Y., Li, W.H., Grant, S.W., Heerding, D.A., Minthorn, E., et al. (2013). Discovery of GSK2656157: An Optimized PERK Inhibitor Selected for Preclinical Development. ACS Med Chem Lett 4, 964–968. 10.1021/ml400228e.

118. Sidrauski, C., McGeachy, A.M., Ingolia, N.T., and Walter, P. (2015). The small molecule ISRIB reverses the effects of eIF2α phosphorylation on translation and stress granule assembly. eLife 4, e05033. 10.7554/eLife.05033.

119. Ganz, J., Shacham, T., Kramer, M., Shenkman, M., Eiger, H., Weinberg, N., Iancovici, O., Roy, S., Simhaev, L., Da’adoosh, B., et al. (2020). A novel specific PERK activator reduces toxicity and extends survival in Huntington’s disease models. Scientific Reports 10, 15, 6875. 10.1038/s41598-020-63899-4.

120. Dodson, M., Redmann, M., Rajasekaran, N.S., Darley-Usmar, V., and Zhang, J. (2015). KEAP1-NRF2 signalling and autophagy in protection against oxidative and reductive proteotoxicity. Biochem J 469, 347–355. 10.1042/bj20150568.

121. Kovac, S., Angelova, P.R., Holmström, K.M., Zhang, Y., Dinkova-Kostova, A.T., and Abramov, A.Y. (2015). Nrf2 regulates ROS production by mitochondria and NADPH oxidase. Biochim Biophys Acta 1850, 794–801. 10.1016/j.bbagen.2014.11.021.

122. Aquilano, K., Baldelli, S., and Ciriolo, M.R. (2014). Glutathione: new roles in redox signaling for an old antioxidant. Front. Pharmacol. 5. 10.3389/fphar.2014.00196.

123. Cybulsky, A.V. (2017). Endoplasmic reticulum stress, the unfolded protein response and autophagy in kidney diseases. Nature Reviews Nephrology 13, 681–696. 10.1038/nrneph.2017.129.

124. Rashid, H.-O., Yadav, R.K., Kim, H.-R., and Chae, H.-J. (2015). ER stress: Autophagy induction, inhibition and selection. Autophagy 11, 1956–1977. 10.1080/15548627.2015.1091141.

125. Settembre, C., Di Malta, C., Polito, V.A., Arencibia, M.G., Vetrini, F., Erdin, S., Erdin, S.U., Huynh, T., Medina, D., Colella, P., et al. (2011). TFEB Links Autophagy to Lysosomal Biogenesis. Science 332, 1429–1433. 10.1126/science.1204592.

126. Sardiello, M., Palmieri, M., di Ronza, A., Medina, D.L., Valenza, M., Gennarino, V.A., Di Malta, C., Donaudy, F., Embrione, V., Polishchuk, R.S., et al. (2009). A gene network regulating lysosomal biogenesis and function. Science 325, 473–477. 10.1126/science.1174447.

127. Palmieri, M., Impey, S., Kang, H., di Ronza, A., Pelz, C., Sardiello, M., and Ballabio, A. (2011). Characterization of the CLEAR network reveals an integrated control of cellular clearance pathways. Hum Mol Genet 20, 3852–3866. 10.1093/hmg/ddr306.

128. Consortium, T.P., coordination, O., Elsasser, S., Elia, L.P., Morimoto, R.I., Powers, E.T., group, H.M.S., Elsasser, S., Finley, D., University of California, S.F., et al. (2023). A Comprehensive Enumeration of the Human Proteostasis Network. 2. Components of the Autophagy-Lysosome Pathway. bioRxiv, 2023.2003.2022.533675. 10.1101/2023.03.22.533675.

129. Settembre, C., De Cegli, R., Mansueto, G., Saha, P.K., Vetrini, F., Visvikis, O., Huynh, T., Carissimo, A., Palmer, D., Jürgen Klisch, T., et al. (2013). TFEB controls cellular lipid metabolism through a starvation-induced autoregulatory loop. Nature Cell Biology 15, 647–658. 10.1038/ncb2718.

130. Grandjean, J.M.D., Plate, L., Morimoto, R.I., Bollong, M.J., Powers, E.T., and Wiseman, R.L. (2019). Deconvoluting Stress-Responsive Proteostasis Signaling Pathways for Pharmacologic Activation Using Targeted RNA Sequencing. Acs Chemical Biology 14, 784–795. 10.1021/acschembio.9b00134.

131. Sherman, B.T., Hao, M., Qiu, J., Jiao, X., Baseler, M.W., Lane, H.C., Imamichi, T., and Chang, W. (2022). DAVID: a web server for functional enrichment analysis and functional annotation of gene lists (2021 update). Nucleic Acids Research 50, W216–W221. 10.1093/nar/gkac194.

132. Martina, J.A., Chen, Y., Gucek, M., and Puertollano, R. (2012). MTORC1 functions as a transcriptional regulator of autophagy by preventing nuclear transport of TFEB. Autophagy 8, 903–914. 10.4161/auto.19653.

133. Roczniak-Ferguson, A., Petit, C.S., Froehlich, F., Qian, S., Ky, J., Angarola, B., Walther, T.C., and Ferguson, S.M. (2012). The transcription factor TFEB links mTORC1 signaling to transcriptional control of lysosome homeostasis. Sci Signal 5, ra42. 10.1126/scisignal.2002790.

134. Settembre, C., Zoncu, R., Medina, D.L., Vetrini, F., Erdin, S., Erdin, S., Huynh, T., Ferron, M., Karsenty, G., Vellard, M.C., et al. (2012). A lysosome-to-nucleus signalling mechanism senses and regulates the lysosome via mTOR and TFEB. Embo j 31, 1095–1108. 10.1038/emboj.2012.32.

135. Hara, K., Yonezawa, K., Weng, Q.-P., Kozlowski, M.T., Belham, C., and Avruch, J. (1998). Amino Acid Sufficiency and mTOR Regulate p70 S6 Kinase and eIF-4E BP1 through a Common Effector Mechanism*. Journal of Biological Chemistry 273, 14484–14494. 10.1074/jbc.273.23.14484.

136. Kim, J., Kundu, M., Viollet, B., and Guan, K.L. (2011). AMPK and mTOR regulate autophagy through direct phosphorylation of Ulk1. Nat Cell Biol. 13, 132–141. 10.1038/ncb2152.

137. Hara, K., Yonezawa, K., Kozlowski, M.T., Sugimoto, T., Andrabi, K., Weng, Q.P., Kasuga, M., Nishimoto, I., and Avruch, J. (1997). Regulation of eIF-4E BP1 phosphorylation by mTOR. The Journal of biological chemistry 272, 26457–26463.

138. Gingras, A.C., Raught, B., and Sonenberg, N. (2001). Regulation of translation initiation by FRAP/mTOR. Genes Dev 15, 807–826. 10.1101/gad.887201.

139. Wang, X., Beugnet, A., Murakami, M., Yamanaka, S., and Proud, C.G. (2005). Distinct signaling events downstream of mTOR cooperate to mediate the effects of amino acids and insulin on initiation factor 4E-binding proteins. Mol Cell Biol 25, 2558–2572. 10.1128/mcb.25.7.2558-2572.2005.

140. Sengupta, S., Peterson, T.R., and Sabatini, D.M. (2010). Regulation of the mTOR complex 1 pathway by nutrients, growth factors, and stress. Mol Cell 40, 310–322. 10.1016/j.molcel.2010.09.026.

141. Tsvetkov, A.S., Miller, J., Arrasate, M., Wong, J.S., Pleiss, M.A., and Finkbeiner, S. (2010). A small-molecule scaffold induces autophagy in primary neurons and protects against toxicity in a Huntington disease model. Proceedings of the National Academy of Sciences of the United States of America 107, 16982–16987. 10.1073/pnas.1004498107.

142. Sharma, P., Ando, D.M., Daub, A., Kaye, J.A., and Finkbeiner, S. (2012). HIGH-THROUGHPUT SCREENING IN PRIMARY NEURONS. In Imaging and Spectroscopic Analysis of Living Cells: Imaging Live Cells in Health and Disease, P.M. Conn, ed. pp. 331–360. 10.1016/b978-0-12-391856-7.00041-x.

143. Ghatak, S., Dolatabadi, N., Trudler, D., Zhang, X., Wu, Y., Mohata, M., Ambasudhan, R., Talantova, M., and Lipton, S.A. (2019). Mechanisms of hyperexcitability in Alzheimer’s disease hiPSC-derived neurons and cerebral organoids vs isogenic controls. eLife 8, e50333. 10.7554/eLife.50333.

144. Talantova, M., Sanz-Blasco, S., Zhang, X., Xia, P., Akhtar, M.W., Okamoto, S., Dziewczapolski, G., Nakamura, T., Cao, G., Pratt, A.E., et al. (2013). Aβ induces astrocytic glutamate release, extrasynaptic NMDA receptor activation, and synaptic loss. Proc Natl Acad Sci U S A 110, E2518–2527. 10.1073/pnas.1306832110.

145. Iwashita, H., Sakurai, H.T., Nagahora, N., Ishiyama, M., Shioji, K., Sasamoto, K., Okuma, K., Shimizu, S., and Ueno, Y. (2018). Small fluorescent molecules for monitoring autophagic flux. FEBS Letters 592, 559–567. 10.1002/1873-3468.12979.

146. Oh, C.K., Dolatabadi, N., Cieplak, P., Diaz-Meco, M.T., Moscat, J., Nolan, J.P., Nakamura, T., and Lipton, S.A. (2022). S-Nitrosylation of p62 Inhibits Autophagic Flux to Promote α-Synuclein Secretion and Spread in Parkinson’s Disease and Lewy Body Dementia. J Neurosci 42, 3011–3024. 10.1523/jneurosci.1508-21.2022.

147. Bruch, J., Xu, H., Rosler, T.W., De Andrade, A., Kuhn, P.H., Lichtenthaler, S.F., Arzberger, T., Winklhofer, K.F., Muller, U., and Huglinger, G.U. (2017). PERK activation mitigates tau pathology in vitro and in vivo. Embo Molecular Medicine 9, 371–384. 10.15252/emmm.201606664.

148. Silva, M.C., Nandi, G.A., Tentarelli, S., Gurrell, I.K., Jamier, T., Lucente, D., Dickerson, B.C., Brown, D.G., Brandon, N.J., and Haggarty, S.J. (2020). Prolonged tau clearance and stress vulnerability rescue by pharmacological activation of autophagy in tauopathy neurons. Nat Commun 11, 3258. 10.1038/s41467-020-16984-1.

149. Silva, M.C., Nandi, G., Donovan, K.A., Cai, Q., Berry, B.C., Nowak, R.P., Fischer, E.S., Gray, N.S., Ferguson, F.M., and Haggarty, S.J. (2022). Discovery and Optimization of Tau Targeted Protein Degraders Enabled by Patient Induced Pluripotent Stem Cells-Derived Neuronal Models of Tauopathy. Front Cell Neurosci 16, 801179. 10.3389/fncel.2022.801179.

150. Silva, M.C., Ferguson, F.M., Cai, Q., Donovan, K.A., Nandi, G., Patnaik, D., Zhang, T., Huang, H.T., Lucente, D.E., Dickerson, B.C., et al. (2019). Targeted degradation of aberrant tau in frontotemporal dementia patient-derived neuronal cell models. Elife 8. 10.7554/eLife.45457.

151. Guo, J.L., and Lee, V.M. (2011). Seeding of normal Tau by pathological Tau conformers drives pathogenesis of Alzheimer-like tangles. J Biol Chem 286, 15317–15331. 10.1074/jbc.M110.209296.

152. Kfoury, N., Holmes, B.B., Jiang, H., Holtzman, D.M., and Diamond, M.I. (2012). Trans-cellular propagation of Tau aggregation by fibrillar species. J Biol Chem 287, 19440–19451. 10.1074/jbc.M112.346072.

153. Silva, M.C., Cheng, C., Mair, W., Almeida, S., Fong, H., Biswas, M. Helal U., Zhang, Z., Huang, Y., Temple, S., Coppola, G., et al. (2016). Human iPSC-Derived Neuronal Model of Tau-A152T Frontotemporal Dementia Reveals Tau-Mediated Mechanisms of Neuronal Vulnerability. Stem Cell Reports 7, 325–340. 10.1016/j.stemcr.2016.08.001.

154. Vierbuchen, T., Ostermeier, A., Pang, Z.P., Kokubu, Y., Südhof, T.C., and Wernig, M. (2010). Direct conversion of fibroblasts to functional neurons by defined factors. Nature 463, 1035–1041. 10.1038/nature08797.

155. Pang, Z.P., Yang, N., Vierbuchen, T., Ostermeier, A., Fuentes, D.R., Yang, T.Q., Citri, A., Sebastiano, V., Marro, S., Südhof, T.C., and Wernig, M. (2011). Induction of human neuronal cells by defined transcription factors. Nature 476, 220–223. 10.1038/nature10202.

156. Mertens, J., Herdy, J.R., Traxler, L., Schafer, S.T., Schlachetzki, J.C.M., Böhnke, L., Reid, D.A., Lee, H., Zangwill, D., Fernandes, D.P., et al. (2021). Age-dependent instability of mature neuronal fate in induced neurons from Alzheimer’s patients. Cell Stem Cell 28, 1533–1548.e1536. 10.1016/j.stem.2021.04.004.

157. Mertens, J., Paquola, A.C.M., Ku, M., Hatch, E., Böhnke, L., Ladjevardi, S., McGrath, S., Campbell, B., Lee, H., Herdy, J.R., et al. (2015). Directly Reprogrammed Human Neurons Retain Aging-Associated Transcriptomic Signatures and Reveal Age-Related Nucleocytoplasmic Defects. Cell Stem Cell 17, 705–718. 10.1016/j.stem.2015.09.001.

158. Chou, C.C., Vest, R., Prado, M.A., Wilson-Grady, J., Paulo, J.A., Shibuya, Y., Moran-Losada, P., Lee, T.T., Luo, J., Gygi, S.P., et al. (2023). Proteostasis and lysosomal quality control deficits in Alzheimer’s disease neurons. bioRxiv. 10.1101/2023.03.27.534444.

159. Stokin, G.B., Lillo, C., Falzone, T.L., Brusch, R.G., Rockenstein, E., Mount, S.L., Raman, R., Davies, P., Masliah, E., Williams, D.S., and Goldstein, L.S.B. (2005). Axonopathy and transport deficits early in the pathogenesis of Alzheimer’s disease. Science 307, 1282–1288. 10.1126/science.1105681.

160. Encalada, S.E., and Goldstein, L.S. (2014). Biophysical challenges to axonal transport: motor-cargo deficiencies and neurodegeneration. Annu Rev Biophys 43, 141–169. 10.1146/annurev-biophys-051013-022746.

161. Zanusso, G., Polo, A., Farinazzo, A., Nonno, R., Cardone, F., Di Bari, M., Ferrari, S., Principe, S., Gelati, M., Fasoli, E., et al. (2007). Novel prion protein conformation and glycotype in Creutzfeldt-Jakob disease. Arch Neurol 64, 595–599. 10.1001/archneur.64.4.595.

162. Sikorska, B., Liberski, P.P., Giraud, P., Kopp, N., and Brown, P. (2004). Autophagy is a part of ultrastructural synaptic pathology in Creutzfeldt-Jakob disease: a brain biopsy study. Int J Biochem Cell Biol 36, 2563–2573. 10.1016/j.biocel.2004.04.014.

163. Lee, S., Sato, Y., and Nixon, R.A. (2011). Primary lysosomal dysfunction causes cargo-specific deficits of axonal transport leading to Alzheimer-like neuritic dystrophy. Autophagy 7, 1562–1563. 10.4161/auto.7.12.17956.

164. Nixon, R.A. (2017). Amyloid precursor protein and endosomal-lysosomal dysfunction in Alzheimer’s disease: inseparable partners in a multifactorial disease. Faseb j 31, 2729–2743. 10.1096/fj.201700359.

165. Chaiamarit, T., Verhelle, A., Chassefeyre, R., Shukla, N., Novak, S.W., Andrade, L.R., Manor, U., and Encalada, S.E. (2023). Mutant Prion Protein Endoggresomes are Hubs for Local Axonal Organelle-Cytoskeletal Remodeling. bioRxiv, 2023.2003.2019.533383. 10.1101/2023.03.19.533383.

166. Vital, A., Laplanche, J.L., Bastard, J.R., Xiao, X., Zou, W.Q., and Vital, C. (2011). A case of Gerstmann-Sträussler-Scheinker disease with a novel six octapeptide repeat insertion. Neuropathol Appl Neurobiol 37, 554–559. 10.1111/j.1365-2990.2011.01174.x.

167. Chassefeyre, R., Chaiamarit, T., Verhelle, A., Novak, S.W., Andrade, L.R., Leitao, A.D.G., Manor, U., and Encalada, S.E. (2021). Endosomal sorting drives the formation of axonal prion protein endoggresomes. Sci. Adv. 7, 20, eabg3693. 10.1126/sciadv.abg3693.

168. Chin, M.Y., Patwardhan, A.R., Ang, K.-H., Wang, A.L., Alquezar, C., Welch, M., Nguyen, P.T., Grabe, M., Molofsky, A.V., Arkin, M.R., and Kao, A.W. (2021). Genetically Encoded, pH-Sensitive mTFP1 Biosensor for Probing Lysosomal pH. ACS Sens. 6, 2168–2180. 10.1021/acssensors.0c02318.

169. Neumann, S., Chassefeyre, R., Campbell, G.E., and Encalada, S.E. (2017). KymoAnalyzer: a software tool for the quantitative analysis of intracellular transport in neurons. Traffic 18, 71–88. 10.1111/tra.12456.

170. Aman, Y., Schmauck-Medina, T., Hansen, M., Morimoto, R.I., Simon, A.K., Bjedov, I., Palikaras, K., Simonsen, A., Johansen, T., Tavernarakis, N., et al. (2021). Autophagy in healthy aging and disease. Nature Aging 1, 634–650. 10.1038/s43587-021-00098-4.

171. Mizushima, N., and Levine, B. (2020). Autophagy in Human Diseases. New England Journal of Medicine 383, 1564–1576. 10.1056/NEJMra2022774.

172. Klionsky, D.J., Petroni, G., Amaravadi, R.K., Baehrecke, E.H., Ballabio, A., Boya, P., Bravo-San Pedro, J.M., Cadwell, K., Cecconi, F., Choi, A.M.K., et al. (2021). Autophagy in major human diseases. Embo j 40, e108863. 10.15252/embj.2021108863.

173. van Beek, N., Klionsky, D.J., and Reggiori, F. (2018). Genetic aberrations in macroautophagy genes leading to diseases. Biochim Biophys Acta Mol Cell Res 1865, 803–816. 10.1016/j.bbamcr.2018.03.002.

174. Johannesson, M., Sahlin, C., Söderberg, L., Basun, H., Fälting, J., Möller, C., Zachrisson, O., Sunnemark, D., Svensson, A., Odergren, T., and Lannfelt, L. (2021). Elevated soluble amyloid beta protofibrils in Down syndrome and Alzheimer’s disease. Mol Cell Neurosci 114, 103641. 10.1016/j.mcn.2021.103641.

175. Tolar, M., Hey, J., Power, A., and Abushakra, S. (2021). Neurotoxic Soluble Amyloid Oligomers Drive Alzheimer’s Pathogenesis and Represent a Clinically Validated Target for Slowing Disease Progression. International Journal of Molecular Sciences 22, 6355.

176. Shi, M., Chu, F., Zhu, F., and Zhu, J. (2022). Impact of Anti-amyloid-β Monoclonal Antibodies on the Pathology and Clinical Profile of Alzheimer’s Disease: A Focus on Aducanumab and Lecanemab. Frontiers in Aging Neuroscience 14. 10.3389/fnagi.2022.870517.

177. Mawuenyega, K.G., Sigurdson, W., Ovod, V., Munsell, L., Kasten, T., Morris, J.C., Yarasheski, K.E., and Bateman, R.J. (2010). Decreased Clearance of CNS β-Amyloid in Alzheimer’s Disease. Science 330, 1774–1774. doi:10.1126/science.1197623.

178. Correia, S.C., Resende, R., Moreira, P.I., and Pereira, C.M. (2015). Alzheimer’s disease-related misfolded proteins and dysfunctional organelles on autophagy menu. DNA Cell Biol 34, 261–273. 10.1089/dna.2014.2757.

179. Chiang, W.C., Wei, Y., Kuo, Y.C., Wei, S., Zhou, A., Zou, Z., Yehl, J., Ranaghan, M.J., Skepner, A., Bittker, J.A., et al. (2018). High-Throughput Screens To Identify Autophagy Inducers That Function by Disrupting Beclin 1/Bcl-2 Binding. ACS Chem Biol 13, 2247–2260. 10.1021/acschembio.8b00421.

180. Zhang, W., Li, X., Wang, S., Chen, Y., and Liu, H. (2020). Regulation of TFEB activity and its potential as a therapeutic target against kidney diseases. Cell Death Discovery 6, 32. 10.1038/s41420-020-0265-4.

181. Napolitano, G., Di Malta, C., and Ballabio, A. (2022). Non-canonical mTORC1 signaling at the lysosome. Trends in Cell Biology 32, 920–931. 10.1016/j.tcb.2022.04.012.

182. Vega-Rubin-de-Celis, S., Peña-Llopis, S., Konda, M., and Brugarolas, J. (2017). Multistep regulation of TFEB by MTORC1. Autophagy 13, 464–472. 10.1080/15548627.2016.1271514.

183. Hardy, J., and Selkoe, D.J. (2002). Medicine - The amyloid hypothesis of Alzheimer’s disease: Progress and problems on the road to therapeutics. Science 297, 353–356. 10.1126/science.1072994.

184. Holtzman, D.M., Morris, J.C., and Goate, A.M. (2011). Alzheimer’s Disease: The Challenge of the Second Century. Science Translational Medicine 3, 77sr1. 10.1126/scitranslmed.3002369.

185. Wolfe, D.M., Lee, J.H., Kumar, A., Lee, S., Orenstein, S.J., and Nixon, R.A. (2013). Autophagy failure in Alzheimer’s disease and the role of defective lysosomal acidification. European Journal of Neuroscience 37, 1949–1961. 10.1111/ejn.12169.

186. Boland, B., Kumar, A., Lee, S., Platt, F.M., Wegiel, J., Yu, W.H., and Nixon, R.A. (2008). Autophagy induction and autophagosome clearance in neurons: Relationship to autophagic pathology in Alzheimer’s disease. Journal of Neuroscience 28, 6926–6937. 10.1523/jneurosci.0800-08.2008.

187. Podlesny-Drabiniok, A., Marcora, E., and Goate, A.M. (2020). Microglial Phagocytosis: A Disease-Associated Process Emerging from Alzheimer’s Disease Genetics. Trends in Neurosciences 43, 965–979. 10.1016/j.tins.2020.10.002.

188. Seo, J.Y., Lim, S.S., Kim, J., Lee, K.W., and Kim, J.-S. (2017). Alantolactone and Isoalantolactone Prevent Amyloid β25–35-induced Toxicity in Mouse Cortical Neurons and Scopolamine-induced Cognitive Impairment in Mice. Phytotherapy Research 31, 801–811. 10.1002/ptr.5804.

189. Encalada, S.E., Szpankowski, L., Xia, C.H., and Goldstein, L.S.B. (2011). Stable Kinesin and Dynein Assemblies Drive the Axonal Transport of Mammalian Prion Protein Vesicles. Cell 144, 551–565. 10.1016/j.cell.2011.01.021.

190. Listenberger, L.L., and Brown, D.A. (2007). Fluorescent Detection of Lipid Droplets and Associated Proteins. Current Protocols in Cell Biology 35, 24.22.21–24.22.11. 10.1002/0471143030.cb2402s35.

191. Bolte, S., and Cordelières, F.P. (2006). A guided tour into subcellular colocalization analysis in light microscopy. Journal of Microscopy 224, 213–232. 10.1111/j.1365-2818.2006.01706.x.

192. Barmada, S.J., Serio, A., Arjun, A., Bilican, B., Daub, A., Ando, D.M., Tsvetkov, A., Pleiss, M., Li, X., Peisach, D., et al. (2014). Autophagy induction enhances TDP43 turnover and survival in neuronal ALS models. Nat Chem Biol 10, 677–685. 10.1038/nchembio.1563.

193. Johnson, T.J. (1987). Glutaraldehyde fixation chemistry: oxygen-consuming reactions. Eur J Cell Biol 45, 160–169.

194. Paxman, R., Plate, L., Blackwood, E.A., Glembotski, C., Powers, E.T., Wiseman, R.L., and Kelly, J.W. (2018). Pharmacologic ATF6 activating compounds are metabolically activated to selectively modify endoplasmic reticulum proteins. eLife 7, e37168. 10.7554/eLife.37168.

195. Grandjean, J.M.D., Madhavan, A., Cech, L., Seguinot, B.O., Paxman, R.J., Smith, E., Scampavia, L., Powers, E.T., Cooley, C.B., Plate, L., et al. (2020). Pharmacologic IRE1/XBP1s activation confers targeted ER proteostasis reprogramming. Nature Chemical Biology 16, 1052-+. 10.1038/s41589-020-0584-z.

196. Napolitano, G., Esposito, A., Choi, H., Matarese, M., Benedetti, V., Di Malta, C., Monfregola, J., Medina, D.L., Lippincott-Schwartz, J., and Ballabio, A. (2018). mTOR-dependent phosphorylation controls TFEB nuclear export. Nature Communications 9, 3312. 10.1038/s41467-018-05862-6.

