## Supplemental Figures and Legends for "An mTOR-independent Macroautophagy Activator Ameliorates Tauopathy and Prionopathy Neurodegeneration Phenotypes"

**Supplemental Figure Legends.**

**Figure S1**: (**A**) Western blot of SW1990 cells either pretreated in RPMI media for 24 h with oleic acid (10µM, 30µM or 100µM) followed by 20-h treatment with 0.1% DMSO, or not pretreated with oleic acid but treated with 20 h the next day with 0.1% DMSO, 250nM rapamycin, 5µM AZD8055 (mTOR inhibitor) or 20µM chloroquine. (See treatment timeline above western blot.) (**B**) Total lipid droplet area (visualized by BODIPY staining) in SW1990 cells treated with known autophagy activators (50nM rapamycin or 10µM tamoxifen) or 0.1% DMSO control for 8, 16, or 24 h in the presence of 50µM oleic acid. * Dunnett's multiple comparisons test following one-way ANOVA $p < .05$, ** $p < .01$, *** $p < .001$, **** $p < .0001$. Bars and error bars depict mean + SEM, n = 8. (**C**) Lipid droplet degradation and LC3-II band intensity quantified from western blot in dose-response curves for PI-103, PP242, and tamoxifen treatment of SW1990 cells pretreated with 10µM oleic acid for 24 h followed by treatment with compound for 20 h. Points and error bars depict mean ± SEM, n = 3. (**D**) Relative lipid droplet fluorescence (BODIPY 493/503) area per SW1990, HeLa, U251, or HT-1080 cell treated with 100nM rapamycin, 1µM PI-103, 1µM PP242, or DMSO control (normalized to DMSO control) for 20 h in the presence of 10µM oleic acid. Bars and error bars depict mean ± SEM, n = 3. (**E**) Normalized lipid droplet clearance data (candidate autophagy activator concentration range: 123nM-10µM) from SW1990, U251, and HT-1080 cells fed with 10µM oleic acid followed by treatment with hit compounds. The heat map tiles of CCT020312 are surrounded by a black rectangle identified by an arrow. (**F**) Representative data for control compounds (DMSO, Torin-1) of the autophagy flux screen and the formula used herein to calculate the percentage of autophagy flux induction by example compound (2µM CCT020312) for each time point. (**G**) Example categories of hit compounds of the autophagy flux assay based on the flux pattern over time. CCT020312 is an example compound within category 3.

**Figure S2**: (**A**) CellTiter-Glo cell viability assay performed after 24-h CCT020312 dose-responsive treatment in SW1990 and U251 cells (n = 8 per cell line). 50% cell viability at 13.0µM CCT020312 in SW1990 cells and 11.5µM CCT020312 in U251 cells. (**B**) Growth curves corresponding to Figure 2D. (**C**) Representative confocal images of HeLa cells stained with LAMP2, LC3B, and Hoechst after 24-h treatment with 250nM rapamycin, 1.25µM or 2.5µM CCT020312, or DMSO. Lower set of images are same respective treatments but also treated with 200nM Bafilomycin A1 (BafA1) for the final 4 h of treatment. Exposure has been increased to improve contrast in this figure. LC3B stain is in red, LAMP2 stain is in green, and Hoescht stain is in blue. (**D**) Quantification of Pearson's Coefficient for colocalization of LC3B and LAMP2 from (**C**).

**Figure S3**: (**A**) Volcano plot depicting RNA-Seq analysis of HEK293T cells treated with 1µM CCT020312 for 4 h. Horizontal dotted line denotes adjusted p-value = 0.05. HSPA1A and HSPA1B have nearly identical adjusted p-value and fold change so therefore appear as one point. (**B**) Gene set enrichment analysis of adjacent RNA-Seq data for activation of stress-responsive signaling pathways: PERK, ATF6, and IRE1/XBP1s arms of the unfolded protein response; heat shock response (HSR); oxidative stress response (OSR); and other genes as described in Table 1 of {Grandjean et al, 2019}. (**C**) Bar graph depicting the activation of the ATF4.Fluc ATF4 transcriptional reporter in HEK293T cells treated for 18 h with vehicle (0.1% DMSO), thapsigargin (500nM), or CCT020312 (1µM). Error bars show SEM for three biological replicates. The p-value for Dunnett's multiple comparisons test to DMSO control following one-way ANOVA has been written above the CCT020312 bar. (**D**) Total lipid droplet fluorescence (BODIPY 493/503) area in U251 cells treated with CCT020312 at varying doses in combination with PERK inhibitor GSK2656157 (3μM or 10μM) or DMSO control for 24 h. Error bars show SEM for three biological replicates. (**E**) Immunoblots of U251 cells treated with CCT020312, rapamycin, PI-103, or vehicle for 20 h and co-treated with ISRIB or DMSO control for the final 2 h of treatment. (**F**) Immunoblots of $PERK^{+/+}$ and $PERK^{-/-}$ MEF cells treated with CCT020312, rapamycin, PI-103, or vehicle for 20 h. (**G**) Relative ARE-LUC luminescence measurements from IMR32 cells treated for 24 h with a concentration response of CCT020312 and reported NRF2 activator bardoxolone (n = 3 biologically independent samples; mean and SEM). Lower schematic depicts NRF2-ARE-Luciferase reporter construct. (**H**) Bar graph depicting the reduced (GSH) to oxidized (GSSG) glutathione ratio in HEK293T cells treated for 2 h with vehicle (0.1% DMSO), radical oxidative stress inducer menadione (40μM) or CCT020312 (1μM, 5μM or 10μM). Error bars show SEM for three technical replicates. The p-value for Dunnett's multiple comparisons test to DMSO control following one-way ANOVA has been written above the 1μM CCT020312 bar. (**I**) Bar graph depicting the activation of the ERSE.FLuc ATF6 transcriptional reporter in HEK293T cells treated for 18 h with vehicle (Veh, 0.1% DMSO), thapsigargin (Tg, 500nM), tunicamycin (Tm, 1 mg/mL), AA147 (10μM), rapamycin (250nM), PI-103 (1μM), or CCT020312 (1μM). Error bars show SEM for three technical replicates. (**J**) Bar graph depicting the activation of the XBP1.RLuc IRE1 transcriptional reporter in HEK293T cells treated for 18 h with vehicle (Veh, 0.1% DMSO), thapsigargin (500nM), IXA4 (10μM), or CCT020312 (10μM). Error bars show SEM for three biological replicates. (**K**) Bar graph depicting the activation of the HSF1.Fluc HSF1 transcriptional reporter in HEK293T cells treated for 18 h with vehicle (0.1% DMSO), MG132 (10μM), or CCT020312 (10μM). Error bars show SEM for three biological replicates. (**L**) Volcano plot depicting RNA-Seq analysis of HeLa cells treated with 2.5μM CCT020312 for 20 h. Genes with fold change less than 0.5 are colored blue and genes with fold change greater than 2 are colored red. Some of the most significant genes are labeled on the plot and the full gene list can be found in Supplemental Table 1. Horizontal dotted line denotes adjusted p-value = 0.01. (**M**) Volcano plot depicting upregulated genes from RNA-Seq analysis of HeLa cells treated with 2.5μM CCT020312 for 20 h. Genes with fold change greater than 1.25 are colored orange, and genes with fold change greater than 2 are colored red. Some genes related to lipid metabolism are labeled on the plot. Horizontal dotted line denotes adjusted p-value = 0.01.

**Figure S4:** *CCT020312 activation of autophagy in patient iPSC-derived tau-P301L neurons.* (**A**) Western blot analysis of CCT020312 concentration effect on autophagy-lysosomal markers LC3-II, LAMP1, p62/SQSTM1 and cathepsin D (CTSD) of FTD tau-P301L neurons at 6 weeks of differentiation after 24-h treatment. LC3-II arrow indicates the lower band (upper band is LC3-I) that was used for densitometry quantification. (**B, C**) Western blot densitometry, quantification of bands' pixel intensity (in arbitrary units, a.u.). Data points represent mean densitometry normalized to actin and relative to vehicle (DMSO) ± SD (n = 2). **(D)** Western blot analysis of rapamycin concentration effect on autophagy-lysosomal markers (LC3-II, LAMP1, p62/SQSTM1) of 6-week differentiated FTD neurons, treated for 24 h and densitometry quantification. Note that above 5μM, rapamycin is toxic to iPSC-neurons.

**Figure S7**: (**A**) Top panels: Schematic of $PrP^{WT}$-mCh and $PrP^{PG14}$-mCh constructs (top). SS: signal sequence, GPI: GPI anchor. The location of the human 3F4 epitope is shown in cyan. Bottom panels: Representative fluorescence images of axons of neurons expressing $PrP^{WT}$-mCh and $PrP^{PG14}$-mCh. Arrows indicate $PrP^{WT}$-mCh vesicles (endolysosomes), and arrowheads indicate $PrP^{PG14}$-mCh endoggresomes. Inverted images shown. (**B**) Left panels: representative fluorescence images of axons of neurons expressing $PrP^{WT}$-mCh or $PrP^{PG14}$-mCh and treated with carboxy CCT or DMSO (vehicle). Inverted images shown. Right panel: endoggresome density quantification of axons in (B) showing no statistically significant decrease in axonal endoggresome density. (**C**) Left panels: Viability of 7 DIV hippocampal neurons treated with DMSO (vehicle), 200nM CCT020312, or 200nM carboxy CCT. Inverted images shown. Right panels: quantification of Nuc-Green positive dead neurons. ns = not statistically significant. (**D**) Left panels: representative fluorescence images of axons of neurons expressing $PrP^{WT}$-mCh or $PrP^{PG14}$-mCh and treated with rapamycin or DMSO (vehicle). Inverted images shown. Right: endoggresome density quantification showing a statistically significant, dose-dependent decrease in endoggresome density starting at 50nM rapamycin. * $p < .05$, ** $p < .01$, *** $p < .001$, **** $p < .0001$, ns = not statistically significant. One-way ANOVA, Šidák correction. (**E**) Cumulative distributions functions (CDFs) of anterograde and retrograde segmental velocities (top) and segmental run lengths (bottom) from $PrP^{WT}$-mCh or $PrP^{PG14}$-mCh vesicles moving in axons expressing $PrP^{WT}$-mCh or $PrP^{PG14}$-mCh, respectively, and treated with CCT020312 or DMSO (vehicle). * Kolmogorov-Smirnov * $p < .05$, ** $p < .01$, *** $p < .001$, ns = not statistically significant.

**Data S2**: (**A**) Representative transmission electron microscopy images of HeLa cells treated for indicated times with indicated treatments. High magnification image depicts the boxed area in the low magnification image. Scale bar of low magnification images, 5 µm; scale bar of high magnification images, 2 µm.

### Figure S1

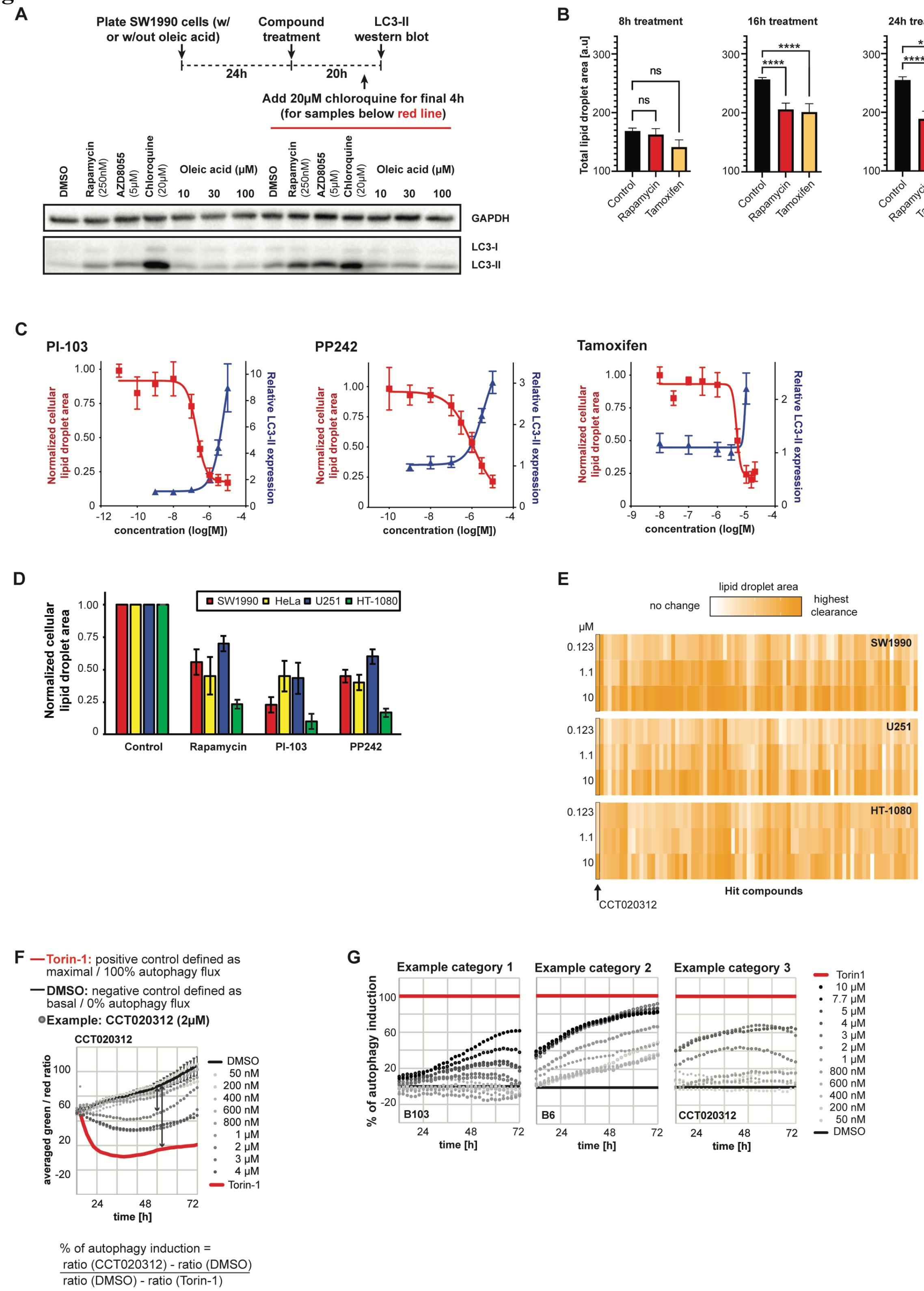
A
Plate SW1990 cells (w/ or w/out oleic acid)
Compound treatment
LC3-II western blot
24h
20h
Add 20μM chloroquine for final 4h (for samples below red line)
DMSO
Rapamycin (250nM)
AZD8055 (5μM)
Chloroquine (20μM)
Oleic acid (μM)
10
30
100
GAPDH
LC3-I
LC3-II
B
8h treatment
16h treatment
24h treatment
Total lipid droplet area [a.u]
ns
****
Control
Rapamycin
Tamoxifen
C
PI-103
PP242
Tamoxifen
Normalized cellular lipid droplet area
Relative LC3-II expression
concentration (log[M])
D
SW1990
HeLa
U251
HT-1080
Normalized cellular lipid droplet area
Control
Rapamycin
PI-103
PP242
E
lipid droplet area
no change
highest clearance
μM
0.123
1.1
10
SW1990
U251
HT-1080
CCT020312
Hit compounds
F
—Torin-1: positive control defined as maximal / 100% autophagy flux
—DMSO: negative control defined as basal / 0% autophagy flux
Example: CCT020312 (2μM)
CCT020312
averaged green / red ratio
time [h]
DMSO
50 nM
200 nM
400 nM
600 nM
800 nM
1 μM
2 μM
3 μM
4 μM
Torin-1
% of autophagy induction =
ratio (CCT020312) - ratio (DMSO)
ratio (DMSO) - ratio (Torin-1)
G
Example category 1
Example category 2
Example category 3
% of autophagy induction
B103
B6
CCT020312
time [h]
Torin1
10 μM
7.7 μM
5 μM
4 μM
3 μM
2 μM
1 μM
800 nM
600 nM
400 nM
200 nM
50 nM
DMSO

### Figure S2

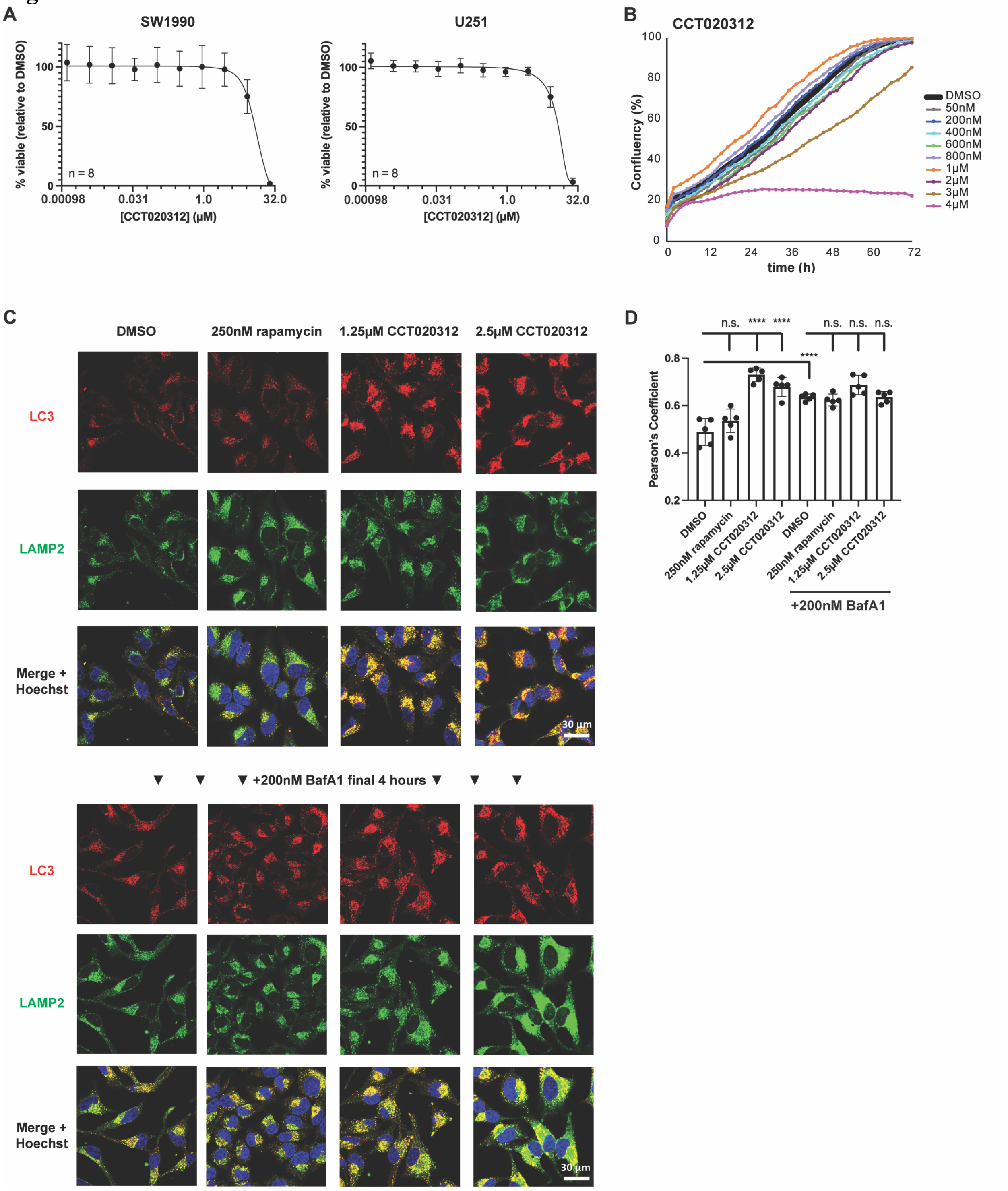
A
SW1990
U251
% viable (relative to DMSO)
n = 8
0.00098
0.031
1.0
32.0
[CCT020312] (µM)
B
CCT020312
Confluency (%)
time (h)
DMSO
50nM
200nM
400nM
600nM
800nM
1µM
2µM
3µM
4µM
C
DMSO
250nM rapamycin
1.25µM CCT020312
2.5µM CCT020312
LC3
LAMP2
Merge + Hoechst
30 µm
+200nM BafA1 final 4 hours
D
Pearson's Coefficient
n.s.
****
DMSO
250nM rapamycin
1.25µM CCT020312
2.5µM CCT020312
+200nM BafA1

### Figure S3

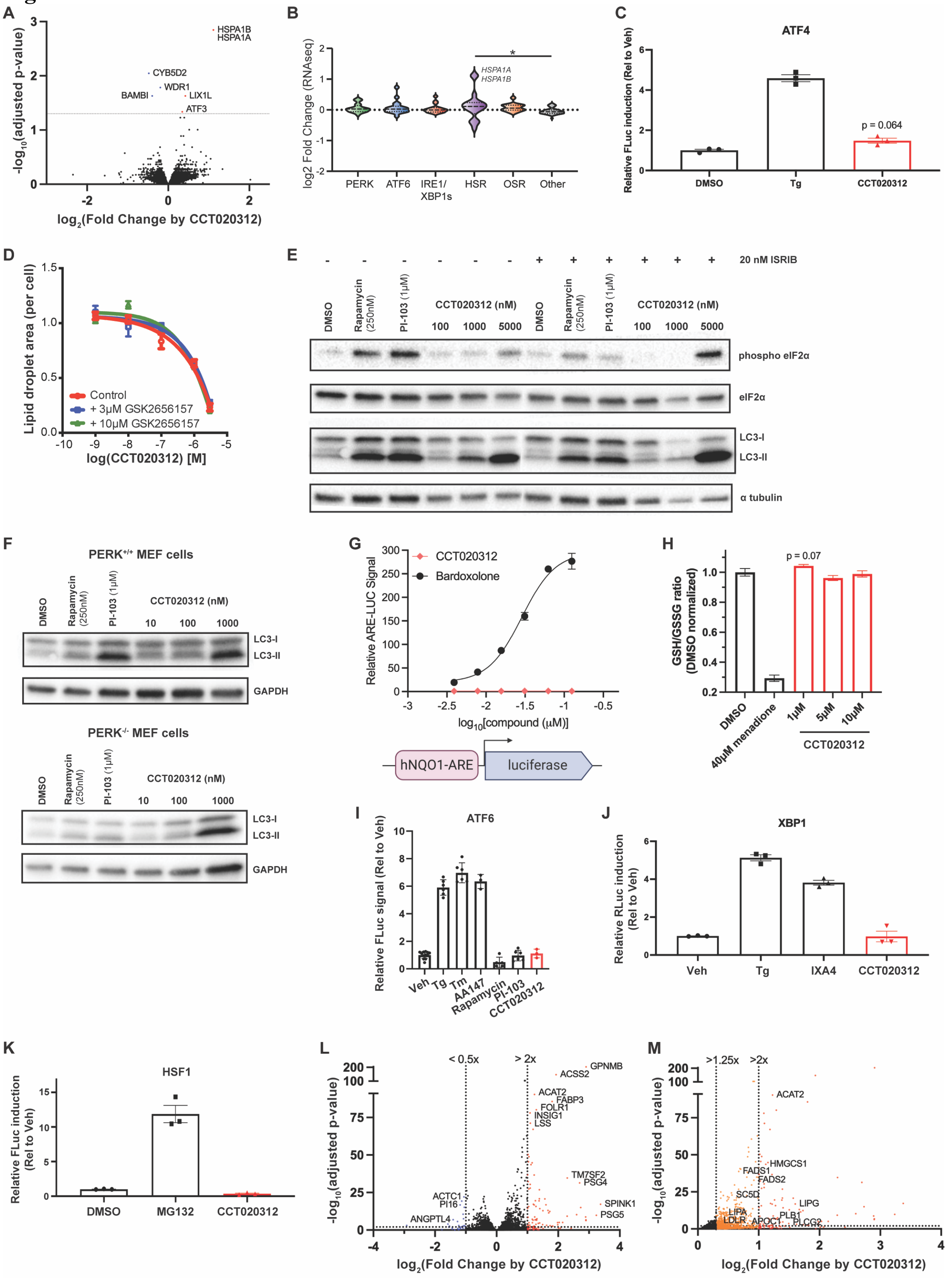

A
-log10(adjusted p-value)
log2(Fold Change by CCT020312)
HSPA1B
HSPA1A
CYB5D2
WDR1
BAMBI
LIX1L
ATF3
B
log2 Fold Change (RNAseq)
PERK
ATF6
IRE1/
XBP1s
HSR
OSR
Other
HSPA1A
HSPA1B
C
ATF4
Relative FLuc induction (Rel to Veh)
DMSO
Tg
CCT020312
p = 0.064
D
Lipid droplet area (per cell)
log(CCT020312) [M]
Control
+ 3μM GSK2656157
+ 10μM GSK2656157
E
20 nM ISRIB
DMSO
Rapamycin (250nM)
PI-103 (1μM)
CCT020312 (nM)
100
1000
5000
phospho eIF2α
eIF2α
LC3-I
LC3-II
α tubulin
F
PERK+/+ MEF cells
PERK-/- MEF cells
CCT020312 (nM)
10
100
1000
LC3-I
LC3-II
GAPDH
G
Relative ARE-LUC Signal
CCT020312
Bardoxolone
log10[compound (μM)]
hNQO1-ARE
luciferase
H
GSH/GSSG ratio (DMSO normalized)
p = 0.07
DMSO
40μM menadione
1μM
5μM
10μM
CCT020312
I
ATF6
Relative FLuc signal (Rel to Veh)
Veh
Tg
Tm
AA147
Rapamycin
PI-103
CCT020312
J
XBP1
Relative RLuc induction (Rel to Veh)
Veh
Tg
IXA4
CCT020312
K
HSF1
Relative FLuc induction (Rel to Veh)
DMSO
MG132
CCT020312
L
-log10(adjusted p-value)
< 0.5x
> 2x
GPNMB
ACSS2
ACAT2
FABP3
FOLR1
INSIG1
LSS
TM7SF2
PSG4
SPINK1
PSG5
ACTC1
PI16
ANGPTL4
log2(Fold Change by CCT020312)
M
-log10(adjusted p-value)
>1.25x
>2x
ACAT2
HMGCS1
FADS1
FADS2
SC5D
LIPG
LIPA
LDLR
APOC1
PLB1
PLCG2
log2(Fold Change by CCT020312)

### Figure S4

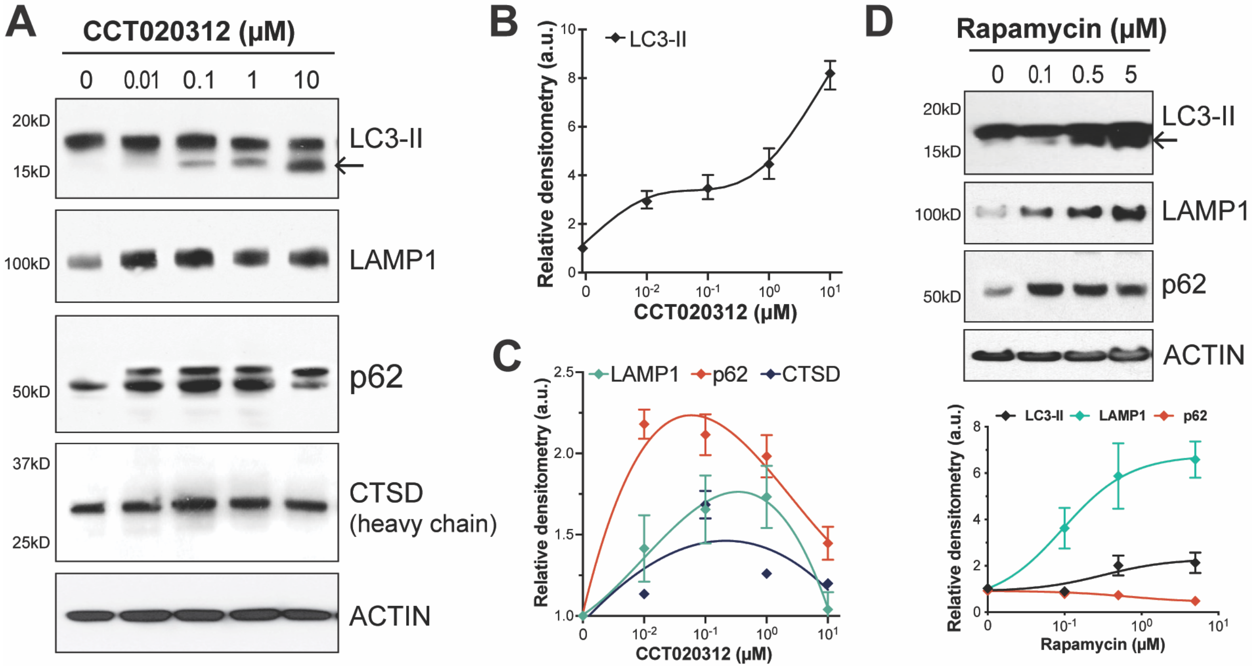
A
CCT020312 (μM)
0 0.01 0.1 1 10
20kD
15kD
LC3-II
100kD
LAMP1
50kD
p62
37kD
25kD
CTSD
(heavy chain)
ACTIN
B
LC3-II
Relative densitometry (a.u.)
CCT020312 (μM)
C
LAMP1 p62 CTSD
Relative densitometry (a.u.)
CCT020312 (μM)
D
Rapamycin (μM)
0 0.1 0.5 5
20kD
15kD
LC3-II
100kD
LAMP1
50kD
p62
ACTIN
LC3-II LAMP1 p62
Relative densitometry (a.u.)
Rapamycin (μM)

### Figure S5

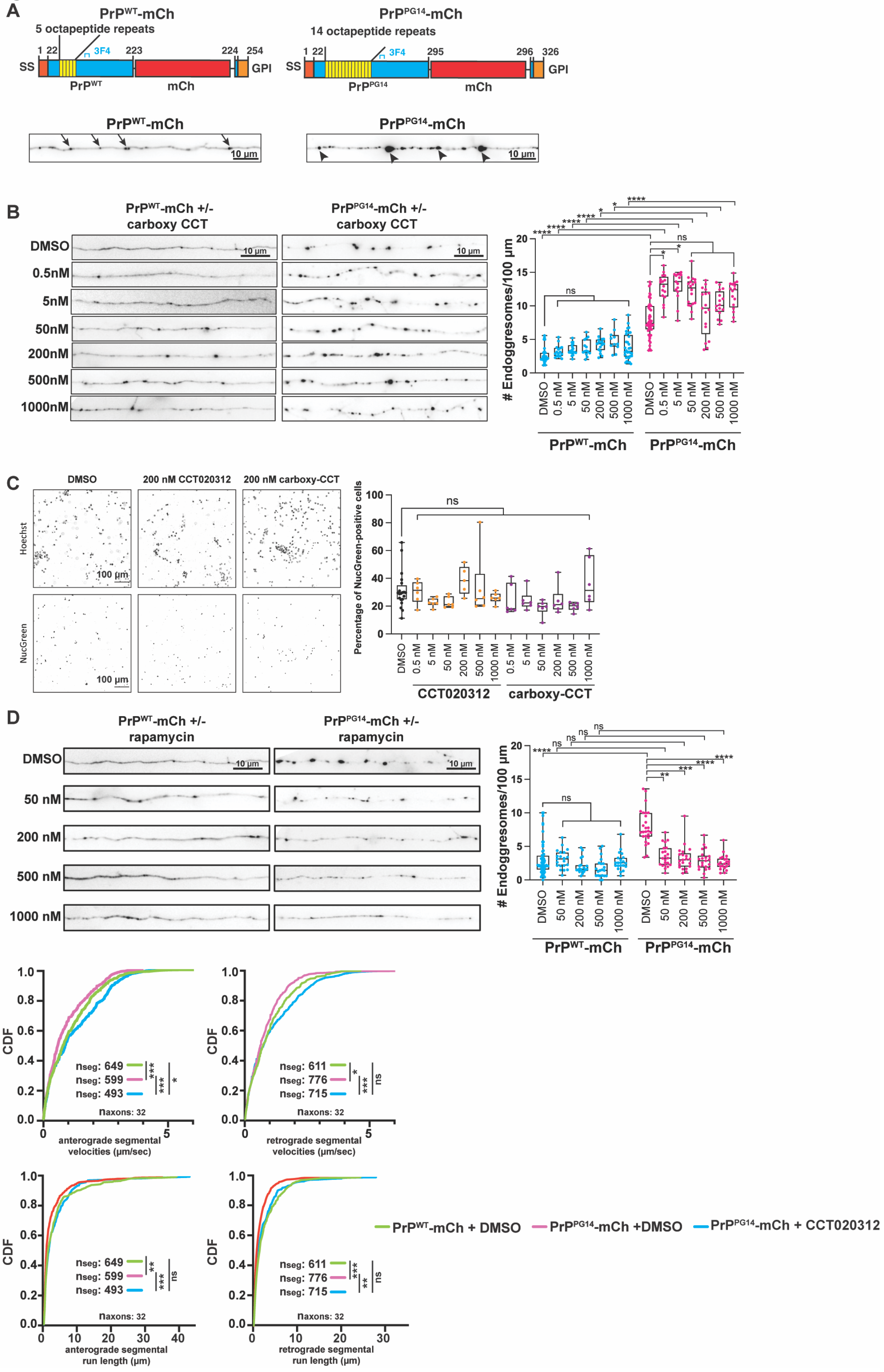
A
PrPWT-mCh
5 octapeptide repeats
3F4
SS
GPI
PrPWT
mCh
PrPPG14-mCh
14 octapeptide repeats
PrPPG14
10 µm
B
PrPWT-mCh +/- carboxy CCT
PrPPG14-mCh +/- carboxy CCT
DMSO
0.5nM
5nM
50nM
200nM
500nM
1000nM
### Endoggresomes/100 µm
C
DMSO
200 nM CCT020312
200 nM carboxy-CCT
Hoechst
NucGreen
100 µm
Percentage of NucGreen-positive cells
CCT020312
carboxy-CCT
D
PrPWT-mCh +/- rapamycin
PrPPG14-mCh +/- rapamycin
E
CDF
anterograde segmental velocities (µm/sec)
retrograde segmental velocities (µm/sec)
anterograde segmental run length (µm)
retrograde segmental run length (µm)
naxons: 32
PrPWT-mCh + DMSO
PrPPG14-mCh +DMSO
PrPPG14-mCh + CCT020312

### Data S2

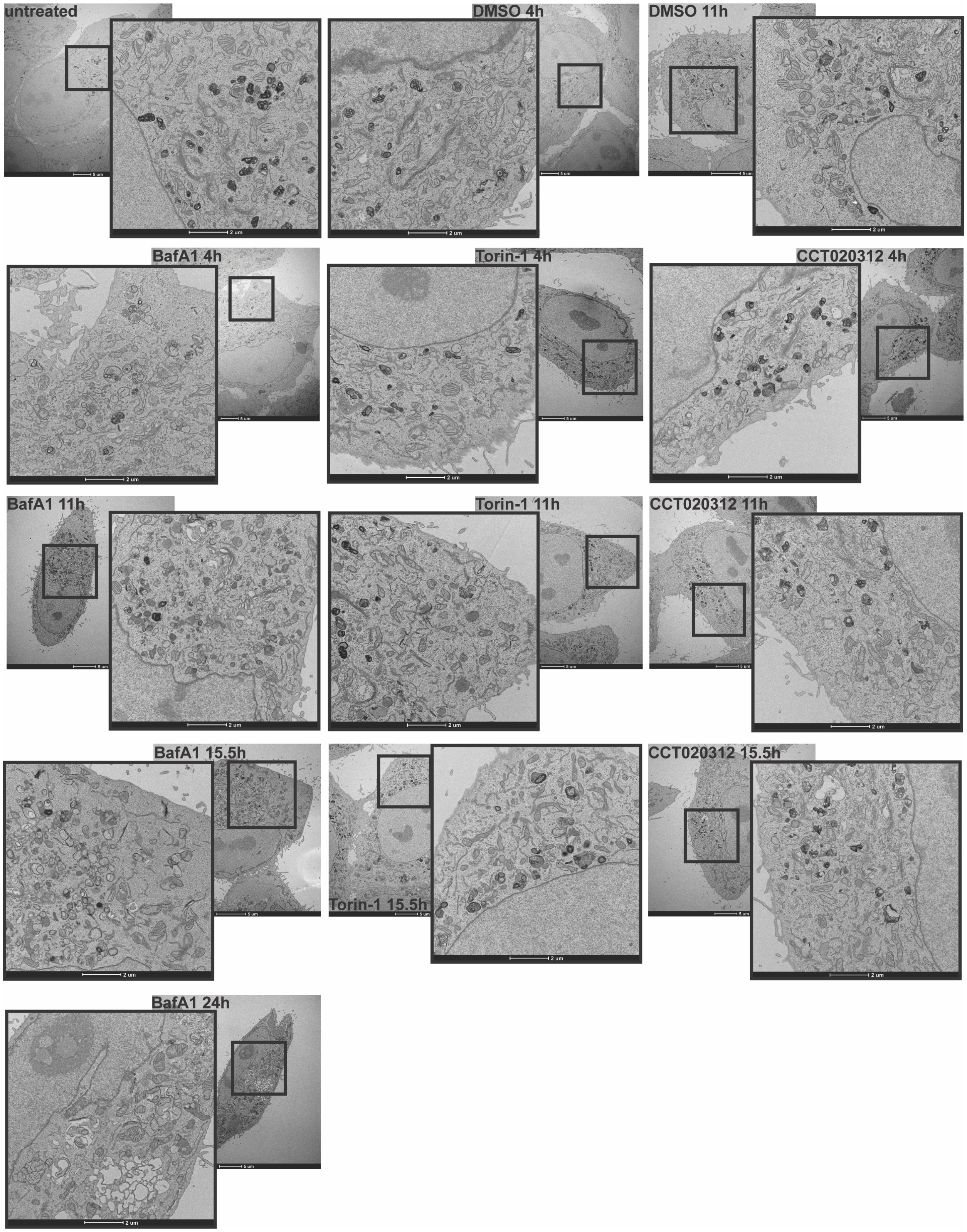
untreated
DMSO 4h
DMSO 11h
BafA1 4h
Torin-1 4h
CCT020312 4h
BafA1 11h
Torin-1 11h
CCT020312 11h
BafA1 15.5h
Torin-1 15.5h
CCT020312 15.5h
BafA1 24h
